# Conditions for maintaining and eroding pseudo-overdominance and its contribution to inbreeding depression

**DOI:** 10.1101/2021.12.16.473022

**Authors:** Diala Abu Awad, Donald Waller

## Abstract

Classical models that ignore linkage predict that deleterious recessive mutations should purge or fix within inbred populations, yet inbred populations often retain moderate to high segregating load. True overdominance could generate balancing selection strong enough to sustain inbreeding depression even within inbred populations, but this is considered rare. However, arrays of deleterious recessives linked in repulsion could generate appreciable pseudo-overdominance that would also sustain segregating load. We used simulations to explore how long pseudo-overdominant (POD) zones persist once created (e.g., by hybridization between populations fixed for alternative mildly deleterious mutations). Balanced haplotype loads, tight linkage, and moderate to strong cumulative selective effects all serve to maintain POD zones. Tight linkage is key, suggesting that such regions are most likely to arise and persist in low recombination regions (like inversions). Selection and drift unbalance the load, eventually eliminating POD zones, but this process is quite slow under strong pseudo-overdominance. Background selection accelerates the loss of weak POD zones but reinforces strong ones in inbred populations by disfavoring homozygotes. Models and empirical studies of POD dynamics within populations help us understand how POD zones may allow the load to persist, greatly affecting load dynamics and mating systems evolution.

## 1 Introduction

Inbreeding depression (δ) is defined as the lower fitness of inbred compared to outbred individuals (Darwin, 1876). It is now generally accepted that δ is mainly due to the expression of segregating deleterious recessive mutations (Charlesworth and Charlesworth, 1987; Crow, 1993; Bataillon and Kirkpatrick, 2000; Roze, 2015). As direct selection, background selection, genetic drift and inbreeding all act to reduce diversity at such loci, maintaining non-negligible levels of inbreeding depression is difficult to explain (Byers and Waller, 1999; Winn et al, 2011). Examples include inbred lines of *Zea mays* Kardos et al (2014); Larièpe et al (2012), *Arabidopsis* (Seymour et al, 2016), *Mimulus* (Brown and Kelly, 2020) and *C. elegans* (Chelo et al, 2019; Bernstein et al, 2019). Such observations led many to conclude that overdominant selection, *i.e*. a higher fitness of heterozygotes compared to either homozygote, was operating (Kimura and Ohta, 1971; Charlesworth and Charlesworth, 1987). But truly overdominant loci are rare, and most effects previously attributed to overdominance (such as heterosis and hybrid vigor) can be explained by simple dominance interactions (Crow, 1999a). Curiously, analyses of inbreeding depression often detect evidence of overdominance (see for example Baldwin and Schoen 2019). These apparent overdominant effects, however, probably reflect the effects of many deleterious recessive mutations linked in repulsion, a phenomenon termed pseudo-overdominance (hereafter POD, introduced by Ohta and Kimura 1969; reviewed by Waller 2021). We have known for half a century that a single strong overdominant locus can generate enough selection against homozygotes to persist even under complete self-fertilization (Kimura and Ohta, 1971). Could such strong effects also arise and persist via pseudo-overdominance?

Pseudo-overdominant selection will only emerge in genomic regions where many deleterious alleles are clustered together and often linked in repulsion, generating complementary haplotypes that express similar inbreeding loads as homozygotes. Genomic regions with reduced recombination, such as centromeric regions and chromosomal inversions, often maintain higher than expected heterozygosity. Centromeric regions in *Zea mays*, for example, maintain heterozygosity even after repeated generations of inbreeding (McMullen et al, 2009). This has also been found in 22 centromeric regions in the human genome (Gilbert et al, 2020). Kremling et al (2018) confirmed that many rare variants in maize express deleterious effects confirming that “even intensive artificial selection is insufficient to purge genetic load.” Brandenburg et al (2017) identified 6,978 genomic segments (≈ 9% of the genome) with unexpectedly high heterozygosity in land races of maize. These heterozygous segments contained more deleterious mutations than other parts of the genome, with several deeply conserved across multiple land races. Inversions, which halt recombination, also appear to accumulate lasting loads of deleterious mutations. Jay et al (2021) found that ancient inversions contribute greatly to heterosis in *Heliconius* butterflies. Kirkpatrick (2010) concluded that although the genetic basis for inversion overdominance has not yet been clearly determined, POD is plausible.

Pseudo-overdominance (POD) at many loci of small effect should mimic overdominant selection at a single locus, favouring heterozygosity for load within particular genomic regions. This could sustain inbreeding depression even in the face of purifying selection and drift. For POD to influence species evolution, it must exist for long enough and generate enough overdominant selection to leave a signature. Recombination, however, acts to break up such regions by unbalancing haplotype loads, allowing selection and drift to purge or fix their mutations. It is thus remarkable that polymorphic inversions expressing balancing selection date back to ancient hybridization events in *Heliconius* butterflies (Jay et al, 2021). Similarly, five ancient polymorphic zones predate the divergence of *Arabidopsis* from *Capsella* (approx. 8 million generations ago, Wu et al, 2017). These observations suggest that polymorphic regions may generate enough selection to sustain themselves for long periods of time. Could this selection derive from POD?

Several mechanisms might generate enough initial overdominance to create a POD zone including crosses between independently inbred lineages or sub-populations (generating high heterosis in the F1), a truly overdominant (e.g., self-incompatibility) locus, or chromosomal inversions where recombination is strongly suppressed, allowing mutations to accumulate. Here, we use simulations to study the evolutionary dynamics of POD zones generated initially by admixture between two populations fixed for different sets of deleterious mutations. In this scenario, high fitness emerges in the F1 where mutations fixed within each population are ‘masked’ as heterozygotes in hybrid offspring (Kim et al, 2018). We extend existing theory regarding the stable polymorphism that can exist at a single bi-allelic overdominant locus to examine the conditions necessary for POD to maintain two haplotypes containing many linked recessive deleterious mutations as heterozygotes. Because pseudo-overdominance depends on tight linkage among these loci, we expect that over time such zones will be vulnerable to being broken up by recombination. We therefore also explore how varying levels of linkage, dominance, selection and selfing rates affect POD zone stability and decay. Finally, we test how selection elsewhere in the genome affects the ability of POD zones to persist and the reciprocal effects of POD zones on load dynamics elsewhere in the genome.

## 2 Approaches

### 2.1 Load needed to generate a POD

Kimura and Ohta (1971) demonstrated that when the selective effects generating true overdominance are strong enough, a stable equilibrium can exist that perpetuates the two overdominant alleles indefinitely even within a fully self-fertilizing population. Consider a scenario in which two haplotypes, noted H1 and H2, occur within a diploid population self-fertilizing at rate *σ*. Each homozygote suffers a fitness reduction (*s*_1_ or *s*_2_) compared to the heterozygote fitness. In the case of true overdominance, Kimura and Ohta (1971) showed that a stable polymorphism will persist at an overdominant locus when:

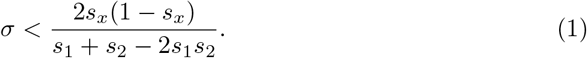

where *s_x_* = *min*(*s*_1_, *s*_2_) < 0.5. When both segregating homozygotes reduce fitness by at least half (*s*1, *s2* > 0.5), selection acts to maintain overdominance even as the selfing rate approaches one, as selection removes homozygotes faster than they are generated (Rocheleau and Lessard, 2000). For situations with stable polymorphism, setting *s*1 = *s*2 results in both alleles being maintained at a frequency of 0.5.

We use this threshold under true overdominance to estimate the number of load loci within pseudo-overdominant (POD) zone required to generate the necessary level of overdominance needed to maintain a stable equilibrium (see Eq. 1). For the sake of simplicity, we assume that each haplotype carries the same number *n_L_* of deleterious mutations all with the same coefficient of selection *s* and dominance *h*. We assume initial complete linkage, as it can then be broken by recombination, with loci evenly spaced, occurring at intervals of *ℓ* Morgans between alternating trans-mutations on opposing haplotypes (Fig. 1). As fitness effects are considered multiplicative across loci, an individual’s fitness is:

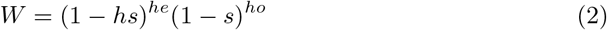

where *he* and *ho* are the number of heterozygous and homozygous mutations, respectively, carried by the individual. In the case of complete linkage homozygosity at these loci only occurs in individuals carrying two copies of the same haplotype (genotype *H*_1_*H*_1_ or *H*_2_*H*_2_). As both haplotypes carry the same number of mutations, the coefficient of selection acting against either homozygote (*s_H_* = *s*_1_, *s*_2_), relative to the fitness of the heterozygote *H*_1_*H*_2_ (*W_AA_/W_Aa_*) is:

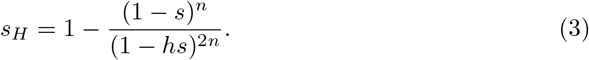

**Figure 1:**
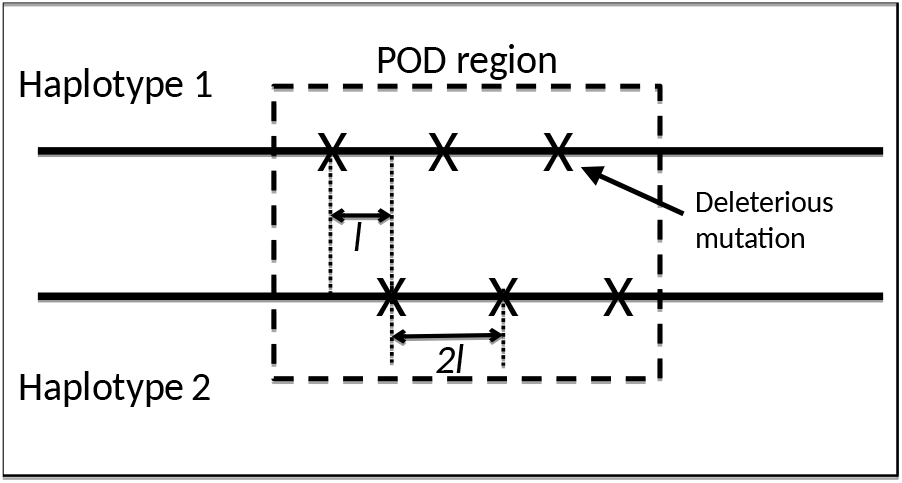
Genetic structure of the POD region (delimited by the dashed box). Deleterious mutations (represented by crosses) linked in cis occur at a distance 2*ℓ* M from each other along the same chromosome, alternating (at a distance *ℓ* M) with trans mutations on the opposite chromosome. Close, regular, and alternating spacing of recessive deleterious mutations along both haplotypes ensure linkage and pseudo-overdominance.

This expression allows us to determine the number of deleterious alleles per haplotype necessary to sustain enough overdominance to preserve both haplotypes via stable balancing selection (see Supp. File 1):

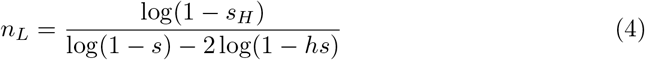

As expected, the number of loci required to obtain a strength of selection against homozygotes *s_H_* decreases for higher values of *s* and *h*. For *s* = 0.01 and *h* = 0.2, *n_L_* = 115 for *s_H_* to be at least 0.5, which should sustain POD selection indefinitely (Supp. File 1, Fig. S1).

### 2.2 Inbreeding depression

Inbreeding depression δ is a population specific variable, reflecting the number of heterozygotes maintained in a population. The general equation used to estimate inbreeding depression is:

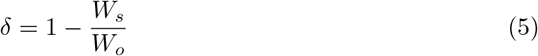

where *W_s_* is the fitness of selfed offspring and *W_o_* that of outcrossed offspring (Charlesworth and Charlesworth, 1987). If there is a POD zone, we can consider that there are two potential forms of selection contributing to inbreeding depression: 1) selection against deleterious mutations that are scattered throughout the genome (noted *δ_s_*) and 2) over-dominant selection generated by POD zones (noted *δ_od_*). If we assume that selection against deleterious mutations elsewhere in the genome and overdominant selection do not interfere with one another (*i.e*. no associative overdominance or effects of background selection) and fitness effects remain multiplicative (see for example Kirkpatrick and Jarne 2000, the upper limit of the expected level of inbreeding depression will be:

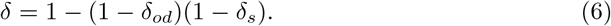

When mutations are deleterious, and accounting for drift, *δ_s_* depends on the haploid mutation rate *U*, the coefficient of selection *s* and the dominance of mutations *h* (see equation 3 from Bataillon and Kirkpatrick 2000):

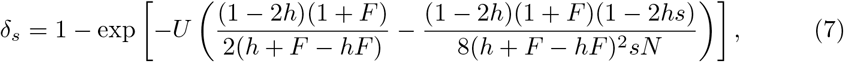

where *F* = *σ*/(2 – *σ*) is the equilibrium inbreeding coefficient (expected deviation from Hardy-Weinberg equilibrium of genotype frequencies). Though this expression for F remains true for weak overdominance (Glémin, 2021), when there is strong overdominance, the inbreeding coefficient depends on the coefficients of selection and allelic frequencies (Appendix A4 from Kimura and Ohta, 1971). In our case with symmetrical selection against homozygotes, this term is given as:

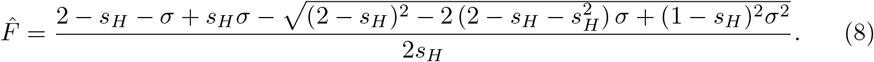

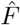 will tend to zero with increasing *s_H_* (see Fig. A1 in Supp. File 1). Selfing populations subject to strong overdominant selection thus tend to behave like outcrossing ones as low fitness homozygotes are eliminated. In the presence of POD selection, we set *F* in Eq. 7 to 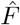.

At equilibrium, the contribution of POD to inbreeding depression *δ_od_* can, for symmetrical overdominance, be written as:

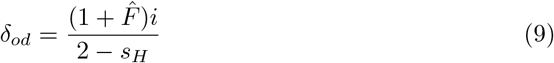

where 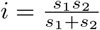, which simplifies to 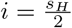 when *s*_1_ = *s*_2_ = *s_H_* - see Eq. A2 from Supp. File 1 and Kimura and Ohta (1971). We provide the general expressions for 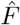 and *δ_od_* in Supp. File 1 (see Eq. A3).

As previously shown, *δ_od_* increases with the selfing rate *σ* for strong overdominant selection and *δ_s_* decreases with *σ* (Charlesworth and Charlesworth, 1987, 1990). It is therefore possible to have similar *δ* (given in Eq. 6) in outcrossers and selfers, depending on the rates of background mutation U and the strength of POD selection (*i.e*. the value of *s_H_*).

### 2.3 Recombination and POD’s

Thus far, we have assumed complete linkage in order to apply one-locus overdominance theory to infer the strength of selection against homozygotes necessary to sustain a stable equilibrium. However, some recombination will occur, allowing the strong linkage disequilibrium among loci within a POD to erode over time. In order to examine the effect of recombination on the stability of POD, we propose a system of Ordinary Difference Equations (ODEs) representing the change in frequencies of the two initial haplotypes (Δ_*P*_1__ and Δ_*p*_2__) and that of a newly introduced recombinant haplotype (Δ_*P*_c__):

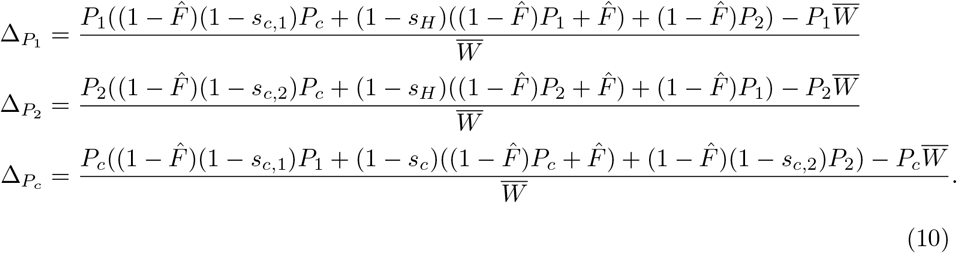

The mean fitness of the population 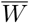 is the sum of the expected genotypic frequencies after selection (see Supp. File 2, Eq. (A4)), and *s_c_*, *s*_*c*,1_ and *s*_*c*,2_ are the coefficients of selection associated respectively with haplotypes *H_c_H_c_*, *H_c_H*_1_ and *H_c_H*_2_. We resolve this system of equations to determine the conditions necessary for a recombinant haplotype *H_c_* to increase in frequency (Δ*_P_c__* > 0).

## 3 Simulations

So as to confirm expectations from the analytical model given above and explore the dynamics of POD selection, we develop an individual-based simulation program in C++, uploaded to Zenodo.org (Abu Awad and Waller, 2022). We consider a scenario where POD selection arises after an admixture event between two initially isolated populations fixed for different mutations within the same genomic region (a “proto-POD” zone). Each population is made up of *N* sexual diploid individuals, self-fertilizing at a fixed rate, σ. Each individual is represented by two vectors, each carrying the positions (between 0 and 1) of deleterious mutations along a single chromosome with map length *R* Morgans.

Recombination occurs uniformly throughout the genome. Mutations within and outside of the POD zone have a fixed effect, with respective coefficients of selection, *s* and *s_d_*, and dominances, *h* and *h_d_*. Individual fitness is calculated as shown in Eq. 2. New mutations are sampled from a Poisson distribution with parameter U, the haploid mutation rate and their positions are uniformly distributed along the genome (infinite-locus model). Generations are discrete (no overlap) and consist of three phases: *i*) introducing new mutations, *ii*) selection, and *iii*) recombination and gamete production.

### 3.1 POD zone architecture and initiation

Two types of simulation are run, one with an arbitrary ideal haplotype structure expected to favour POD persistence and one with a more realistic distribution of mutations within the POD zone. The former consists of constructing two perfectly complementary haplotypes, *H*_1_ and *H*_2_. Cis-mutations occur at regular intervals (every 2*ℓ* M) along each haplotype and mutations are staggered, spreading the load evenly through the POD and ensuring pseudo-overdominance (Fig. 1). The expected number of recombination events occurring between two trans-mutations is then *ℓ*. The second type of POD zone architecture is one with randomly placed mutations in a predefined genomic region, their positions sampled from a uniform distribution, while ensuring that a locus with the same position is not sampled for both haplotypes. In both cases the center of the POD zone is kept constant for both haplotypes and the size of the POD zone is 2*ℓn_L_* M, with *n_L_* potentially different for each haplotype. The POD zone is arbitrarily positioned around the center of the genome, its exact center at position 0.5 along the chromosome.

After a burn-in period of 4 000 generations, allowing the two source populations (each fixed for a given haplotype in the proto-POD zone) to reach mutation-selection-drift equilibrium, a new population of size *N* is created by randomly sampling individuals from both populations. We arbitrarily consider that each source population contributes 50% of individuals to the new population. The new population is then allowed to evolve for a further 4000 generations. Samples of 100 individuals are taken every 10 generations to estimate inbreeding depression, which we compare to the theoretical expectations presented above (Eqs. 7, 9 and 6). We also use these samples to estimate heterozygosity within and outside the POD zone (POD *H_e_* and genome *H_e_*, respectively) as:

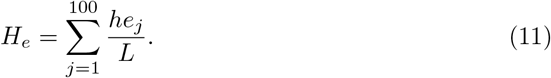

where *he_i_* is the number of heterozygous mutations carried by individual *j* (out of a sample of 100) and *L* is the total number of segregating sites in the genomic region of interest. A decrease of *H_e_* with time signals the erosion of the POD zone, either through loss or fixations of mutations.

Unless stated otherwise, all variable plotted are values obtained 4000 generations after the hybridisation event. Figures are made using the ggplot2 package (v3.3.6, Wickham 2016), with, in most cases, lines generated using the geom_smooth option. When this gave results that were too divergent compared to plotting the mean, the mean was used.

### 3.2 Simulations run

Simulations are run for population size *N* = 100, 1000 and 5000 and for selfing rates σ between 0 and 0.95. The haploid background mutation U is set to 0, 0.1 and 0.5, with new mutations outside the POD zone having a fixed coefficient of selection (*s_d_* = 0.01) and dominance (*h_d_* = 0.2 or 0.5). We explore the effect of genome map length *R*, choosing *R* =1 and 10 Morgans for tight and loose linkage respectively, and we examine different strengths of linkage between loci in the POD zone, with *ℓ* = 10^-4^, 10^-5^ and 10^-6^. We consider both weak and strong selection against homozygotes, setting *s_H_* to *s_H_* = 0.14, 0.26 and 0.45. These correspond to stable (polymorphic) overdominant selection when *σ* = 0, 0.5 or even (with a narrow range of stability) 0.95 (Fig. A2, dotted lines). To determine the effects of POD selection on heterozygosity elsewhere in the genome, we also run simulations where all alleles within the initial POD zone are neutral for all parameter sets mentioned above (achieved by setting *s* and *h* = 0 within the POD). We run 100 repetitions for each parameter set.

## 4 Results

### 4.1 POD persistence and degradation

We first examine how recombination, the strength of selection against linked load loci, and their arrangement within the POD zone, influence POD persistence.

#### 4.1.1 Recombination and POD degradation

Under the assumption that recombination within the POD block is rare (reflecting tight linkage), any new haplotype *H_c_* will be generated by a single recombination event. This is reflected in the ODEs introduced in Eq. (10) which compute changes in frequency of the two initial haplotypes (*H*_1_ and *H*_2_) and a recombinant (*H_c_*). For simplicity, we initially assume an ideal case where mutations are arranged alternately within the POD zone (see Fig 1). Positions of deleterious alleles in *H*_1_*H*_2_ heterozygotes alternate in trans relative to flanking mutations on the same chromosome (Fig. 1). Each haplotype carries *n_L_* deleterious mutations. Consider two cases: 1) the recombinant haplotype *H_c_* (and its complement) each carry *n_L_* deleterious mutations; 2) *H_c_* carries *n_L_* – 1 mutations because recombination has cleaved one from one end of the POD zone.

Given arbitrary values of *s*_*c*_, *s*_*c*,1_ and *s*_*c*,2_ (the coefficients of selection against *H_c_H_c_*, *H_c_H*_1_ and *H_c_H*_2_ genotypes, respectively), the only possible equilibria involve fixing one of the three haplotypes or maintaining only two of them. Hence any rare haplotype, *H_c_*, should either be lost, go to fixation, or replace one of the initial haplotypes (co-existing with the other). For *H_c_* to increase in frequency, Δ_*P_c_*_ (Eq. (10)) must be positive when it enters the population (or it would be eliminated). Assuming the frequency of a recombinant *P_c_* is of order *ϵ* (*ϵ* being very small), the expression for Δ_*P_c_*_ for the leading order of *P_c_* (noted 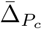) can be derived. In a population at equilibrium with *P*_1_ = *P*_2_ = (1 – *ϵ*)/2 and setting *s*_1_ = *s*_2_ = *s_H_*:

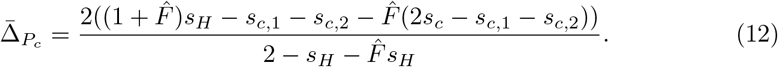

The denominator of this expression is always greater than 0 for *s_H_* < 1. To understand the behavior of 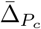, we simplify the above equation by setting 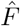 to 0 (no self-fertilisation or very strong overdominant selection with *s_H_* ≈ 1, see Supp Fig. A1). In this case Eq. 12 simplifies to 2(*s_H_* – *s*_*c*,1_ – *s*_*c*,2_)/(2 – *s_H_*). If no mutations have been cleaved off by recombination (*i.e H_c_* carries *n_L_* mutations), the numerator 2(*s_H_* – *s*_*c*,1_ – *s*_*c*,2_) ≤ 0 (see Eq. B1 in Supp. File 2 for expressions of *s*_*c*,1_ and *s*_*c*,2_) making 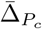 negative (Fig. B2 in Supp. File 2). Hence Hc haplotypes will be selected against. This is because recombinant *H_c_* haplotypes will share mutations with both the initial *H*_1_ and *H*_2_ haplotypes and a proportion of loci in *H_c_H*_1_ and *H_c_H*_2_ genotypes will inevitably be homozygous, resulting in a lower fitness of these genotypes compared to *H*_1_*H*_2_ heterozygotes. In this case neither the homozygous nor heterozygous genotypes with a recombinant haplotype present a selective advantage. If instead H_c_ carries *n_L_* – 1 mutations, the resulting coefficients of selection (Eq. B2, Supp. File 2) lead to a positive 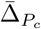 (the numerator in this case can be positive). The larger 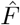 (or the selfing rate σ) the more positive the resulting 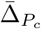.

This result leads us to predict that if a POD is initially stable, its eventual loss will usually occur gradually as recombination events near the distal ends of the POD cleave off mutations creating haplotypes with improved relative fitness. The reduced zones of stable equilibria for *s_c_* = *s_H_* in selfing populations (Fig. A2, in Supp. File 1) means that selection will more easily act to destabilise the POD zone by eroding mutations. This should fix one of the original haplotypes or a recombinant with the strength of selection affecting the rate at which this occurs.

Using simulations, we confirm results from single locus overdominance that stronger selection is more likely to result in stable polymorphism even for high selfing rates (Supp. Fig. S2). Drift and selection can both act to erode POD (shown by the rate of decrease of heterozygosity in Supp. Fig. S2). Strong drift renders selection neutral when *N_e_s_H_* << 1, accelerating the loss of supposedly stable POD selection (*N* = 100 in Supp Fig. S2). Increasing the efficacy of selection will also favour the loss of POD selection, but unlike for strong drift, this is due to a more efficient purging (and higher effective recombination rate) of loci contributing to POD selection (*N* = 5000 in Supp Fig. S2).

As the differences between population sizes are quantitative, and *s_H_* is a good predictor of mid/long-term stability of POD zones, in the following, we examine simulations only for *N* = 1000, for which both drift and selection act on POD stability, and *s_H_* = 0.45, for which overdominant selection is stable for all self-fertilisation rates simulated.

#### 4.1.2 Effect of the strength of selection against individual loci

As mutations are progressively lost from POD zones, recombinants can go to fixation. This will eventually destabilize the POD zone. We next assess how varying the coefficients of selection *s* and dominance *h* against individual loci affects POD persistence. For a fixed value of selection against homozygotes, *s_H_*, varying *s, h* and *n_L_* (obtained using Eq. (4)), we calculate the expected increase in frequency a recombinant haplotype 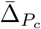 using Eq. (12). If no mutation is lost (*H_c_* also carries *n_L_* mutations), 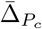 remains negative except under high rates of self-fertilisation when they can be positive (though close to 0). However, a mutation lost through recombination generates a positive 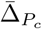 that increases with increasing strengths of selection and dominance of the mutations for all rates of self-fertilisation (Figs. 2 a and b for *s_H_* = 0.45). We confirm this prediction via simulations. These show that most losses of diversity (fixation or loss of mutations) occur at the ends of the POD zone (Figs.2c and d for selfing rate *σ* = 0.95). Losses of diversity within the POD zone intensify as *s* and *h* increase.

**Figure 2:**
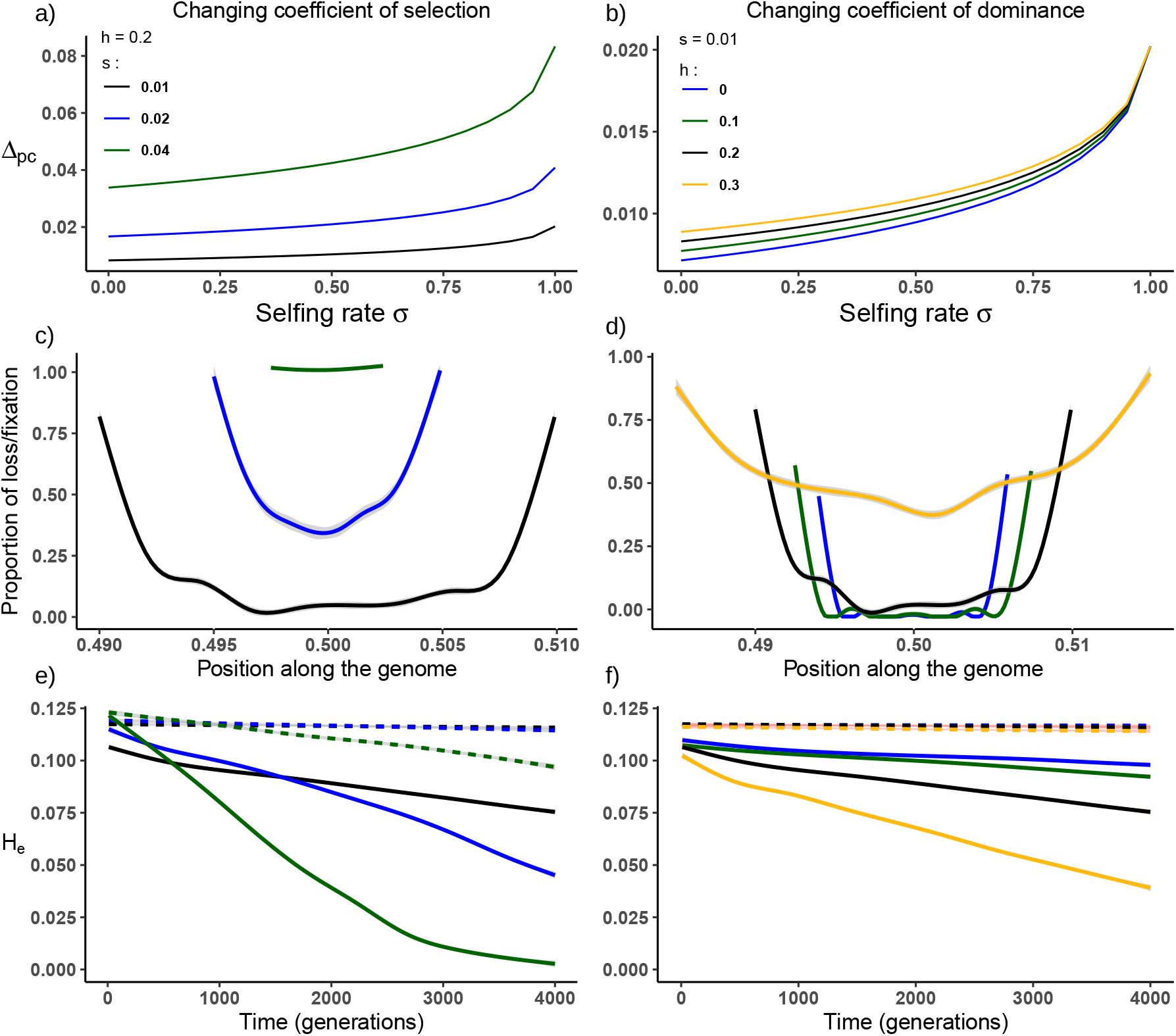
Effects of levels of selection and dominance on selection dynamics within a POD zone. Left panels show the effects of varying the coefficient of selection at a load locus *s* (*s* = 0.01, 0.02 and 0.04, corresponding to *n_L_* = 100, 50 and 25 loci). Dominance is fixed at *h* = 0.2 and *s_H_* = 0.45. Right panels show the effects of varying dominance (*h* = 0,0.1, 0.2 and 0.5 with *n_L_* = 60, 75, 100 and 150) with selection fixed at *s* = 0.01. Panels a) and b) show theoretical rates of increase in frequency for a recombinant haplotype that loses a mutation from one end. Panels c) and d) show observed frequencies of fixation/loss along the POD zone at generation 4000 (x values represent the position of the loci along the chromosome). The selfing rate *σ* = 0.95 and linkage *ℓ* = 10^-4^M. Panels e) and f) show losses in heterozygosity (*H_e_*) over time in populations with a high selfing rate (*σ* = 0.95) and either loose linkage (*ℓ* = 10^-4^M, solid lines) or tight linkage (*ℓ* = 10^-5^M, dashed lines). Population size N = 1000.

Stronger selection against individual mutations sustains heterozygosity more effectively as fewer mutations suffice to generate the same amount of balancing selection. However, the loss of a stronger mutation as a result of recombination will more likely unbalance and destabilise the POD zone. This accelerates the fixation or loss of mutations (Fig.2c). Increasing the dominance of load loci has similar effects as increasing *s* but requires more mutations to reach the same *s_H_* (*i.e*. *n_L_* = 60 and 150 for *h* = 0 and 0.3 respectively, Fig. 2f). This is because increased dominance increases the relative fitness of both the fitter homozygote (*i.e*. the haplotype with one less mutation due to recombination) and the heterozygote, increasing the overall fitness advantage of losing a mutation. The same patterns are observed in outcrossing populations to a lesser extent (Supp. Fig. S3). Increased linkage within the POD zone reduces the rate at which these higher fitness recombinants occur, slowing this process (dashed lines, Figs. 2e and f; see Supp. Fig. S4 for patterns of mutation loss within the POD zone).

#### 4.1.3 POD region architecture

So far, we have considered only an ideal genetic architecture that favours maintaining POD, namely homozygotes of both haplotypes having identical fitness disadvantages relative to the heterozygote and equally spaced cis and trans mutations within the POD zone. We now relax these assumptions by considering initial haplotypes carrying different numbers of mutations, *n_L_*, within the POD region (while maintaining equal spacing) and then by placing randomly spaced mutations within the POD zone.

To unbalance the segregating homozygotes, consider alternative POD zone haplotypes with *n_L_* = 80, 100, *or120* mutations paired with a haplotype *H*_1_ with *n_L_* = 100 mutations (denoted by relative lengths of 0.8 1 and 1.2 respectively in Figs. 3a and c). These generate substantial fitness differentials with relative selection coefficients against homozygotes *s*_1_ = 0.47 and *s*_2_ = 0.35 (blue lines), *s*_1_ = *s*_2_ = 0.45 (black lines), or *s*_1_ = 0.43 and *s*_2_ = 0.53 (green lines). In outcrossing populations, selection trims down longer, more loaded haplotypes as recombination makes variants available. This shrinks more loaded haplotypes to sizes close to the smaller haplotype (Fig. 3a, solid lines). Overdominant selection, however, sustains the core POD region’s heterozygosity, *H_e_*(Fig. 3b, solid lines). Self-fertilising populations, in contrast, show less POD zone stability under asymmetric selection despite the fact that populations with balanced loads showed only slight observed losses or fixations of mutations (dashed black lines in Figs. 3a and c). When the alternative haplotype has less load (a relative size of 0.8), it quickly goes to fixation (dashed blue lines in Figs. 3a and c). This result matches the theoretical expectation that no overdominant polymorphism can be maintained with these coefficients of selection against homozygotes when the selfing rate is 0.95 (see Fig.A2 in the Supp. File 1). When the total load of the second haplotype increases to a relative size of 1.2, the POD zone is more commonly sustained as mutations are trimmed off the ends of the POD zone (Fig. 3a, c). This difference in behavior reflects the need for segregating load to exceed a threshold to sustain a POD zone. As for outcrossing, most mutations of the larger haplotype will be trimmed off the edges, but there is some fixation and/or loss of mutations along the whole POD region (dashed green line in Fig. 3a), lowering the mean observed *H_e_* (dashed green line in Fig. 3c). This is most probably due to a larger range of recombinants having a higher selective advantage, provided that they trim the larger haplotype and thus help destabilize POD selection.

**Figure 3:**
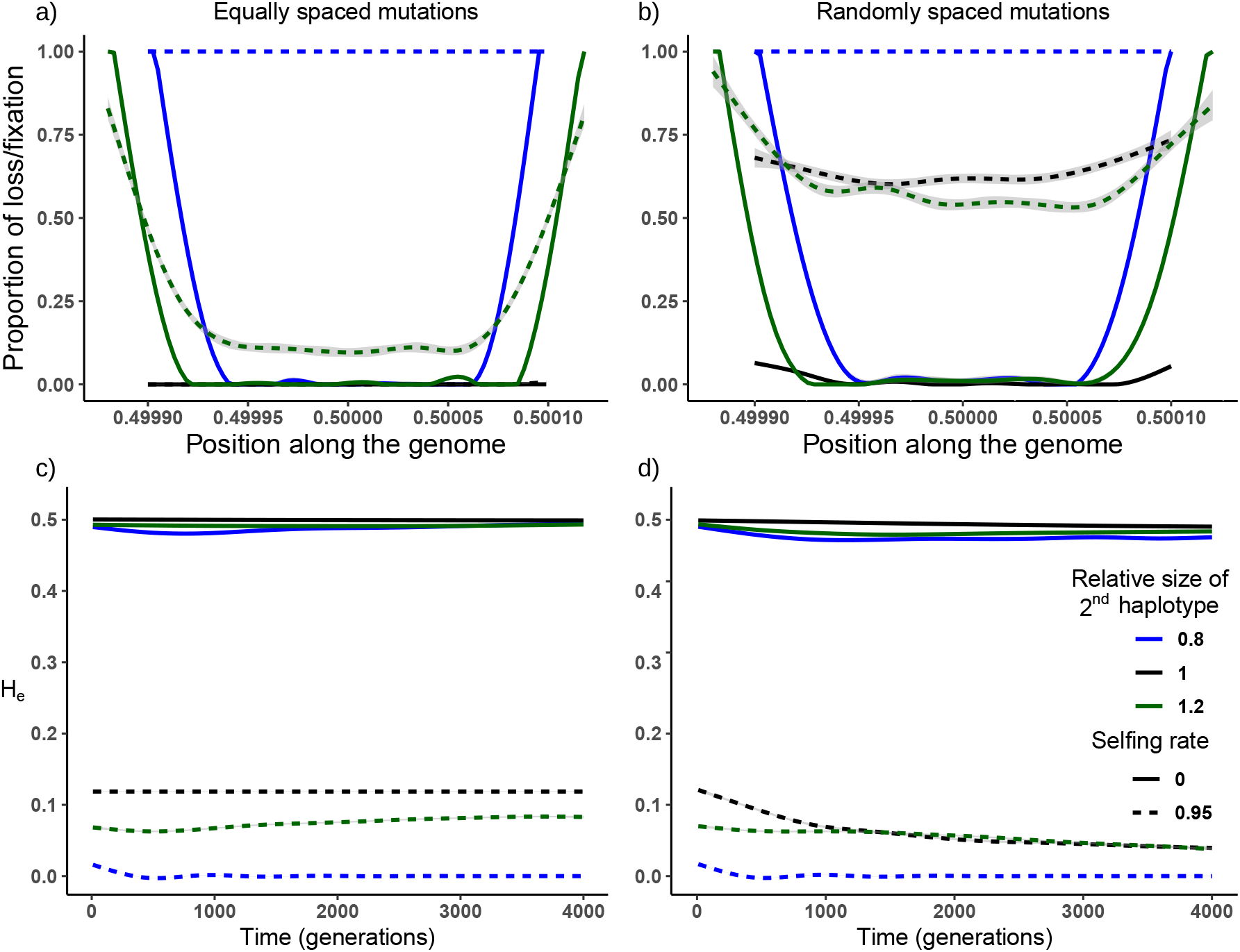
Effects of relaxing the assumptions of symmetric overdominance and evenly spaced mutations. Upper panels show locations within the POD zone where load mutations are most likely to be lost (a, b) and how this depends on whether mutations are evenly spaced (a) or randomly distributed (b). Results are shown for both symmetric (black) and asymmetric (green and blue) loads. Outcomes under both outcrossing and high selfing (solid vs. dotted lines) are shown. Note erosion of mutations via recombination and selection at both ends of the POD zone. Lower panels show overall stability of the POD zone (shown as heterozygosity, *He*) over time. As in the upper panels, graphs show results for both symmetric (black) and asymmetric (green and blue) loads and for evenly and randomly placed mutations (panel c vs. d). The coefficients of selection and dominance are *s* = 0.01 and *h* = 0.2 respectively, linkage within the POD zone is *ℓ* = 10^-6^ and population size *N* = 1000.

When the mutations are not in an ideal configuration, but randomly positioned throughout the designated POD zone, stability of the POD zone is barely affected in outcrossing populations (solid lines in Figs. 3b and d), even when the haplotypes are initially uneven. Selfing populations, however, require stronger linkage to retain the POD zone (compare dashed lines in Fig. 3 for *ℓ* = 10^-6^ M to Fig. S5 for *ℓ* = 10^-5^). Despite more frequent fixations/losses of mutations, some heterozygosity nonetheless persists for approximately 1000 generations even with lower linkage (Supp. Fig. S5).

### 4.2 Background mutations

Mutations introduced elsewhere in the genome influence POD selection dynamics and persistence and vice versa as POD’s affect purifying selection across the genome. In general, when a POD zone is stable, background mutations will not destabilise it. Background selection does, however, affect heterozygosity within and outside the POD zone. Let us compare heterozygosity within the POD zone in simulations with background mutations to simulations lacking it (i.e. *U* > 0 vs. *U* = 0; Fig. 4a). Interestingly, in self-fertilising populations, *H_e_* within the POD zone rises when background selection occurs elsewhere in the genome. These effects increase when mutation rates rise (green *vs*. blue lines, *U* = 0.5 and 0.1 respectively) and linkage increases (full *vs*. dashed lines reflecting map lengths of *R* = 1 and 10 Morgans respectively).

**Figure 4:**
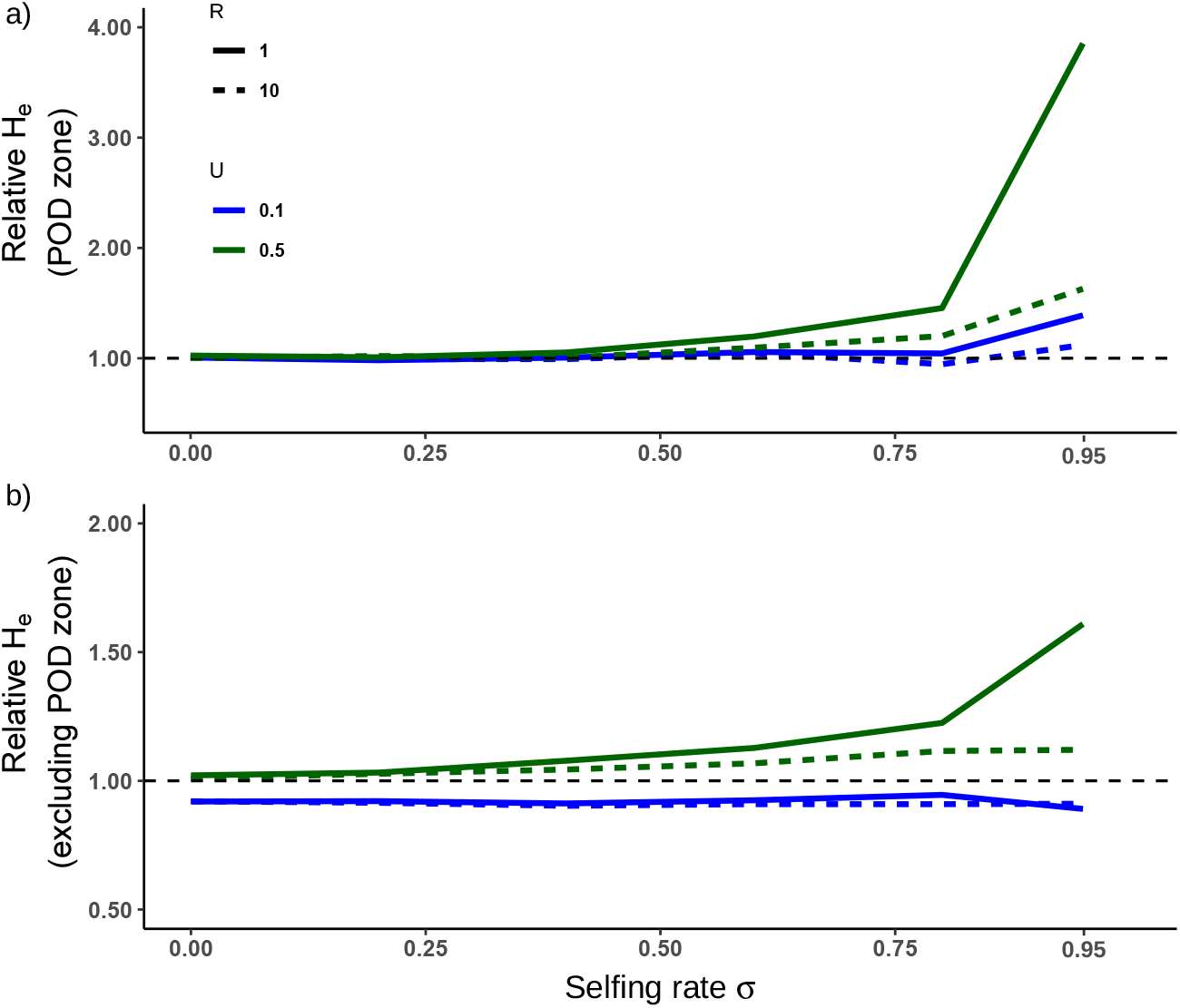
Background mutations affect POD selection and vice versa. Graph (a) shows heterozygosity, *H_e_*, within the POD zone with background mutations relative to *H_e_* in the absence of background mutations and graph (b) *H_e_* elsewhere in the genome with a POD zone relative to without, both as a function of the selfing rate. Populations are subject to different background mutation rates (*U*) and shorter and longer map lengths (*R* in Morgans). These simulations use 100 POD load loci (*n_L_* = 100) and a map length of *ℓ* = 10^-6^ Morgans. Mutations within the POD zone are randomly placed. Selection coefficients in- and outside the POD zone (*s* and *s_d_* respectively) are 0.01 with dominances *h* and *h_d_* = 0.2.

Similarly, the presence of a stable POD zone affects the heterozygosity of deleterious mutations observed elsewhere in the genome. When mutation rates are low (*U* = 0.1), POD selection slightly decreases the mutational heterozygosity elsewhere in the genome (blue lines Fig. 4b). Conversely, a higher genomic mutation rate (*U* = 0.5, green lines) results in increased heterozygosity, especially in highly selfing populations with small map lengths (implying tight linkage - solid green line in Fig. 4b). Effects of POD selection on effective population size are complex but in most cases, POD selection tends to decrease *N_e_* (Supp. Fig. S6).

To confirm that these effects derive from overdominance rather than some other effect of background selection, we simulated effects of co-dominant background mutations (*h_d_* = 0.5). Because such mutations are expressed in heterozygotes and thus easily removed by selection, they generate few associations with other loci. Co-dominant background mutations have little effect on within-POD zone heterozygosity in contrast to simulations with more recessive mutations (*h_d_* = 0.2). This is true even within selfing populations (Supp. Fig. S7a). This confirms that it is associative overdominance between the POD zone and other load loci that increases heterozygosity (Supp. Fig. S7b). Varying rates of background mutation and POD zone length also have complex effects on effective population size *N_e_* (Supp. Fig. S7c).

### 4.3 Inbreeding depression

As expected, the overdominance generated in a POD zone increases the inbreeding depression, *δ*, populations express (Supp. Fig. S8). Observed *δ* in outcrossing populations can be predicted using Eq. (6), which accounts for overdominant selection and unlinked deleterious mutations. In selfing populations variable erosion of the POD zone and POD selection dynamics generate bimodal distributions of *δ* (see Supp. Fig. S9 for clearer representations). Some simulations generate values of *δ* close to those predicted by Eq. (6) (dashed lines in Fig. 5) while others generate values predicted when selection acts only against the unlinked recessive deleterious mutations (Eq. (7), dotted lines in Fig. 5). This may reflect loss of the POD zone. Genomes with smaller map lengths (*e.g*., *R* =1 Morgans) generally increase the observed *δ*, especially in selfing populations (see Supp. Figs. S8 and S10).

**Figure 5:**
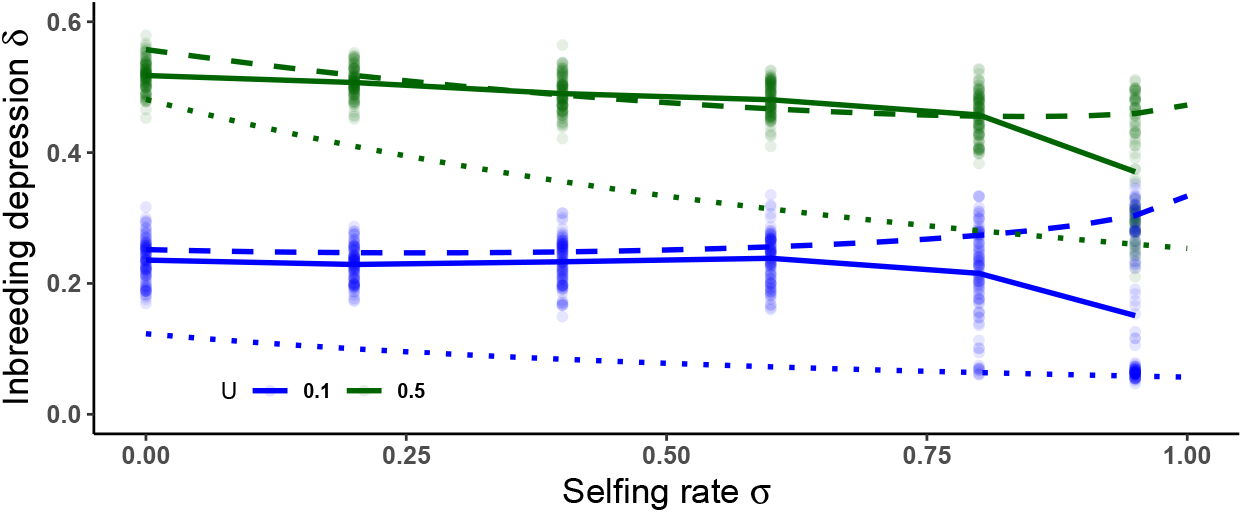
Inbreeding depression *δ* as a function of the selfing rate for different values of the haploid mutation rate, *U*. Solid lines show means of the simulations run. Dotted lines show the inbreeding depression expected in the absence of overdominance (Eq. (7)) while dashed lines show increases in delta expected with overdominant selection over all selfing rates (Eq. (6)). Other parameter values are *n_L_* = 100, *ℓ* = 10^-6^ Morgans and randomly placed mutations in the POD zone. Selection coefficients in- and outside the POD zone, *s* and *s_d_* are set to 0.01 with dominances *h* and *h_d_* = 0.2. The total map length (setting the recombination rate) is *R* = 10 Morgans.

## 5 Discussion

Given that purging, drift, and background selection all reduce segregating variation and thus inbreeding depression, we face the question of what force perpetuates these, even within small and inbred populations. Waller (2021) emphasized this enigma and reviewed mechanisms that might account for it. Selective interference among loci might act to slow or block purging (Lande and Schemske, 1985a; Winn et al, 2011). Recurrent mutations might also replenish the load fast enough to regenerate δ (Fisher, 1930; Charlesworth, 2018). A third possibility is that clusters of recessive mutations linked in repulsion emerge, creating enough balancing selection via pseudo-overdominance (POD) to counter purging and drift, sustaining selection for outcrossing or mixed mating systems (Waller, 2021). Our goals here were to explore the dynamic stability of POD zones (initially ignoring how they arise) using both classical one-locus overdominant theory (Kimura and Ohta, 1971) and simulations. We found that strong and balanced POD zones can persist for hundreds to many thousands of generations.

Whether POD zones are fragile or robust depends critically on several genetic parameters. These include the number and severity of deleterious mutations, their proximity and cis-/trans-positions, and their levels of dominance/recessivity (Figs. 2 and S3). Strong and balanced selection plus tight linkage allow POD zones to persist as these conditions enhance the associations (linkage disequilibria) that generate POD effects. Recombination dissolves these associations, allowing purifying selection and drift to disrupt POD zones, purging and fixing mutations. Mutations erode from either end of the POD zone or the load becomes unbalanced enough to fix one haplotype. The importance of linkage and small mutational effects are evident in the radically enhanced purging seen in models that ignore linkage and assume major mutational effects (Lande and Schemske, 1985b). We also found that new recessive mutations that occur elsewhere in the genome generate associations with load alleles within POD zones that enhance POD zone heterozygosity and persistence (Fig. 4). Such mutations add to the segregating load, increasing heterozygote advantage. Because levels of heterozygosity are correlated across the genome in partially inbred populations (identity disequilibrium), the background selection generated by mutations outside the POD zone tend to reinforce the balancing selection favoring heterozygotes in the POD zone. POD zones also exert reciprocal effects, enhancing the heterozygosity of mutations occurring elsewhere in the genome when mutation rates are moderate (U=0.5, Fig. 4b). This effect was amplified within selfing populations, presumably reflecting how selection against POD zone homozygotes favors heterozygosity across the genome when more identity disequilibrium occurs. These effects would be further enhanced if mutations were to have varying dominance effects, a scenario which we did not consider here. However, recent work has shown that POD selection can be generated in a single population by the clustering of mutations in repulsion, even without heterogenous recombination rates along the chromosome (Sianta et al, 2021). These results coupled with ours lead us to hypothesize that any genomic region displaying reduced recombination could provide a haven for POD zones to emerge and persist.

### 5.1 How do POD zones originate?

Many empirical observations could be explained by the existence of POD zones (see Introduction and Waller 2021). Whether POD zones that are conserved across populations exist in sufficient number and strength to affect evolutionary dynamics hinges on the relative rates at which they are created and destroyed. We focused on POD zone erosion and loss, not how they arise. As our results show, a requirement for POD stability is strong linkage within a given genomic region in which mutations can accumulate through the actions of selection and genetic drift. Inversions and centromeric regions with restricted recombination provide preconditions favoring POD zone emergence, as do genomic regions neighbouring loci currently or previously under overdominant selection, where recombination is suppressed. Examples where this has been observed include self-incompatibility loci (Takebayashi, 2003; Igic et al, 2008; Mable, 2008), MHC loci (Garrigan and Hedrick, 2003; Gemmell and Slate, 2006), and loci with balanced polymorphisms generated by ecological selection (van Oosterhout et al, 2000; Jay et al, 2021). In such regions, mutations of small effect become effectively neutral when the product of the effective population size and the selection coefficient *N_e_s* << 1 (Crow and Kimura, 1970; Hedrick et al, 2016)). These will drift in frequency and often fix increasing the “drift load” to the point where it may compromise population viability (Whitlock et al, 2000; Charlesworth, 2018). Selection against strongly deleterious mutations will accentuate fixation of milder mutations linked in repulsion via “background selection” (Charlesworth et al, 1997; Zhao and Charlesworth, 2016). Pairwise and higher associations (linkage disequilibria) also increase within small and inbred populations even among alleles at unlinked loci limiting selection (Hill and Robertson, 1966; Sved, 1971; Ohta and Cockerham, 1974; Lewontin, 1974).

The scenario we suggested that might create POD zones involved drift fixing alternative sets of recessive deleterious mutations among isolated populations. When such populations hybridize, their F1 progeny experience high heterosis reflecting the cumulative effects of POD across the whole genome (Crow, 1999b). Under free recombination, this heterosis is expected to erode by 50% in the F2 and each subsequent generation as recombination dissipates the associations generating the POD (Harkness et al, 2019) (ignoring the presence of epistatic Dobzhansky-Muller incompatibilities -(Ehiobu et al, 1989). However, where clumps of mutations occur within short genomic regions (or in low recombination zones), POD zones may be spawned. Inter-population crosses often reveal high heterosis (Willi et al, 2013; Spigler et al, 2017) as do crosses between low-fitness inbred lines in plant and animal breeding programs. Theory suggests that any incipient POD zone generating heterozygous progeny at least twice as fit as homozygous progeny will allow that POD zone to persist even in highly selfing populations. Dramatic examples of “hybrid vigor” in F1 crosses include cases where progeny have up to 35 times the fitness of parental lineages (Tallmon et al, 2004; Hedrick and Garcia-Dorado, 2016) easily satisfying this condition.

Proto-POD zones may be fragile. Our models show that recombination and selection eliminate proto-POD zones with weak, unbalanced, or loosely linked loads. However, in some regions, cumulative selective effects from localized mutations may be large and balanced enough to allow a persistent POD zone to emerge. Such zones eliminate many homozygous progeny, reducing effective rates of inbreeding (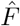, Eq. 8). This, in turn, reduces rates at which deleterious recessive mutations are lost both within POD zones and elsewhere in the genome (Fig. 4). Selection against low-fitness recombinants might even favor the evolution of reduced rates of recombination within POD zones providing another mechanism to stabilize POD zones (cf. Olito et al 2022). We ignore the potential of POD zones to gain strength over time by accumulating additional internal mutations sheltered from selection as heterozygotes, which would augment the overdominance as observed at the S-locus in *Arabidopsis halleri* – (Llaurens et al, 2009)).

### 5.2 Evolutionary consequences of POD selection

POD zones could affect the architecture and the dynamics of the genetic load in various ways. Most conspicuously, our simulations of background selection show how POD zones could increase the segregational load elsewhere in the genome and vice versa. Such effects imply that mutations both within and outside the POD zone could reinforce the selection maintaining POD zones sustaining more variability and segregating loads than otherwise expected. Such loads could favor self-incompatibility mechanisms for their ability to produce fewer low-fitness homozygous genotypes. Our scenario where population hybridization spawns POD zones suggests a mechanism whereby fixed drift loads might regularly be converted into segregating loads which then persist in regions expressing strong overdominance.

Although we expect positive heterozygosity-fitness correlations within partially inbred populations (given that heterozygosity inversely measures inbreeding), heterozygosity and variation within POD zones reflects the opposite: non-adaptive variation emerging from sustained mutational and segregational genetic loads. This may help to explain why heterozygosity-fitness correlations can be weak and inconsistent (David, 1998). POD zones might increase loads within populations by creating safe havens within which new deleterious mutations could accumulate while increasing the load of mutations segregating elsewhere in the genome. Small, inbred populations might also become vulnerable to “mutational meltdown” threatening population viability (Gabriel et al, 1993). Conversely, POD zones may provide individual or population advantages by sustaining inbreeding depression and favoring outcrossing in ways that better sustain adaptive genetic variability.

### 5.3 POD effects on mating system evolution

The presence of POD conspicuously affects the evolution of plant and animal mating systems by sustaining more segregational load and higher inbreeding depression than expected especially in small, inbred populations. Early models of mating system evolution sought to explain variable levels of self-fertilization as equilibria reflecting how selection acted on progeny with more or less inbreeding depression. In these simple static models, inbreeding depression less than 0.5 would result in exclusive selfing while higher levels would favor exclusive outcrossing. More dynamic simple models that allow selection make mixed mating systems even more improbable by allowing inbreeding to purge deleterious mutations, generating “run-away” selection for ever-increasing levels of selfing (Lande and Schemske, 1985b). If drift instead fixes many segregating mutations, similar effects emerge as this, too, causes inbreeding depression to decline. The ability of many small, inbred populations to nevertheless retain genetic variation and inbreeding depression plus the absence of purely inbreeding taxa thus pose a paradox (Byers and Waller, 1999; Winn et al, 2011). More complex and realistic models that incorporate effects of linkage, drift, and the associations among loci that arise in small, inbred populations show far more complex dynamics (Charlesworth and Charlesworth, 1987; Uyenoyama et al, 1993). One relevant model showed that a single unlinked over-dominant viability locus anywhere in the genome generates positive associations with modifier alleles enhancing outcrossing (Uyenoyama and Waller, 1991). Such associations favor a persistently mixed mating system. Because POD also favors heterozygotes, we expect POD zones to exert similar effects. The presence of POD zones might thus help to account for the paradoxes of persistent segregating loads and populations and species that maintain mixed mating systems. If, instead, POD zones regularly arise and then deteriorate, selection could alternately favor selfing and outcrossing. This might provide an entirely different mechanism favoring mixed mating systems.

### 5.4 Conclusions

Understanding the mechanisms that create and sustain POD zones cast light on how commonly POD zones may arise and persist and the genetic and demographic circumstances that enhance their longevity. Comparative genomic data will be particularly useful for searching for POD zones and analyzing their structure and history. Our models demonstrate how several genetic, demographic, and mating system parameters may affect load dynamics within and beyond POD zones. Any POD zones that persist are likely to strongly affect mating system evolution by reducing both purifying selection and drift, sapping the power these forces would otherwise have to reduce inbreeding depression. Our models demonstrate that POD zones can persist given the right conditions. We encourage further research to extend and refine our understanding of this phenomenon.

## Supporting information

R script to generate figures

## 6 Acknowledgements

We thank Sylvain Glémin, Lei Zhao, Yaniv Brandvain and an anonymous reviewer for useful comments. Diala Abu Awad was funded by the Alexander von Humboldt Stiftung.

## Supplementary File 1

### Overdominant selection and inbreeding

#### A1 Genetic frequencies at equilibrium

Here we give the full terms used to obtain the equations in the main text. From Appendix A4 from Kimura and Ohta (1971), we have a population with a single bi-allelic overdominant locus, carrying alleles *A* and *B*. The equilibrium frequencies of the three genotype (*P*_11_, *P*_12_ and *P*_22_ for *AA, AB* and *BB*) are:

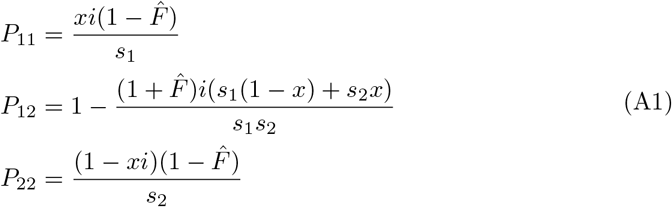

with *s*_1_ and *s*_2_ the coefficients of selection associated with *P*_11_ and *P*_22_ respectively. As mentioned in the main text *x*, the frequency of allele *A*, and *i* are:

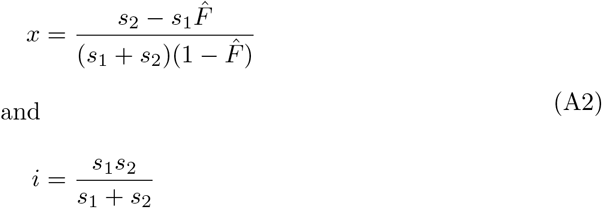

#### A2 Coefficient of inbreeding 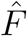

The coefficient of inbreeding for overdominant selection 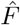 is expressed in terms of the coefficients of selection and the rate of self-fertilisation *σ*:

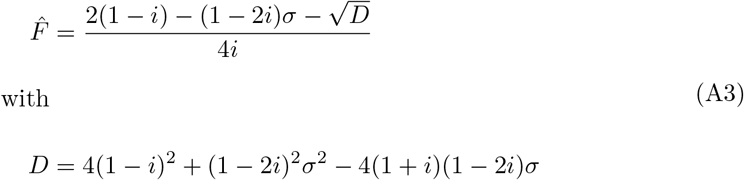

As shown in Figure A1, the smaller the selection against homozygotes, the more 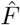 becomes equivalent to the classically used definition of the coefficient of inbreeding, which only depends on the rate of self-fertilization *σ*, 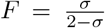 (see Glémin, 2021). However, very strong selection against homozygotes completely cancels the effect of selffertilization on genetic frequencies, with 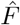 tending to zero for all values of *σ*.

**Figure A1:**
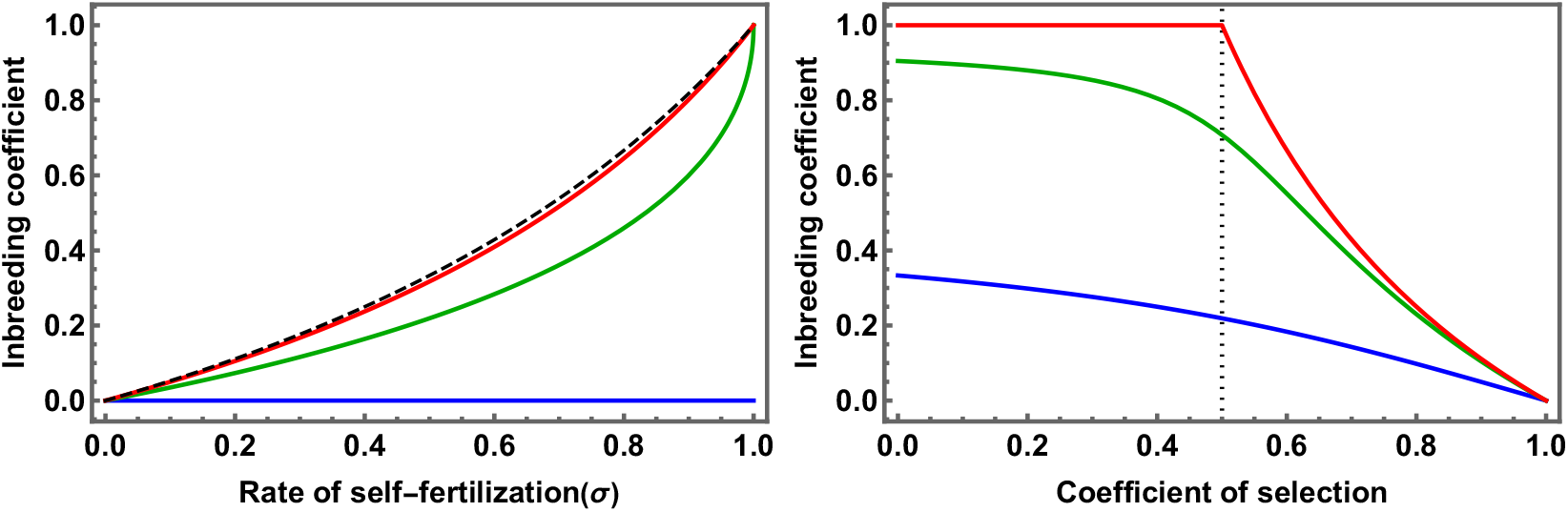
The effect of self-fertilisation and selection on the value of 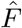 (equation A3). In the left panel, we compare the coefficient of inbreeding *F* (black, dashed line) to 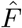 for *s*_1_ = *s*_2_ = 0.1 (red line), 0.5 (green line) and 1 (lethal homozygotes, blue line). In the right panel we show the value of 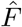 as a function of the coefficient of selection *s_H_* (i.e considering *s*_1_ = *s*_2_ out of convenience) for three values of *σ* (0.5, 0.95 and 1, in blue, green and red, respectively). The black dotted line is set at *s_H_* = 0.5 to highlight the threshold above which overdominance is stable for any value of *σ*.

#### A3 Stability of polymorphism with overdominance

From Equation 1 in the main text, we can plot the conditions necessary, with regards to the values of *s*_1_, *s*_2_ and σ for which there is a stable polymorphic equilibrium (*i.e*. 0 > *x* < 1, see also Figure A.1 in Kimura and Ohta (1971), Appendix 4). From this figure, it is clear that the maintenance of a stable equilibrium is more constrained the higher the rate of self-fertilisation.

#### A4 Inbreeding depression *δ_od_* (overdominant selection)

The expected level of inbreeding depression (*δ*) depends on the fitnesses of selfed and outcrossed offspring, respectively *W_s_* and *W_o_*. These variables can be expressed in

**Figure A2:**
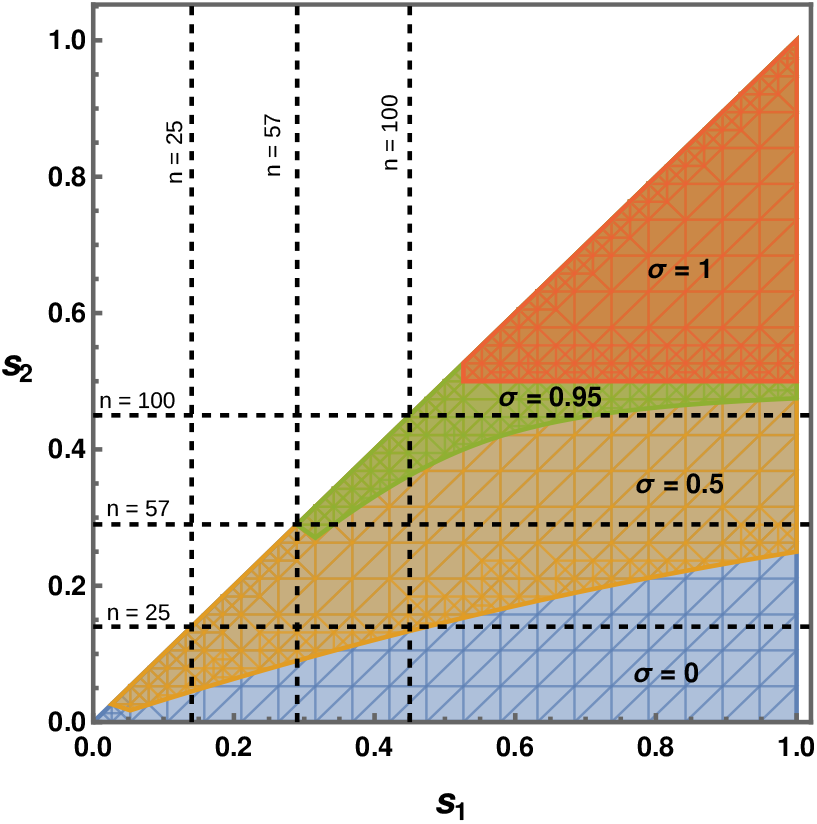
Representing the combinations of *s*_1_ and *s*_2_ (assuming *s*_1_ > *s*_2_) for which the inequality presented in Equation 1 from the main text is true (the regions above and to the right of the lines) for different values of σ (0, 0.5, 0.95 and 1, in blue, orange, green and red, respectively). The dotted lines represent the coefficients of selection based on the number of loci carrying mutations with a coefficient of selection *s* = 0.01 and dominance *h* = 0.2 (see Eq. 3 in the main text).

terms of the coefficients of selection associated with each homozygote and the expected genotypic frequencies (see Charlesworth and Charlesworth 1990):

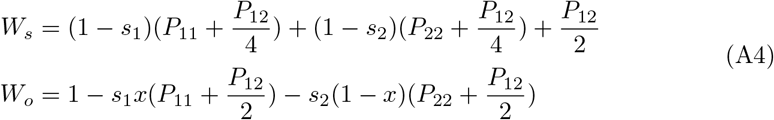

Using the expressions for the genotypic frequencies given in Eq. A1and the definitions in Eq. A2, and setting *s*_1_ = *s*_2_, we obtain the simplified expression for *δ*, given in Eq. 9 of the main text.

## Supplementary File 2

### Selection for recombinant haplotypes

#### Coefficients of selection associated with recombinants

Any recombinant haplotype *H_c_* will have a heterozygote advantage that depends on the haplotype with which it is inherited (*H*_1_ or *H*_2_). If we assume that no mutation has been cleaved during recombination, and that the fitness of reference is that of the initial heterozygote *H*_1_*H*_2_ (as in Eq. 3 in the main text) then the coefficient of selection against the homozygote *H_c_H_c_* is simply *s_c_* = *s_H_*. The coefficients of selection against heterozygotes *H_c_H*_1_ and *H_c_H*_2_ are given by:

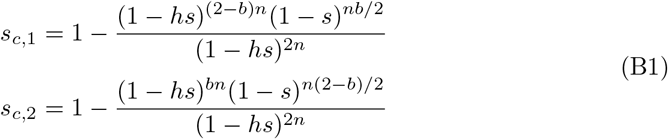

Here *n* is the number of loci carrying deleterious mutations. The parameter *b* (0 ≤ *b* ≤ 2) reflects the similarity between the recombinant haplotype *H_c_* and (arbitrarily) haplotype *H*_1_. When *b* ≈ 0 (respectively 2), this implies that the recombinant is made up mostly of the haplotype *H*_2_ (respectively *H*_1_), giving *s*_*c*,1_ =0 (respectively *s*_*c*,1_ = *s_H_*). If *b* =1 then *H_c_* is made up of equal parts of *H*_1_ and *H*_2_ giving *s*_*c*,1_ = *s*_*c*,2_.

**Figure B1:**
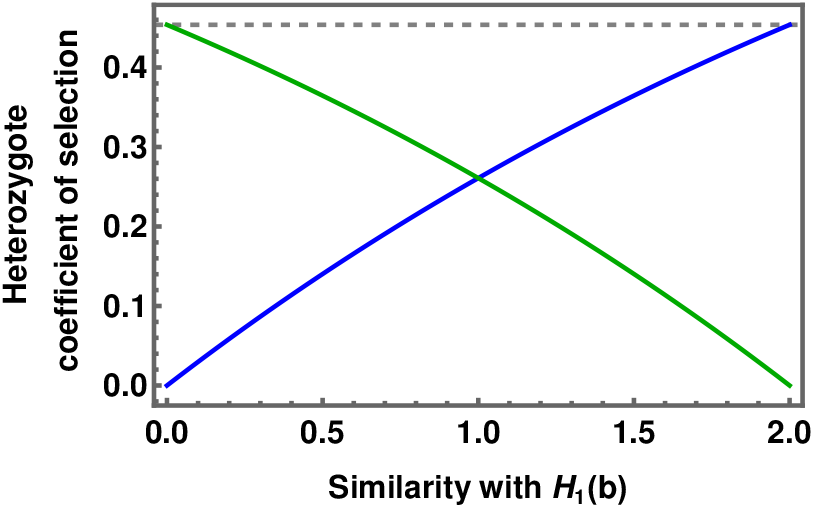
The relative values of the coefficients of selection against heterozygotes *H_c_H*_1_ and *H_c_H*_2_, respectively *s*_*c*,1_ (blue line) and *s*_*c*,2_ (green line) for different values of parameter b (See Eq. B1. The dotted line represents *s_H_*, the coefficient of selection against homozygotes.

In the case of a loss of a single mutation during recombination, arbitrarily initially present on *H*_1_, then the coefficients of selection involving *H_c_* become:

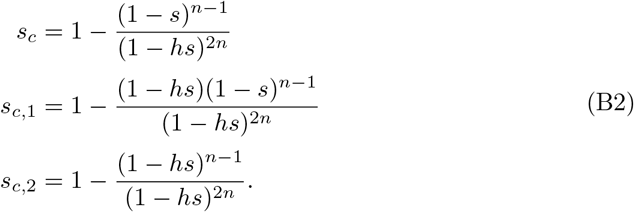

#### Changes in frequencies and equilibrium expectations

Previous works have shown that several overdominant haplotypes can co-exist when heterozygotes all have the same relative fitness of 1 (see for example Glémin, 2021). From the expressions for the coefficients of selection in Eqs. B1 and B2it is clear that this assumption cannot be made in the context of POD. To evaluate whether more than two haplotypes can exist, we derive expressions for the changes in frequencies *P*_1_, *P*_2_ and *P_c_* of haplotypes *H*_1_, *H*_2_ and *H_c_* respectively:

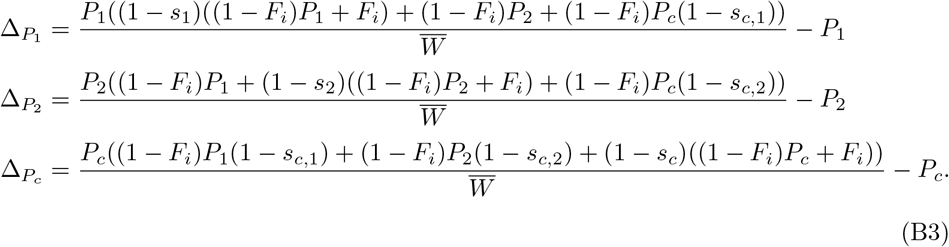

where 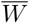 is the mean expected fitness, given by:

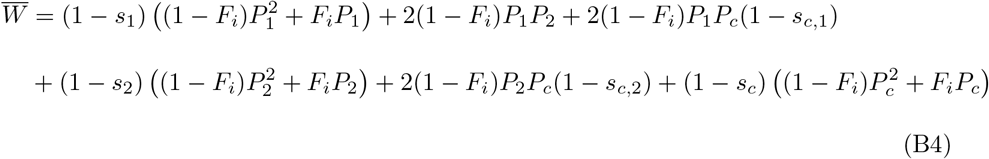

and *F_i_* is a generic coefficient of inbreeding, which may not necessarily be equal to 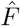 (see Eq. A3 in Supp. File 1) in a context of overdominance with three alleles or haplotypes. Setting *s*_1_ = *s*_2_ and using Wolfram Mathematica (REF) to resolve the system of Ordinary Differential Equations above, we find that the only possible equilibria would be for either the fixation of *H*_1_, *H*_2_ or *H_c_*, or the loss one of the haplotypes.

As shown in the main text (Eq. 12), by assuming *P_c_* is of order *ϵ* (*ϵ* being very close to zero), we can determine whether a rare recombinant *H_c_* can increase in frequency in a population at equilibrium for the two initial haplotypes. We derive the expression for Δ*_P_c__* to the first order of *ϵ* with *s_H_* = *s*_1_, *s*_2_ and *P*_1_ = *P*_2_ = (1 – *ϵ*)/2:

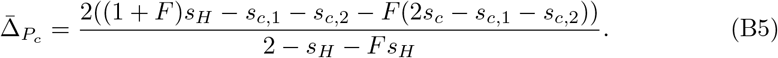

Deriving the above expression while considering 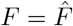 gives an expression for Δ_*P_c_*_ that depends on the initial coefficient of selection of the recombinant haplotype *s_c_*:

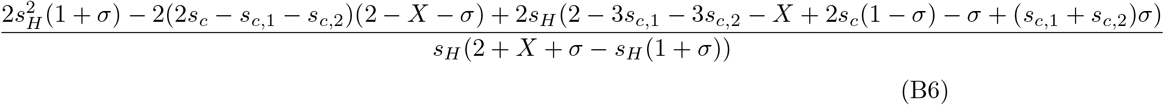

with 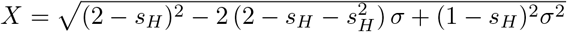

If *s_c_* = *s_H_*, the same conditions as presented in the two-haplotype case (see Supp. File 1) need to be filled for *H_c_* to replace either of the initial haplotypes (i.e. *H_c_* must be similar enough to either *H*_1_ or *H*_2_ to replace the genotype in question). This is the case when *b* ≈ 0 or 2 (see Equation B1 and Fig. B2). As Δ*_P_c__* ≈ 0, *H_c_* is neither selected for or against, and drift should play a major role in determining the frequency *P_c_*. For 0 < *b* < 2 the recombinant will always be selected against (Δ*_P_c__* < 0).

**Figure B2:**
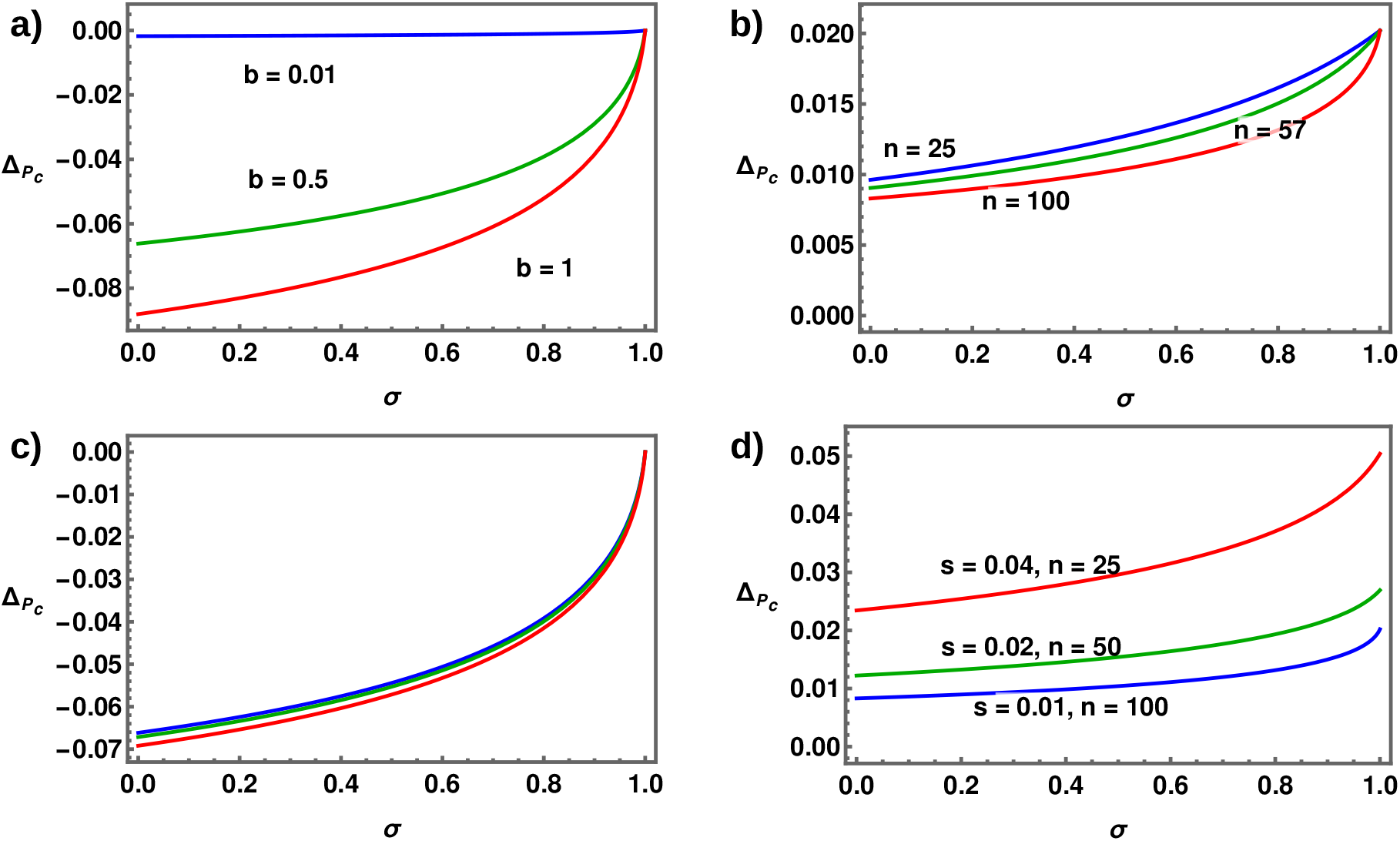
The change in frequency of a haplotype *H_c_* (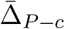, see Eq. B5) as a function of the selfing rate *σ*. a) and c) No mutations are lost and *n* = 100, b) and d) a single mutation is cleaved. a) Each coloured line represents a different value of the parameter *b* (see Eq. B1 and accompanying text) for *n* = 100. b) Each coloured line represents a different value of *n*. For both a) and b) *s* = 0.01 and *h* = 0.2. c) and d) show values of 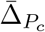 for the same *s_H_*, but for different *s* and *n* (but fixed *h* = 0.2).

## Supplementary Figures

**Figure S1:**
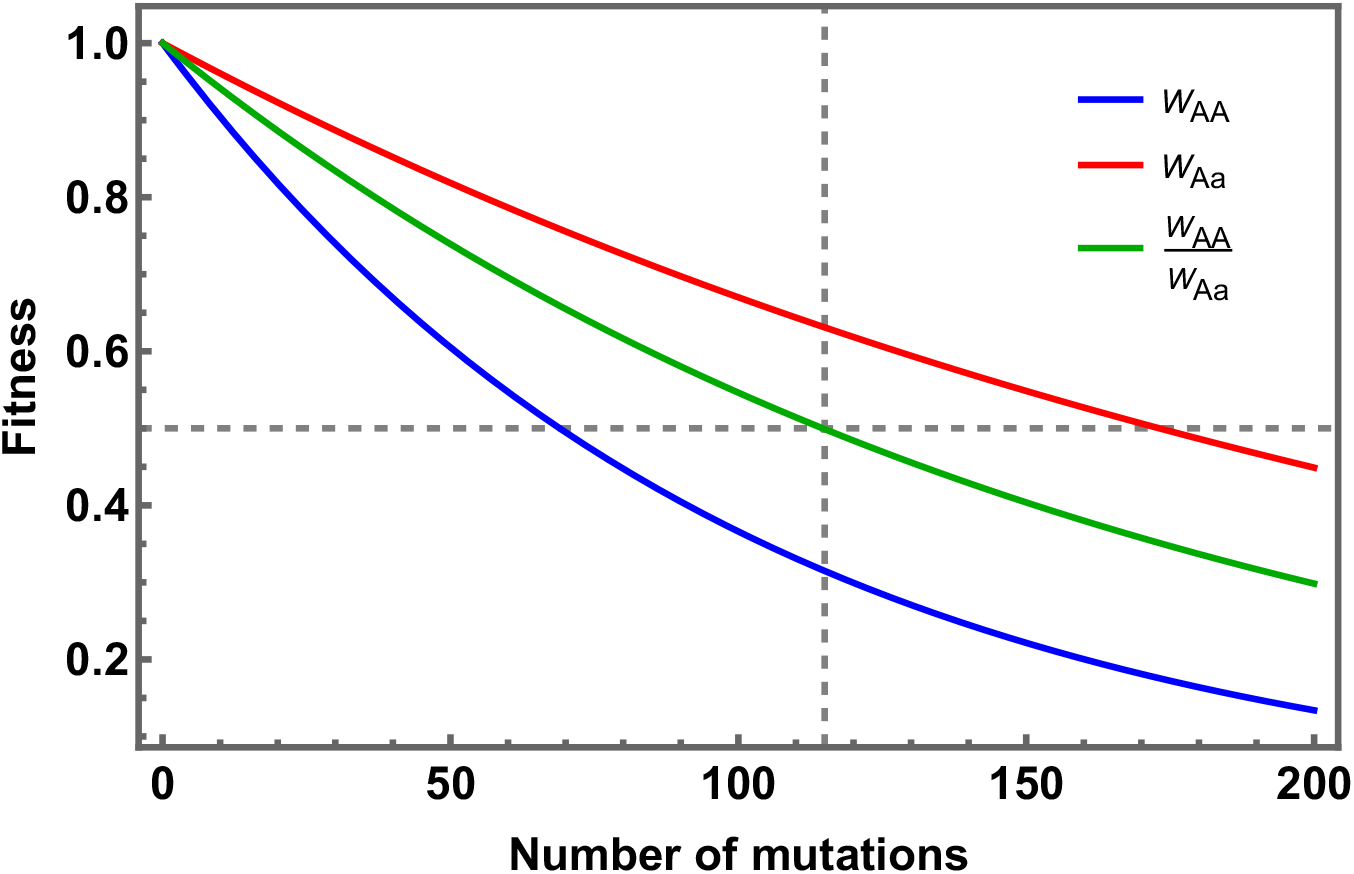
Declines in homozygote (AA) and heterozygote (Aa) fitness and the fitness of homozygotes relative to heterozygotes as a function of the number of completely linked mutations expressing pseudo-overdominance. Mutations are identical, slight (*s* = 0.01), recessive (dominance *h* = 0.2), and have independent (multiplicative) fitness effects. Homozygote fitness is given by (1 – *s*)^*n*^ while heterozygote fitness is (1 – *hs*)^2*n*^ (see Eqs. 2 and 3 in the main text). The horizontal dashed line shows the threshold fitness at which homozygote fitness drops to less than half the fitness of the heterozygote - a condition for POD indefinite persistence even in fully selfing populations. For these parameter values, this occurs for *n* = 115 (the vertical dashed line). (cf. Supp. File 1 Fig. A2).

**Figure S2:**
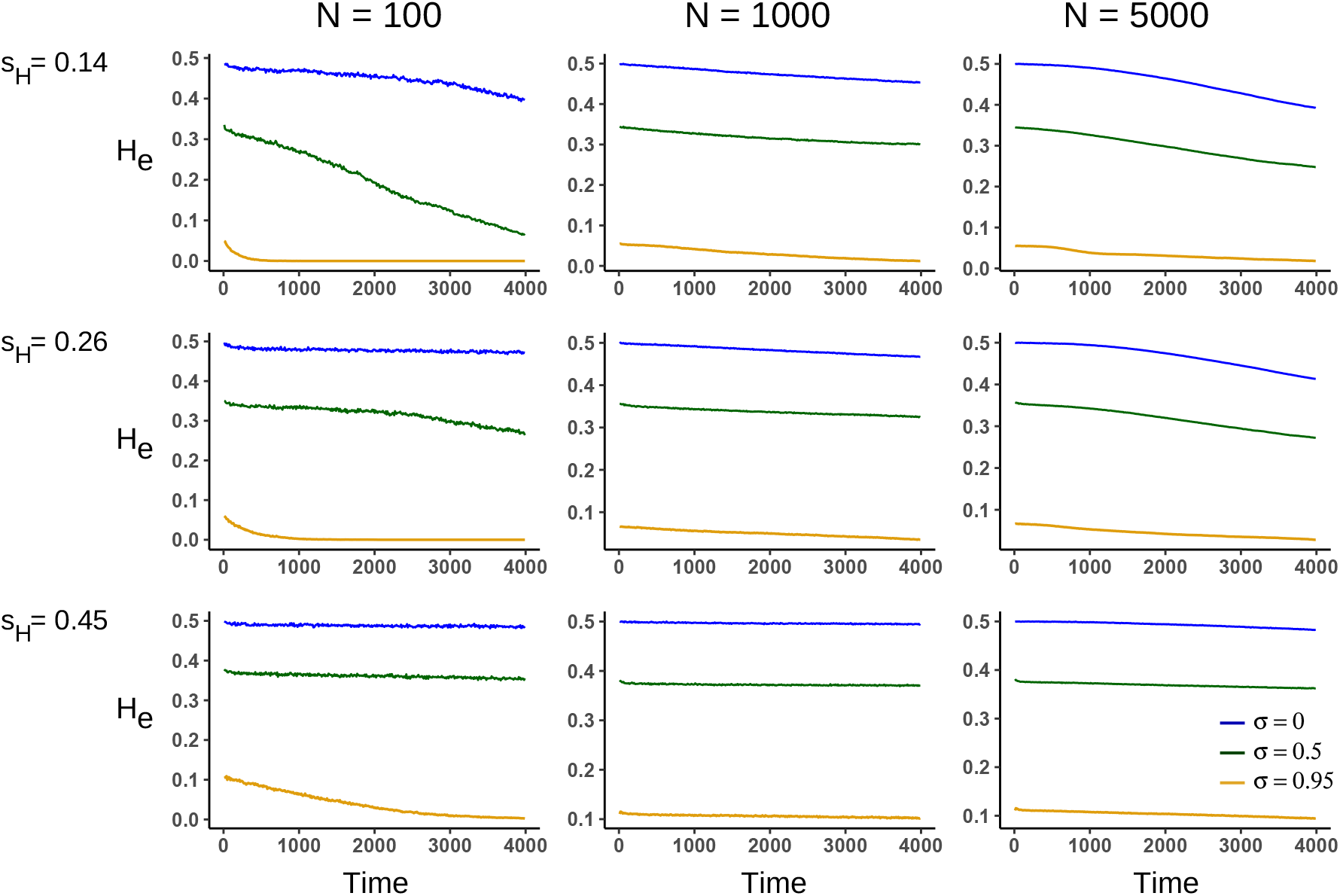
Mean declines in heterozygosity *H_e_* with time for different population sizes, *N*, and selection coefficients against homozygous haplotypes, *s_H_*. Simulations reflect three selfing rates (*σ* = 0, 0.5 and 0.95). Individual loci within the POD zone have a selection coefficient *s* = 0.01, dominance *h* = 0.2, and map length (or the recombination rate)between loci in the POD zone is *ℓ* = 10^-5^. There are no background mutations.

**Figure S3:**
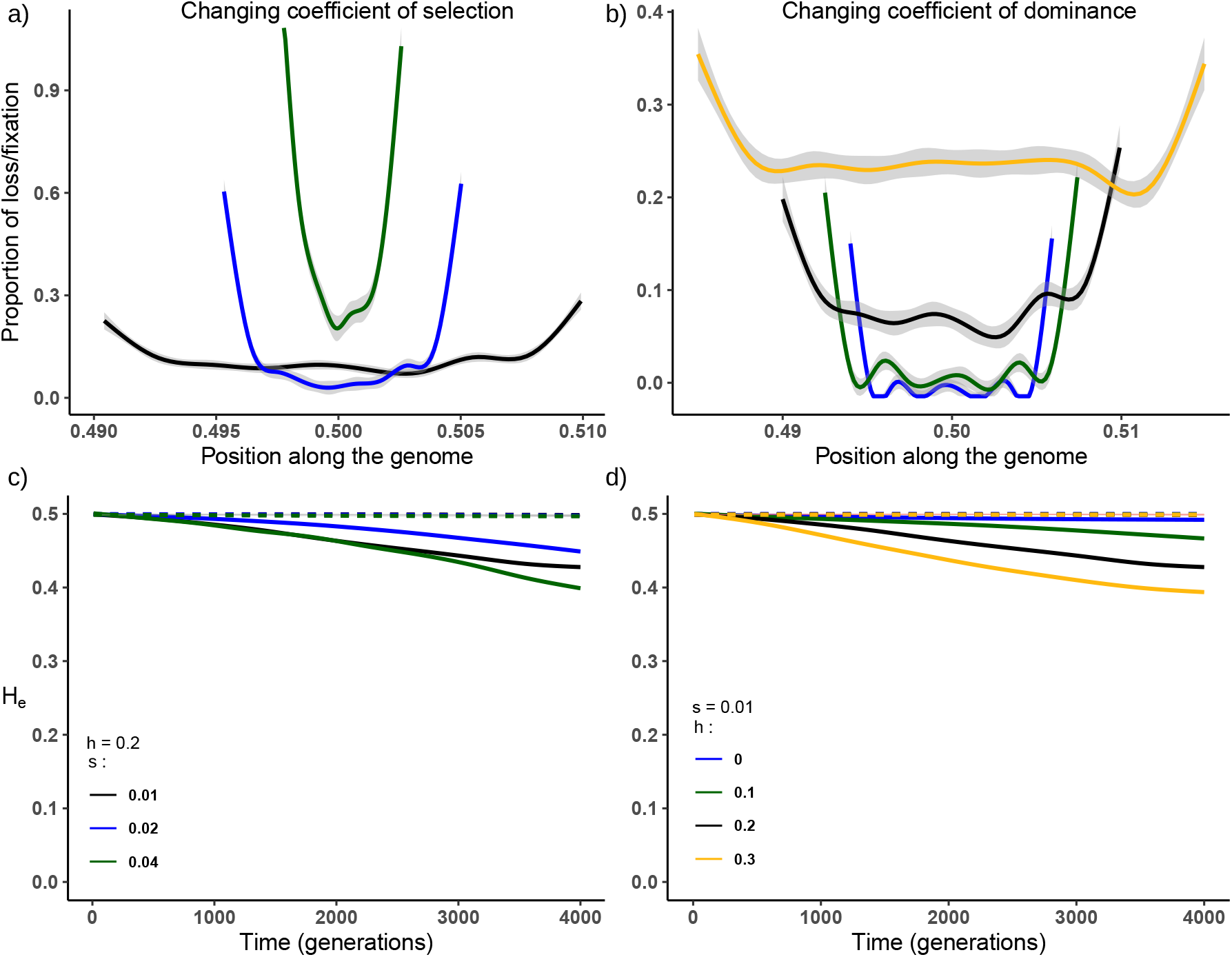
Effects of levels of selection and dominance on selection dynamics within a POD zone (compare to Fig. 2). Left panels show the effects of varying coefficients of selection at each load locus (*s* = 0.01, 0.02 and 0.04, corresponding to *n_L_* = 100, 50 and 25 loci). Dominance is fixed at *h* = 0.2 and *s_H_* = 0.45. Right panels show the effects of varying dominance (*h* = 0, 0.1, 0.2 and 0.5 with *n_L_* = 60, 75, 100 and 150) with selection fixed at *s* = 0.01. Panels a) and b) show observed frequencies of fixation/loss along the POD zone (x values represent the position of the loci along the chromosome). The selfing rate *σ* = 0 and linkage *ℓ* = 10^-4^M. Panels c) and d) show losses in heterozygosity (*H_e_*) over time in populations with a high selfing rate (*σ* = 0.95) and either loose linkage (*ℓ* = 10^-4^*M*, solid lines) or tight linkage (*ℓ* = 10^-5^*M*, dashed lines).

**Figure S4:**
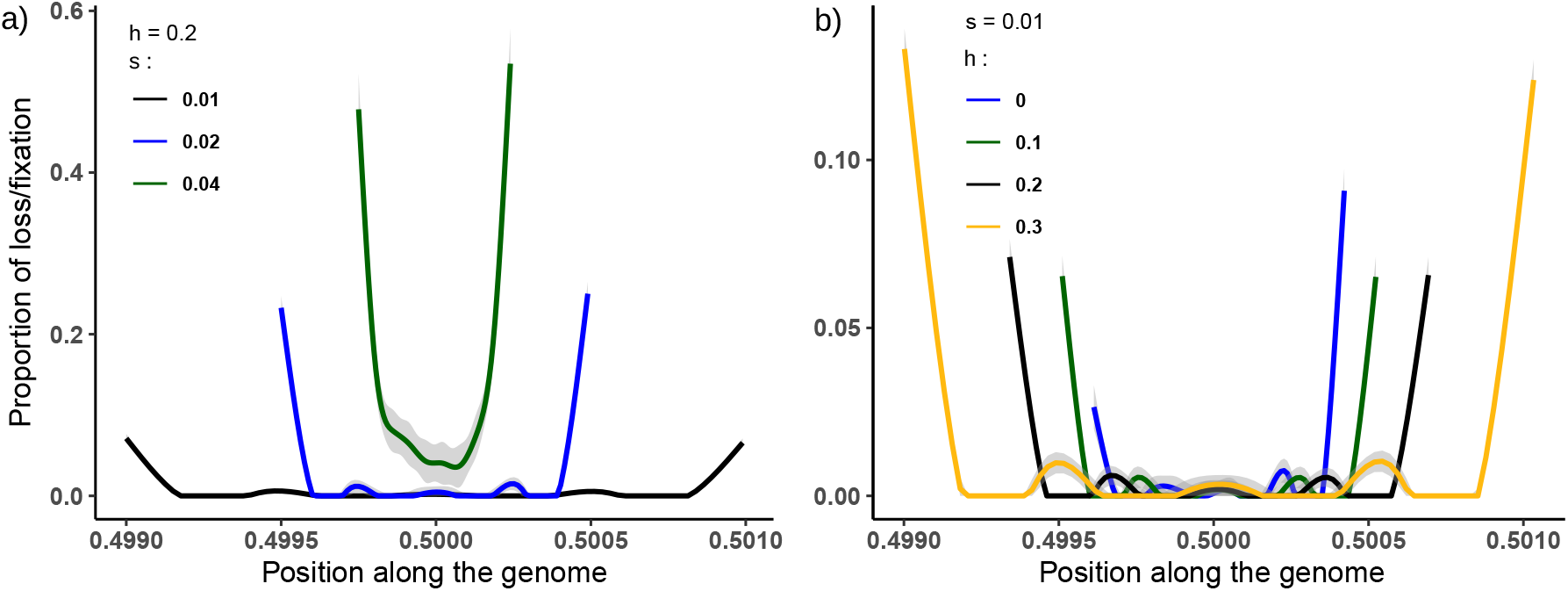
Effects of levels of selection and dominance on selection dynamics within a POD zone. Observed frequencies of fixation/loss along the POD zone (x values represent the position of the loci along the chromosome) after 4000 generations. The selfing rate *σ* = 0.95 and linkage *ℓ* = 10^-5^*M*. a) Effects of varying the coefficient of selection at load loci (*s* = 0.01, 0.02 and 0.04, corresponding to *n_L_* = 100, 50 and 25 loci) with dominance fixed at *h* = 0.2 and *s_H_* = 0.45. b) Effects of varying dominance (*h* = 0, 0.1, 0.2 and 0.5 with *n_L_* = 60, 75, 100 and 150) with selection fixed at *s* = 0.01.

**Figure S5:**
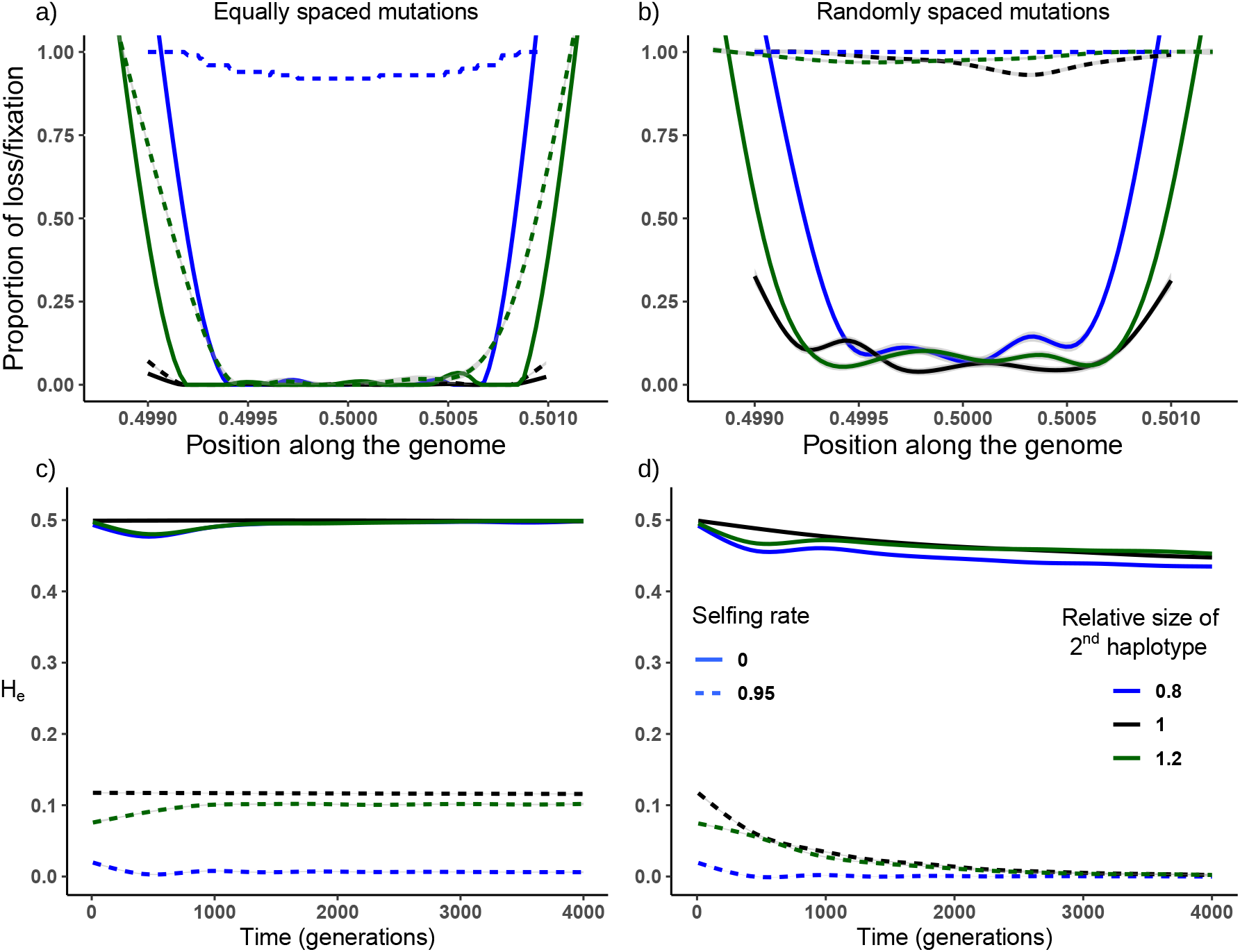
Effects of relaxing the assumptions of symmetric overdominance and evenly spaced mutations. Upper panels show locations within the POD zone where load mutations are most likely to be lost after 4000 generations (a, b) and how this depends on whether mutations are evenly spaced (a) or randomly distributed (b). Results are shown for both symmetric (black) and asymmetric (green and blue) loads. Outcomes under both outcrossing and high selfing (solid vs. dotted lines) are shown. Note erosion of mutations via recombination and selection at both ends of the POD zone. Lower panels show overall stability of the POD zone (shown as heterozygosity, *He*) over time. As in the upper panels, graphs show results for both symmetric (black) and asymmetric (green and blue) loads and for evenly (panel c) and randomly (panel d) placed mutations. The coefficients of selection and dominance are *s* = 0.01 and *h* = 0.2 respectively, linkage within the POD zone is *ℓ* = 10^-5^ and population size *N* = 1000.

**Figure S6:**
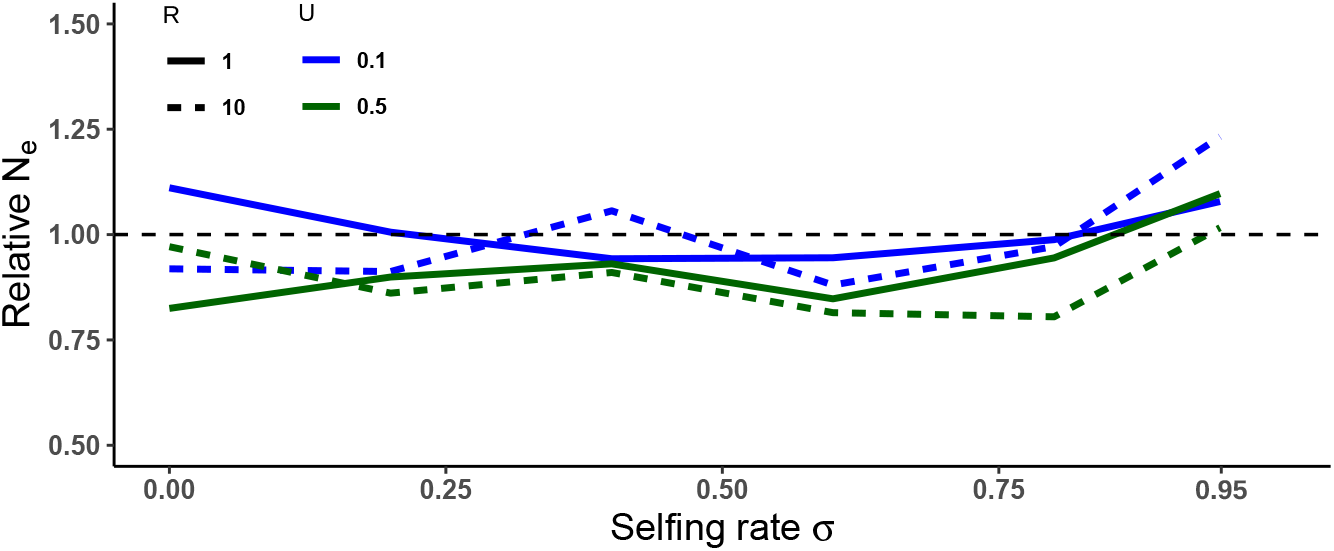
Effective population size, *N_e_*, in a populations supporting a POD zone relative to a population without one (done by setting *s* = *h* = 0). *N_e_* varies as a function of the selfing rate for populations subject to different background mutation rates (*U*) and shorter and longer map lengths (*R* in Morgans). These simulations use 100 POD load loci (*n_L_* = 100) and a map length of *ℓ* = 10^-6^ Morgans. Mutations within the POD zone are randomly placed. Selection coefficients in- and outside the POD zone (*s* and s_d_ respectively) are 0.01 with dominances *h* and *h_d_* = 0.2.

**Figure S7:**
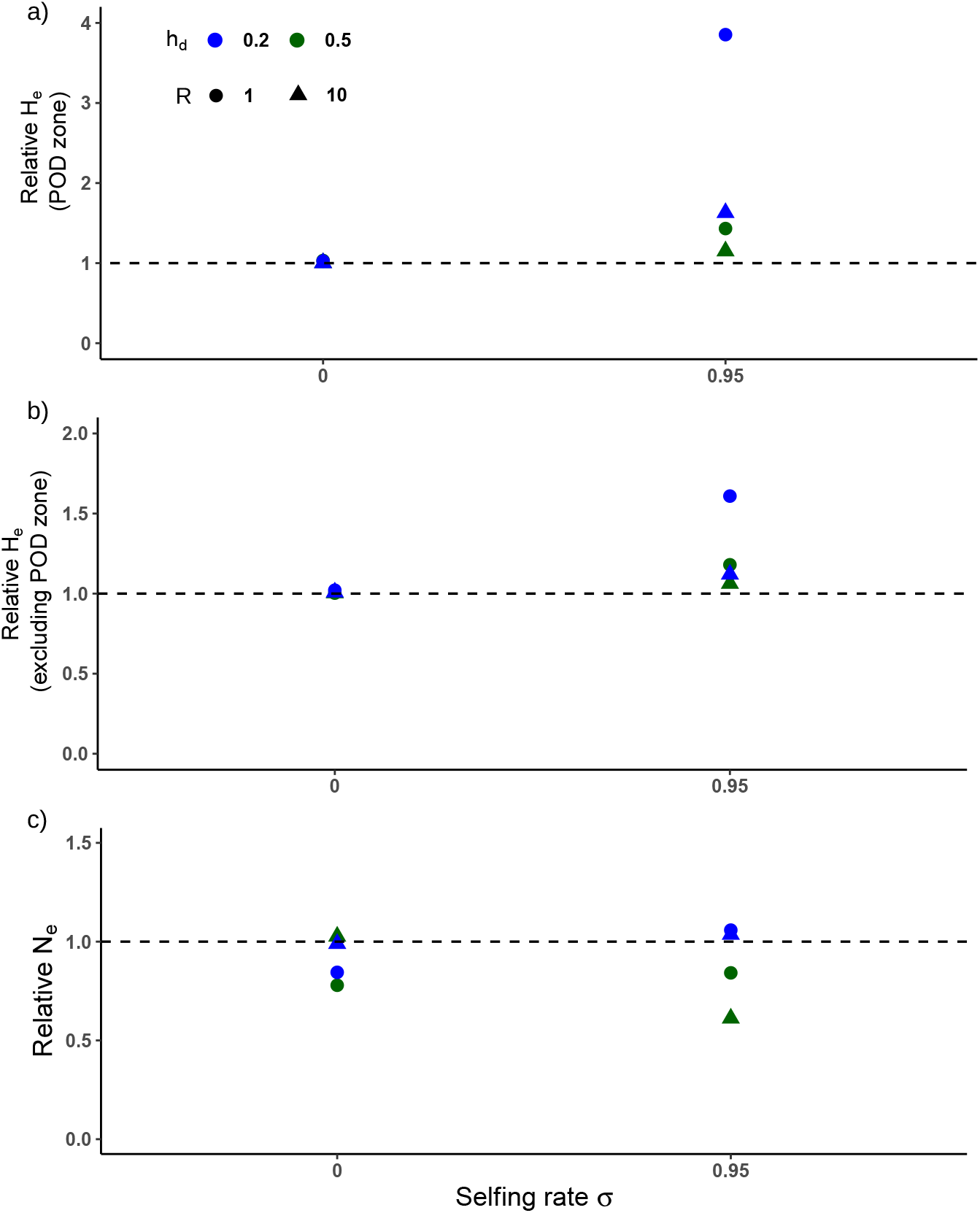
Associative-Overdominance of background mutations affects selection dynamics within POD zones and vice versa. Graphs compare the effect of recessive (*h_d_* = 0.2 blue) to codominant (*h_d_* = 0.5 green) background mutations on population heterozygosity as a function of the selfing rate. a) Heterozygosity, *H_e_*, within a POD zone with background mutations occurring elsewhere in the genome relative to *H_e_* in a population lacking background mutations. b) *H_e_* outside the POD zone in a population with a POD zone relative to one without. c) *N_e_* in a population with POD selection relative to one without. The background mutation rate *U* = 0.5 and the map lengths simulated are *R* =1 and 10 Morgans. These simulations use 100 POD load loci (*n_L_* = 100) randomly placed within the POD zone over a map length of *ℓ* = 10^-6^ Morgans. Selection coefficients within the POD zone are *s* = 0.01with dominance *h* = 0.2 and population size *N* = 1000.

**Figure S8:**
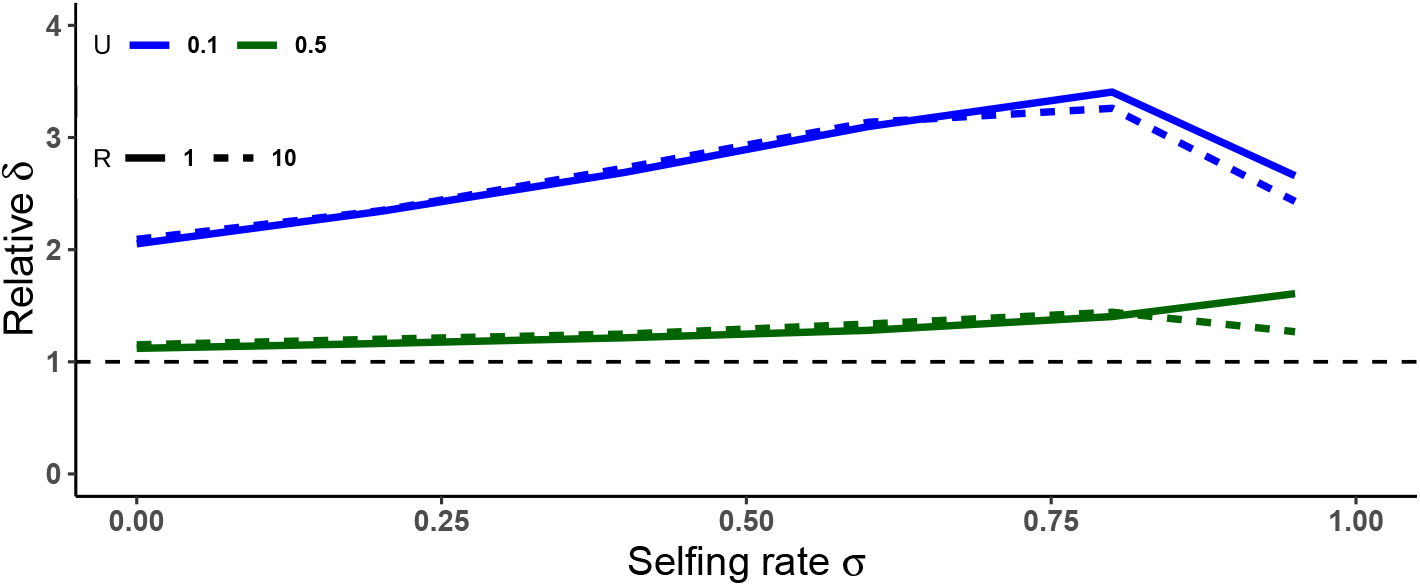
Inbreeding depression in a population with a POD zone relative to one without as a function of the selfing rate *σ* at two genomic mutations rates, *U*, and two genomic map lengths, *R*. Other parameter values are *n_L_* = 100 and *ℓ* = 10^-6^ Morgans with mutations randomly placed within the POD zone. Selection coefficients within and outside the POD zone, *s* and *s_d_* respectively are set to 0.01, and dominances *h* and *h_d_* = 0.2.

**Figure S9:**
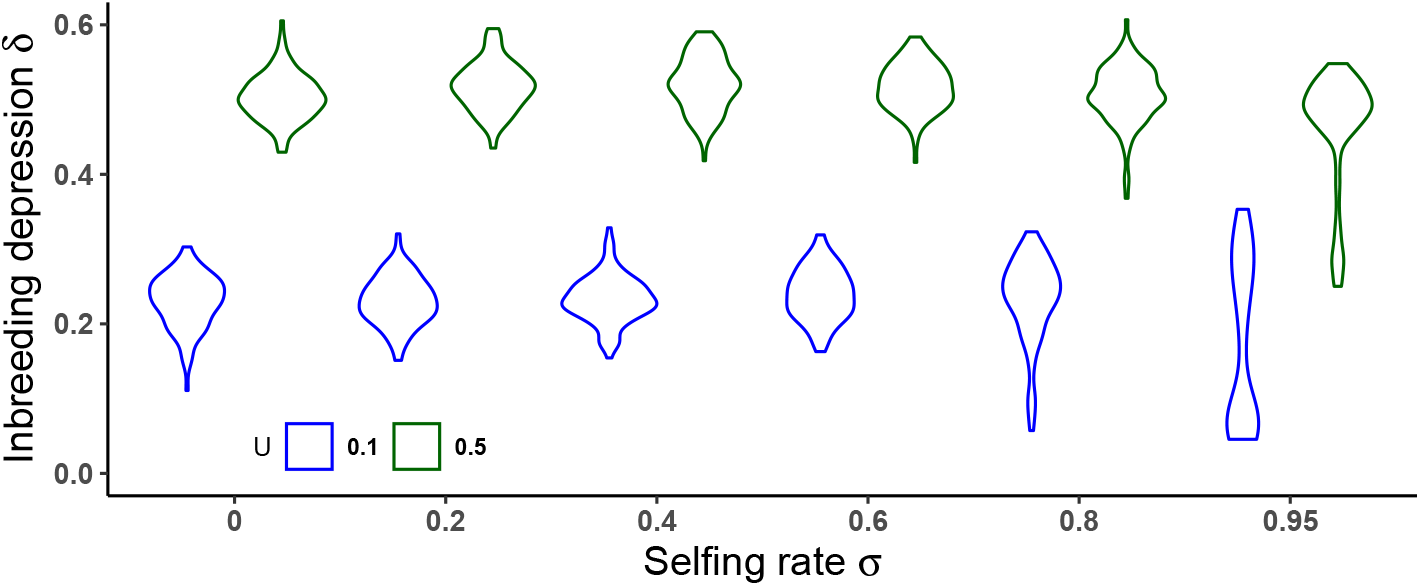
Distributions of inbreeding depression, *δ*, as a function of the population selfing rate in populations with a higher (*U* = 0.5) or lower (*U* = 0.1) haploid genomic mutation rate. Other parameter values: *n_L_* = 100, *ℓ* = 10^-6^ Morgans with mutations randomly placedin the POD zone. Selection coefficients within and outside the POD zone, *s* and *s_d_* respectively are set to 0.01, and dominances *h* and *h_d_* = 0.2. The genome size (setting the recombination rate) is *R* = 10 Morgans.

**Figure S10:**
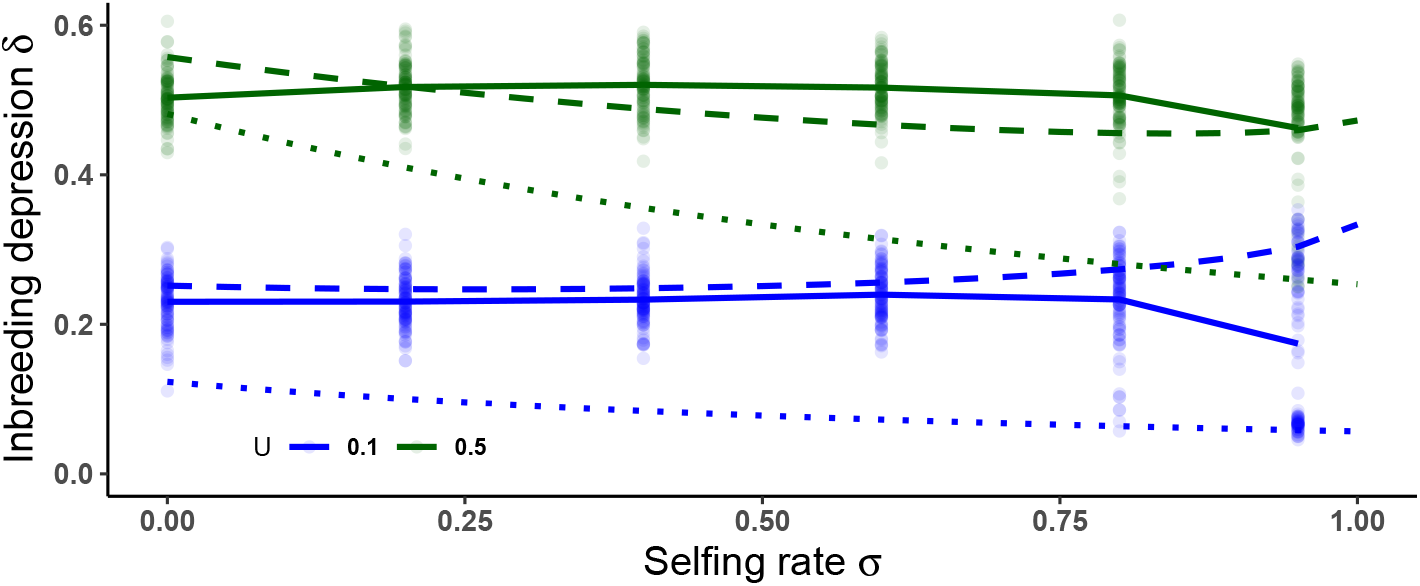
POD zones increase inbreeding depression. Inbreeding depression, *δ*, as a function of the selfing rate for populations with higher (*U* = 0.5) and lower (*U* = 0.1) haploid genomic mutation rates. Solid lines represent means of the simulations run with a POD zone while dotted lines show the *δ* expected in the absense of overdominance (Eq. (7)). Dashed lines show the *δ* expected with overdominant selection (Eq. (6)). Other parameter values are *n_L_* = 100 and *ℓ* = 10^-6^ Morgans. Mutations are randomly placed within the POD zone. Selection coefficients within and outside the POD zone, *s* and *s_d_* respectively are set to 0.01, and dominances *h* and *h_d_* = 0.2. The genome size setting the recombination rate is *R* = 1 Morgans.

## Bibliography

Abu Awad D, Waller DM (2022) POD selection - individual based model. DOI 10.5281/ZENODO.6874327, URL https://zenodo.org/record/6874327

Baldwin SJ, Schoen DJ (2019) Inbreeding depression is difficult to purge in self-incompatible populations of *Leavenworthia alabamica*. New Phytologist 224(3):1330–1338, DOI 10.1111/nph.15963, URL https://onlinelibrary.wiley.com/doi/10.1111/nph.15963

Bataillon T, Kirkpatrick M (2000) Inbreeding depression due to mildly deleterious mu-tations in finite populations: size does matter. Genetical Research 75(1):75–81, DOI 10.1017/S0016672399004048

Bernstein MR, Zdraljevic S, Andersen EC, Rockman MV (2019) Tightly linked antagonistic-effect loci underlie polygenic phenotypic variation in C. elegans. Evolution Letters 3(5):462–473, DOI 10.1002/evl3.139, URL https://onlinelibrary.wiley.com/doi/10.1002/evl3.139

Brandenburg JT, Mary-Huard T, Rigaill G, Hearne SJ, Corti H, Joets J, Vitte C, Charcosset A, Nicolas SD, Tenaillon MI (2017) Independent introductions and admixtures have contributed to adaptation of european maize and its american counterparts. PLoS Genetics 13(3), DOI 10.1371/journal.pgen.1006666

Brown KE, Kelly JK (2020) Severe inbreeding depression is predicted by the “rare allele load” in *Mimulus guttatus*. Evolution 74(3):587–596, DOI 10.1111/evo.13876, URL https://onlinelibrary.wiley.com/doi/abs/10.1111/evo.13876

Byers D, Waller D (1999) Do plant populations purge their genetic load? effects of population size and mating history on inbreeding depression. Annual Review of Ecology and Systematics 30:479–513, DOI 10.1146/annurev.ecolsys.30.1.479

Charlesworth B (2018) Mutational load, inbreeding depression and heterosis in subdivided populations. Molecular Ecology 27(24):4991–5003, DOI 10.1111/mec.14933

Charlesworth B, Nordborg M, Charlesworth D (1997) The effects of local selection, balanced polymorphism and background selection on equilibrium patterns of genetic diversity in subdivided populations. Genetics Research 70(2):155–174, DOI 10.1017/S0016672397002954, URL https://www.cambridge.org/core/journals/genetics-research/article/effects-of-local-selection-balanced-polymorphism-and-background-selection-on-equilibrium-patterns-1F946DDC83F5277DC964D76329A6BB66

Charlesworth D, Charlesworth B (1987) Inbreeding depression and its evolutionary consequences. Annual Review of Ecology, Evolution, and Systematics 18:237–268

Charlesworth D, Charlesworth B (1990) Inbreeding Depression with Heterozygote Advantage and Its Effect on Selection for Modifiers Changing the Outcrossing Rate. Evolution 44(4):870–888, DOI 10.1111/j.1558-5646.1990.tb03811.x, URL https://onlinelibrary.wiley.com/doi/abs/10.1111/j.1558-5646.1990.tb03811.x

Chelo IM, Afonso B, Carvalho S, Theologidis I, Goy C, Pino-Querido A, Proulx SR, Teotónio H (2019) Partial Selfing Can Reduce Genetic Loads While Maintaining Diversity During Experimental Evolution. G3 Genes|Genomes|Genetics 9(9):2811–2821, DOI 10.1534/g3.119.400239, URL https://academic.oup.com/g3journal/article/9/9/2811/6026387

Crow JF (1993) Mutation, mean fitness, and genetic load. Oxford Series in Evol Biol 9:3–42

Crow JF (1999a) The Rise and Fall of Overdominance. In: Plant Breeding Reviews, John Wiley & Sons, Ltd, pp 225–257, DOI 10.1002/9780470650134.ch5, URL https://onlinelibrary.wiley.com/doi/abs/10.1002/9780470650134.ch5

Crow JF (1999b) The rise and fall of overdominance. Plant Breeding Reviews 17:225–257

Crow JF, Kimura M (1970) An Introduction to Population Genetics Theory. Burgess Pub. Co., 591 pp.

Darwin C (1876) The effects of cross and self fertilization in the vegetable kingdom. J. Murray and Co.

David P (1998) Heterozygosity–fitness correlations: new perspectives on old problems. Heredity 80(5):531–537, DOI 10.1046/j.1365-2540.1998.00393.x, URL http://www.nature.com/doifinder/10.1046/j.1365-2540.1998.00393.x

Ehiobu NG, Goddard ME, Taylor JF (1989) Effect of rate of inbreeding on inbreeding depression in Drosophila melanogaster. Theoretical and Applied Genetics 77(1):123–127, DOI 10.1007/BF00292326, URL http://link.springer.com/10.1007/BF00292326

Fisher S Ronald A (1930) The Genetical Theory of Natural Selection. Oxford University Press, Oxford

Gabriel W, Lynch M, Bürger R (1993) Muller’s Ratchet and Mutational Meltdowns. Evolution 47(6):1744–1757, DOI 10.1111/j.1558-5646.1993.tb01266.x, URL https://onlinelibrary.wiley.com/doi/abs/10.1111/j.1558-5646.1993.tb01266.x

Garrigan D, Hedrick PW (2003) Perspective: Detecting Adaptive Molecular Polymorphism: Lessons from the MHC. Evolution 57(8):1707–1722, DOI 10.1111/j.0014-3820.2003.tb00580.x, URL https://onlinelibrary.wiley.com/doi/10.1111/j.0014-3820.2003.tb00580.x

Gemmell NJ, Slate J (2006) Heterozygote Advantage for Fecundity. PLoS ONE 1(1):e125, DOI 10.1371/journal.pone.0000125, URL https://dx.plos.org/10.1371/journal.pone.0000125

Gilbert KJ, Pouyet F, Excoffier L, Peischl S (2020) Transition from Background Selection to Associative Overdominance Promotes Diversity in Regions of Low Recombination. Current Biology 30(1):101–107.e3, DOI 10.1016/j.cub.2019.11.063, URL https://www.cell.com/current-biology/abstract/S0960-9822(19)31565-9

Glémin S (2021) Balancing selection in self-fertilizing populations. Evolution 75(5):1011–1029, DOI 10.1111/evo.14194, URL https://onlinelibrary.wiley.com/doi/10.1111/evo.14194

Harkness A, Brandvain Y, Goldberg EE (2019) The evolutionary response of mating system to heterosis. Journal of Evolutionary Biology 32(5):476–490, DOI 10.1111/jeb.13430, URL https://onlinelibrary.wiley.com/doi/10.1111/jeb.13430

Hedrick PW, Garcia-Dorado A (2016) Understanding Inbreeding Depression, Purging, and Genetic Rescue. Trends in Ecology & Evolution 31(12):940–952, DOI 10.1016/j.tree.2016.09.005, URL https://linkinghub.elsevier.com/retrieve/pii/S0169534716301586

Hedrick PW, Hellsten U, Grattapaglia D (2016) Examining the cause of high inbreeding depression: analysis of whole-genome sequence data in 28 selfed progeny of eucalyptus grandis. New Phytologist 209(2):600–611, DOI 10.1111/nph.13639

Hill WG, Robertson A (1966) The effect of linkage on limits to artificial selection. Genetical Research 89(5-6):311–336, DOI 10.1017/S001667230800949X

Igic B, Lande R, Kohn J (2008) Loss of self-incompatibility and its evolutionary consequences. International Journal of Plant Sciences 169(1):93–104, DOI 10.1086/523362, URL https://doi.org/10.1086/523362, https://doi.org/10.1086/523362

Jay P, Chouteau M, Whibley A, Bastide H, Parrinello H, Llaurens V, Joron M (2021) Mutation load at a mimicry supergene sheds new light on the evolution of inversion polymorphisms. Nature Genetics 53(3):288–293, DOI 10.1038/s41588-020-00771-1, URL https://www.nature.com/articles/s41588-020-00771-1

Kardos M, Allendorf FW, Luikart G (2014) Evaluating the role of inbreeding depression in heterozygosity-fitness correlations: how useful are tests for identity disequilibrium? Molecular Ecology Resources 14(3):519–530, DOI 10.1111/1755-0998.12193, URL https://onlinelibrary.wiley.com/doi/10.1111/1755-0998.12193

Kim BY, Huber CD, Lohmueller KE (2018) Deleterious variation shapes the genomic landscape of introgression. PLOS Genetics 14(10):e1007,741, DOI 10.1371/journal.pgen.1007741, URL https://dx.plos.org/10.1371/journal.pgen.1007741

Kimura M, Ohta T (1971) Theoretical Aspects of Population Genetics, monographs edn. Princeton University Press

Kirkpatrick M (2010) How and Why Chromosome Inversions Evolve. PLoS Biology 8(9):e1000,501, DOI 10.1371/journal.pbio.1000501, URL https://dx.plos.org/10.1371/journal.pbio.1000501

Kirkpatrick M, Jarne P (2000) The Effects of a Bottleneck on Inbreeding Depression and the Genetic Load. The American Naturalist 155(2):154–167, DOI 10.1086/303312, URL https://www.journals.uchicago.edu/doi/10.1086/303312

Kremling KAG, Chen SY, Su MH, Lepak NK, Romay MC, Swarts KL, Lu F, Lorant A, Bradbury PJ, Buckler ES (2018) Dysregulation of expression correlates with rare-allele burden and fitness loss in maize. Nature 555(7697):520–523, DOI 10.1038/nature25966, URL http://www.nature.com/articles/nature25966

Lande R, Schemske DW (1985a) The evolution of self-fertilization and inbreeding depression in plants. i. genetic model. Evolution 39(1):24

Lande R, Schemske DW (1985b) The Evolution of Self-Fertilization and Inbreeding Depression in Plants. I. Genetic Models. Evolution 39(1):24, DOI 10.2307/2408514, URL https://www.jstor.org/stable/2408514?origin=crossref

Larièpe A, Mangin B, Jasson S, Combes V, Dumas F, Jamin P, Lariagon C, Jolivot D, Madur D, Fiévet J, Gallais A, Dubreuil P, Charcosset A, Moreau L (2012) The Genetic Basis of Heterosis: Multiparental Quantitative Trait Loci Mapping Reveals Contrasted Levels of Apparent Overdominance Among Traits of Agronomical Interest in Maize (*Zea mays* L.). Genetics 190(2):795–811, DOI 10.1534/genetics.111.133447, URL https://academic.oup.com/genetics/article/190/2/795/6064102

Lewontin RC (1974) The Genetic Basis of Evolutionary Change. Columbia University Press

Llaurens V, Gonthier L, Billiard S (2009) The Sheltered Genetic Load Linked to the *S* Locus in Plants: New Insights From Theoretical and Empirical Approaches in Sporophytic Self-Incompatibility. Genetics 183(3):1105–1118, DOI 10.1534/genetics.109.102707, URL https://academic.oup.com/genetics/article/183/3/1105/6063166

Mable BK (2008) Genetic causes and consequences of the breakdown of self-incompatibility: case studies in the Brassicaceae. Genetics Research 90(1):47–60, DOI 10.1017/S0016672307008907

McMullen MD, Kresovich S, Villeda HS, Bradbury P, Li H, Sun Q, Flint-Garcia S, Thornsberry J, Acharya C, Bottoms C, Brown P, Browne C, Eller M, Guill K, Harjes C, Kroon D, Lepak N, Mitchell SE, Peterson B, Pressoir G, Romero S, Rosas MO, Salvo S, Yates H, Hanson M, Jones E, Smith S, Glaubitz JC, Goodman M, Ware D, Holland JB, Buckler ES (2009) Genetic Properties of the Maize Nested Association Mapping Population. Science 325(5941):737–740, DOI 10.1126/science.1174320, URL https://www.science.org/doi/10.1126/science.1174320

Ohta T, Cockerham CC (1974) Detrimental genes with partial selfing and effects on a neutral locus. Genetical Research 23(2):191–200, DOI 10.1017/S0016672300014816

Ohta T, Kimura M (1969) Linkage desequilibrium at steady state determined by random genetic drift and recurrent mutation. Genetics 63(1):229–238, DOI 10.1093/genetics/63.1.229, URL https://academic.oup.com/genetics/article/63/1/229/5989253

Olito C, Ponnikas S, Hansson B, Abbott JK (2022) Consequences of partially recessive deleterious genetic variation for the evolution of inversions suppressing recombination between sex chromosomes. Evolution 76(6):1320–1330, DOI https://doi.org/10.1111/evo.14496, URL https://onlinelibrary.wiley.com/doi/abs/10.1111/evo.14496, https://onlinelibrary.wiley.com/doi/pdf/10.1111/evo.14496

van Oosterhout C, Zulstra WG, van Heuven MK, Brakefield PM (2000) Inbreeding depression and genetic load in laboratory metapopulations of the butterfly *Bicyclus anynana*. Evolution 54(1):218–225, DOI 10.1111/j.0014-3820.2000.tb00022.x, URL https://onlinelibrary.wiley.com/doi/10.1111/j.0014-3820.2000.tb00022.x

Rocheleau G, Lessard S (2000) Stability analysis of the partial selfing selection model. Journal of Mathematical Biology 40(6):541–574, DOI 10.1007/s002850000030, URL https://doi.org/10.1007/s002850000030

Roze D (2015) Effects of interference between selected loci on the mutation load, inbreeding depression, and heterosis. Genetics 201:745–757

Seymour DK, Chae E, Grimm DG, Martín Pizarro C, Habring-Müller A, Vasseur F, Rakitsch B, Borgwardt KM, Koenig D, Weigel D (2016) Genetic architecture of nonadditive inheritance in *Arabidopsis thaliana* hybrids. Proceedings of the National Academy of Sciences 113(46), DOI 10.1073/pnas.1615268113, URL https://pnas.org/doi/full/10.1073/pnas.1615268113

Sianta SA, Peischl S, Moeller DA, Brandvain Y (2021) Genetic load may increase or decrease with selfing depending upon the recombination environment. preprint, Genetics, DOI 10.1101/2021.05.20.445016, URL http://biorxiv.org/lookup/doi/10.1101/2021.05.20.445016

Spigler RB, Theodorou K, Chang SM (2017) Inbreeding depression and drift load in small populations at demographic disequilibrium. Evolution 71(1):81–94, DOI 10.1111/evo.13103

Sved JA (1971) Linkage disequilibrium and homozygosity of chromosome segments in finite populations. Theoretical Population Biology 2(2):125–141, DOI https://doi.org/10.1016/0040-5809(71)90011-6, URL https://www.sciencedirect.com/science/article/pii/0040580971900116

Takebayashi N (2003) Patterns of Variation Within Self-Incompatibility Loci. Molecular Biology and Evolution 20(11):1778–1794, DOI 10.1093/molbev/msg209, URL https://academic.oup.com/mbe/article-lookup/doi/10.1093/molbev/msg209

Tallmon DA, Luikart G, Waples RS (2004) The alluring simplicity and complex reality of genetic rescue. Trends in Ecology & Evolution 19(9):489–496, DOI 10.1016/j.tree.2004.07.003, URL https://www.sciencedirect.com/science/article/pii/S0169534704001843

Uyenoyama M, Holsinger K, Waller D (1993) Ecological and genetic factors directing the evolution of self-fertilization. Oxford Surveys in Evolutionary Biology 9:327 – 381

Uyenoyama MK, Waller DM (1991) Coevolution of self-fertilization and inbreeding depression II. Symmetric overdominance in viability. Theoretical Population Biology 40(1):47–77, DOI 10.1016/0040-5809(91)90046-I, URL https://linkinghub.elsevier.com/retrieve/pii/004058099190046I

Waller DM (2021) Addressing Darwin’s dilemma: Can pseudo-overdominance explain persistent inbreeding depression and load? Evolution 75(4):779–793, DOI 10.1111/evo.14189, URL https://onlinelibrary.wiley.com/doi/10.1111/evo.14189

Whitlock M, Ingvarsson PK, Hatfield T (2000) Local drift load and the heterosis of interconnected populations. Heredity 84:452–457

Wickham H (2016) ggplot2: Elegant Graphics for Data Analysis. Springer-Verlag New York, URL https://ggplot2.tidyverse.org

Willi Y, Griffin P, Van Buskirk J (2013) Drift load in populations of small size and low density. Heredity 110(3):296–302, DOI 10.1038/hdy.2012.86

Winn AA, Elle E, Kalisz S, Cheptou PO, Eckert CG, Goode C, Johnston MO, Moeller DA, Ree RH, Sargent RD, et al (2011) Analysis of inbreeding depression in mixed-mating plants provides evidence for selective interference and stable mixed mating. Evolution 65:3339–3359

Wu Q, Han TS, Chen X, Chen JF, Zou YP, Li ZW, Xu YC, Guo YL (2017) Long-term balancing selection contributes to adaptation in Arabidopsis and its relatives. Genome Biology 18(1):217, DOI 10.1186/s13059-017-1342-8, URL https://doi.org/10.1186/s13059-017-1342-8

Zhao L, Charlesworth B (2016) Resolving the conflict between associative overdominance and background selection. Genetics 203(3):1315–1334, DOI 10.1534/genetics.116.188912, URL https://www.genetics.org/content/203/3/1315, https://www.genetics.org/content/203/3/1315.full.pdf

