## Supplementary Figures for "Conditions for maintaining and eroding pseudo-overdominance and its contribution to inbreeding depression"

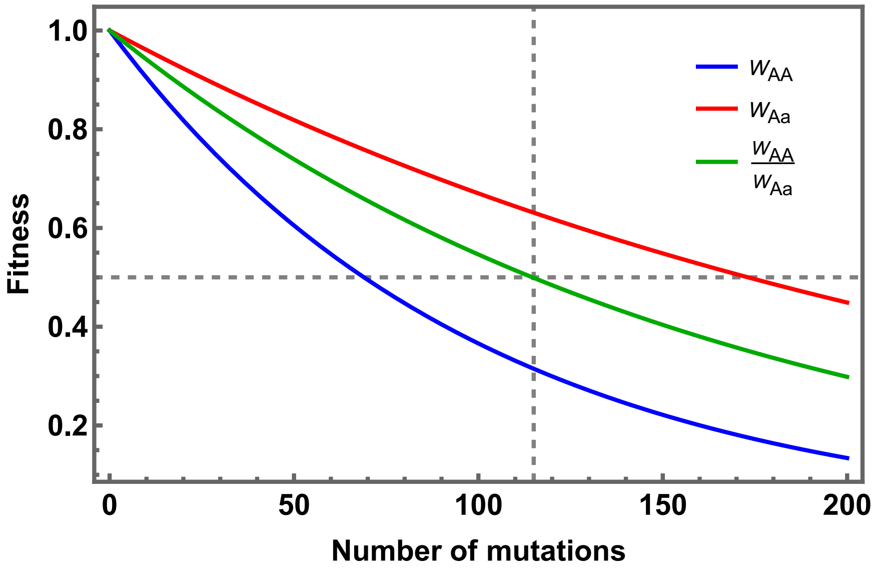


Figure S1: Declines in homozygote (AA) and heterozygote (Aa) fitness and the fitness of homozygotes relative to heterozygotes as a function of the number of completely linked mutations expressing pseudo-overdominance. Mutations are identical, slight ($s = 0.01$), recessive (dominance $h = 0.2$), and have independent (multiplicative) fitness effects. Homozygote fitness is given by $(1-s)^n$ while heterozygote fitness is $(1-hs)^{2n}$ (see Eqs. 2 and 3 in the main text). The horizontal dashed line shows the threshold fitness at which homozygote fitness drops to less than half the fitness of the heterozygote - a condition for POD indefinite persistence even in fully selfing populations. For these parameter values, this occurs for $n = 115$ (the vertical dashed line). (cf. Supp. File 1 Fig. A2).

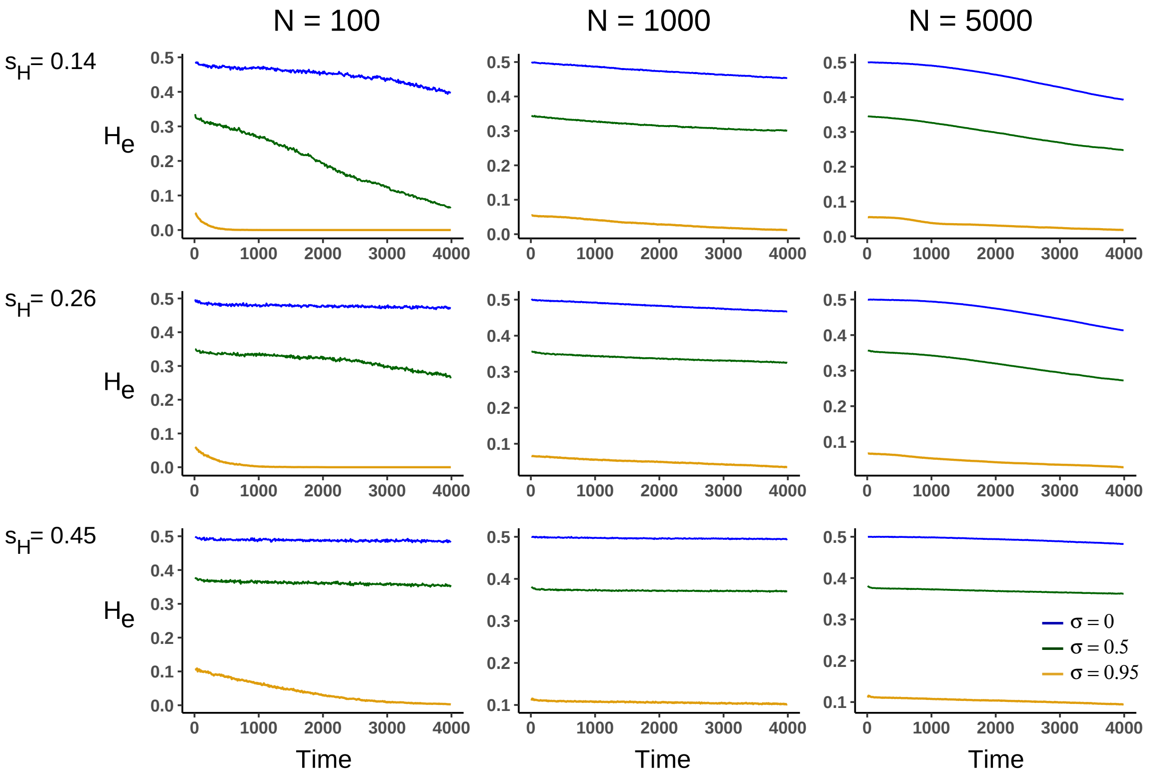


Figure S2: Mean declines in heterozygosity $H_e$ with time for different population sizes, $N$, and selection coefficients against homozygous haplotypes, $s_H$. Simulations reflect three selfing rates ($\sigma = 0, 0.5$ and 0.95). Individual loci within the POD zone have a selection coefficient $s = 0.01$, dominance $h = 0.2$, and map length (or the recombination rate)between loci in the POD zone is $\ell = 10^{-5}$. There are no background mutations.

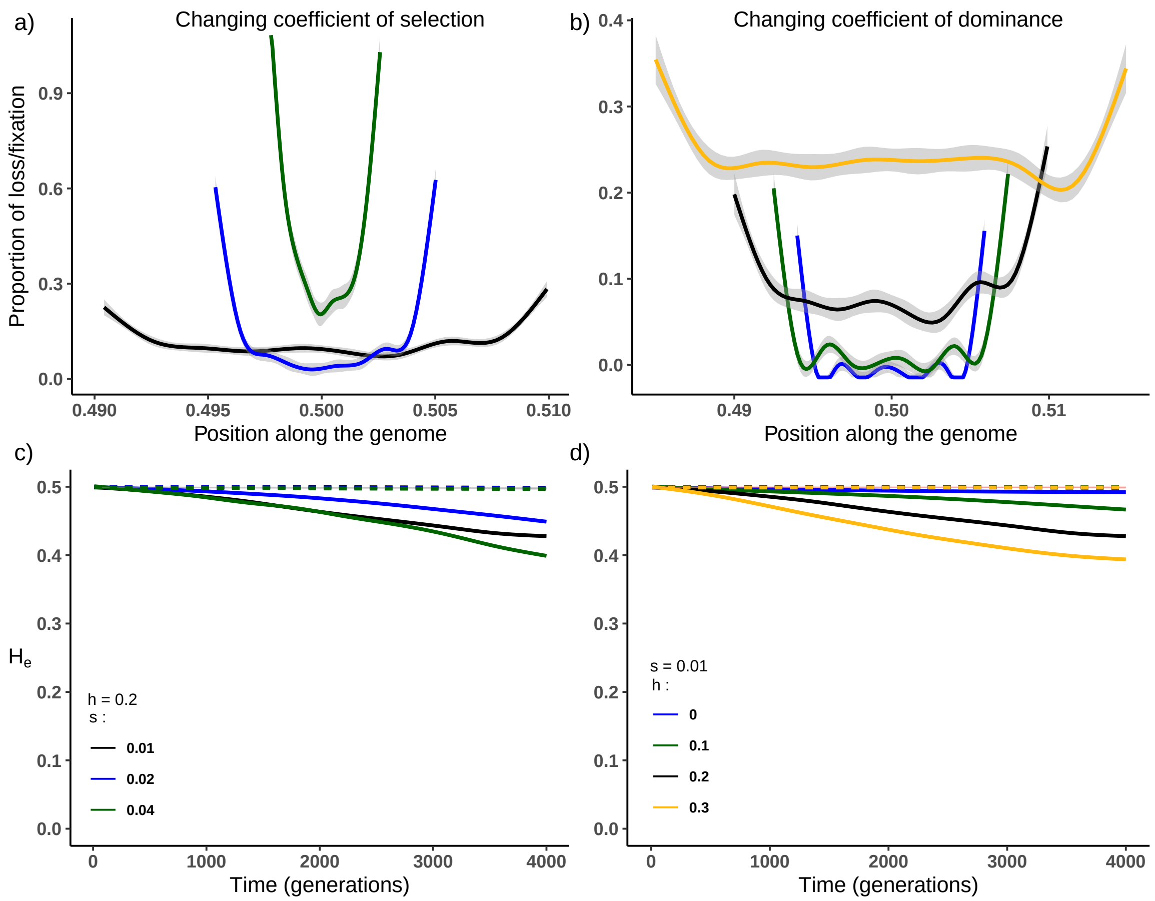


Figure S3: Effects of levels of selection and dominance on selection dynamics within a POD zone (compare to Fig. 2). Left panels show the effects of varying coefficients of selection at each load locus ($s = 0.01, 0.02$ and $0.04$, corresponding to $n_L = 100, 50$ and 25 loci). Dominance is fixed at $h = 0.2$ and $s_H = 0.45$. Right panels show the effects of varying dominance ($h = 0, 0.1, 0.2$ and $0.5$ with $n_L = 60, 75, 100$ and 150) with selection fixed at $s = 0.01$. Panels a) and b) show observed frequencies of fixation/loss along the POD zone (x values represent the position of the loci along the chromosome). The selfing rate $\sigma = 0$ and linkage $\ell = 10^{-4}M$. Panels c) and d) show losses in heterozygosity ($H_e$) over time in populations with a high selfing rate ($\sigma = 0.95$) and either loose linkage ($\ell = 10^{-4}M$, solid lines) or tight linkage ($\ell = 10^{-5}M$, dashed lines).

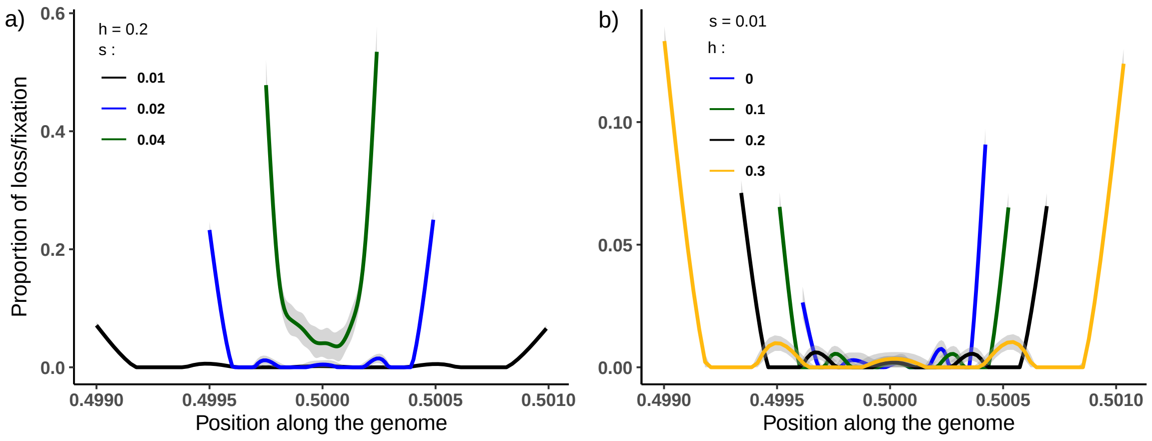


Figure S4: Effects of levels of selection and dominance on selection dynamics within a POD zone. Observed frequencies of fixation/loss along the POD zone (x values represent the position of the loci along the chromosome) after 4000 generations. The selfing rate $\sigma = 0.95$ and linkage $\ell = 10^{-5}M$. a) Effects of varying the coefficient of selection at load loci ($s = 0.01, 0.02$ and $0.04$, corresponding to $n_L = 100, 50$ and 25 loci) with dominance fixed at $h = 0.2$ and $s_H = 0.45$. b) Effects of varying dominance ($h = 0, 0.1, 0.2$ and $0.5$ with $n_L = 60, 75, 100$ and 150) with selection fixed at $s = 0.01$.

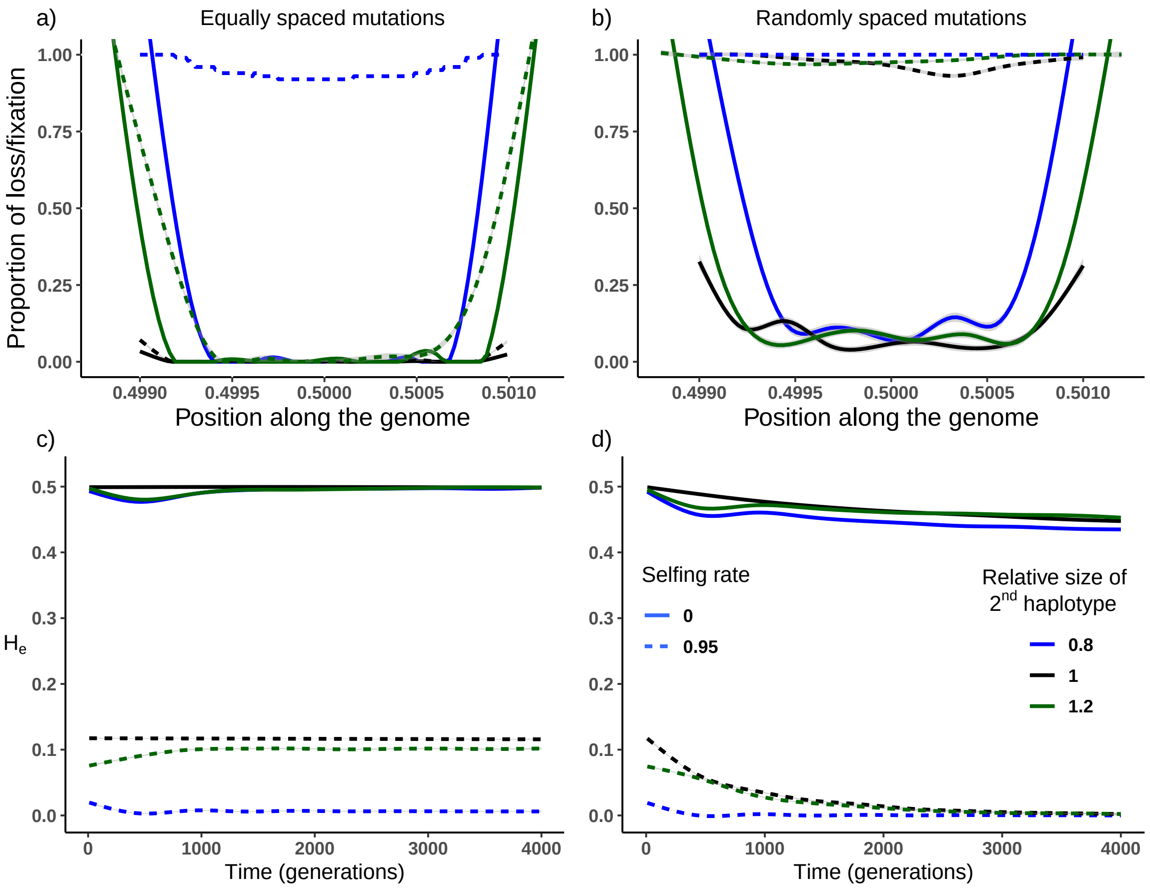


Figure S5: Effects of relaxing the assumptions of symmetric overdominance and evenly spaced mutations. Upper panels show locations within the POD zone where load mutations are most likely to be lost after 4000 generations (a, b) and how this depends on whether mutations are evenly spaced (a) or randomly distributed (b). Results are shown for both symmetric (black) and asymmetric (green and blue) loads. Outcomes under both outcrossing and high selfing (solid vs. dotted lines) are shown. Note erosion of mutations via recombination and selection at both ends of the POD zone. Lower panels show overall stability of the POD zone (shown as heterozygosity, $He$) over time. As in the upper panels, graphs show results for both symmetric (black) and asymmetric (green and blue) loads and for evenly (panel c) and randomly (panel d) placed mutations. The coefficients of selection and dominance are $s = 0.01$ and $h = 0.2$ respectively, linkage within the POD zone is $\ell = 10^{-5}$ and population size $N = 1000$.

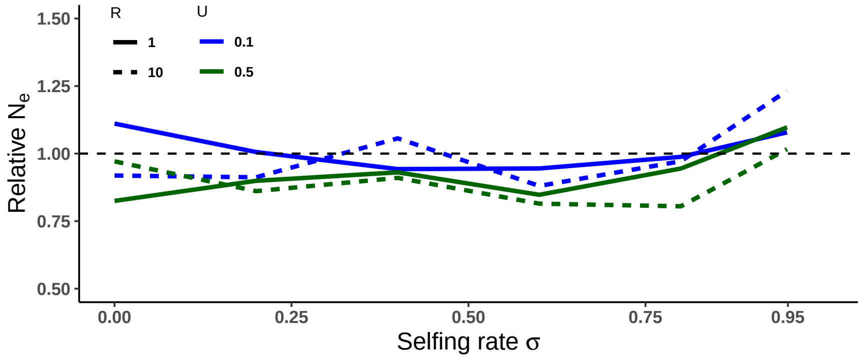


Figure S6: Effective population size, $N_e$, in a populations supporting a POD zone relative to a population without one (done by setting $s = h = 0$). $N_e$ varies as a function of the selfing rate for populations subject to different background mutation rates ($U$) and shorter and longer map lengths ($R$ in Morgans). These simulations use 100 POD load loci ($n_L = 100$) and a map length of $\ell = 10^{-6}$ Morgans. Mutations within the POD zone are randomly placed. Selection coefficients in- and outside the POD zone ($s$ and $s_d$ respectively) are 0.01 with dominances $h$ and $h_d = 0.2$.

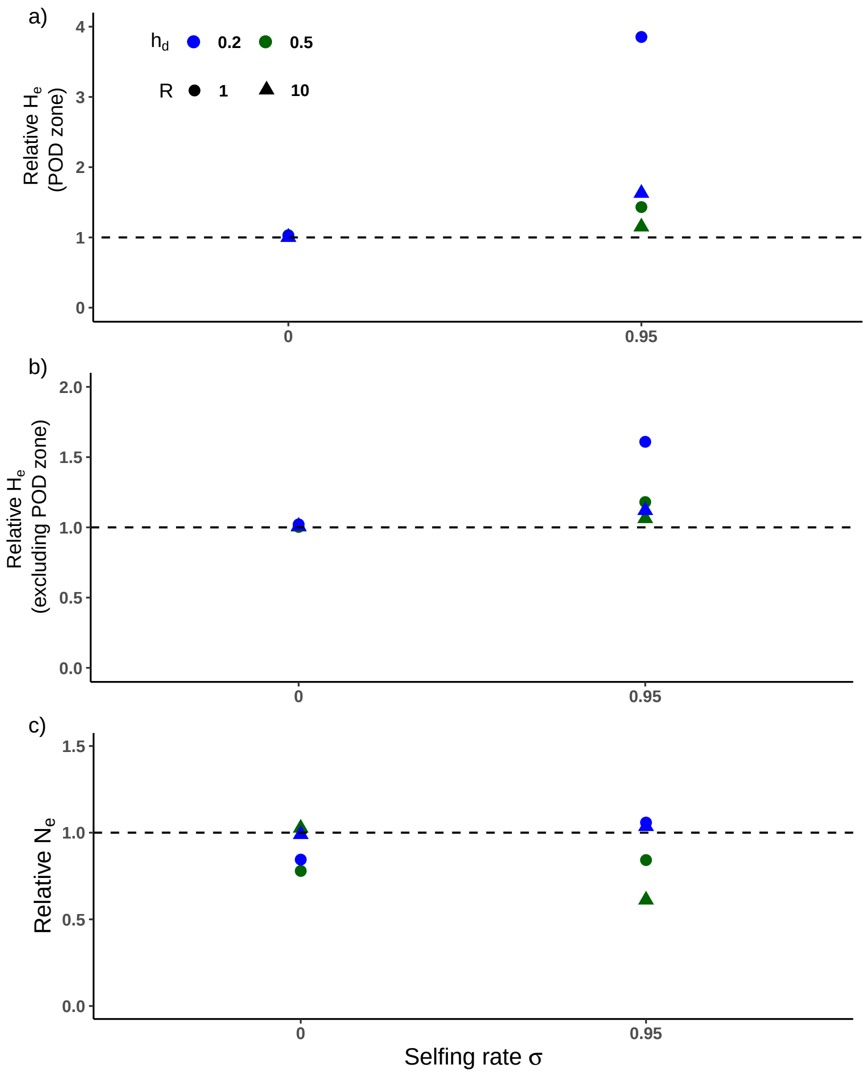


Figure S7: Associative-Overdominance of background mutations affects selection dynamics within POD zones and vice versa. Graphs compare the effect of recessive ($h_d = 0.2$ blue) to codominant ($h_d = 0.5$ green) background mutations on population heterozygosity as a function of the selfing rate. a) Heterozygosity, $H_e$, within a POD zone with background mutations occurring elsewhere in the genome relative to $H_e$ in a population lacking background mutations. b) $H_e$ outside the POD zone in a population with a POD zone relative to one without. c) $N_e$ in a population with POD selection relative to one without. The background mutation rate $U = 0.5$ and the map lengths simulated are $R = 1$ and 10 Morgans. These simulations use 100 POD load loci ($n_L = 100$) randomly placed within the POD zone over a map length of $\ell = 10^{-6}$ Morgans. Selection coefficients within the POD zone are $s = 0.01$with dominance $h = 0.2$ and population size $N = 1000$.

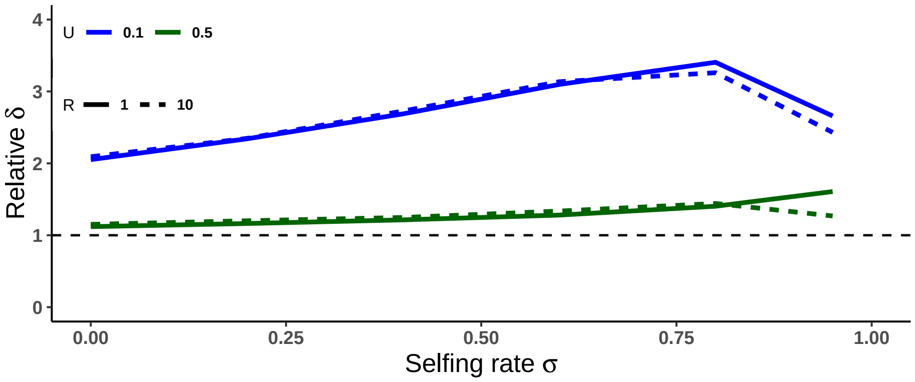


Figure S8: Inbreeding depression in a population with a POD zone relative to one without as a function of the selfing rate $\sigma$ at two genomic mutations rates, $U$, and two genomic map lengths, $R$. Other parameter values are $n_L = 100$ and $\ell = 10^{-6}$ Morgans with mutations randomly placed within the POD zone. Selection coefficients within and outside the POD zone, $s$ and $s_d$ respectively are set to 0.01, and dominances $h$ and $h_d = 0.2$.

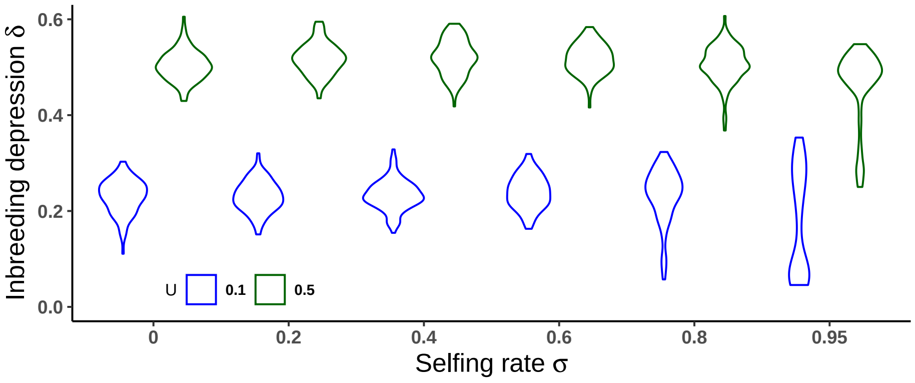


Figure S9: Distributions of inbreeding depression, $\delta$, as a function of the population selfing rate in populations with a higher ($U = 0.5$) or lower ($U = 0.1$) haploid genomic mutation rate. Other parameter values: $n_L = 100$, $\ell = 10^{-6}$ Morgans with mutations randomly placedin the POD zone. Selection coefficients within and outside the POD zone, $s$ and $s_d$ respectively are set to 0.01, and dominances $h$ and $h_d = 0.2$. The genome size (setting the recombination rate) is $R = 10$ Morgans.

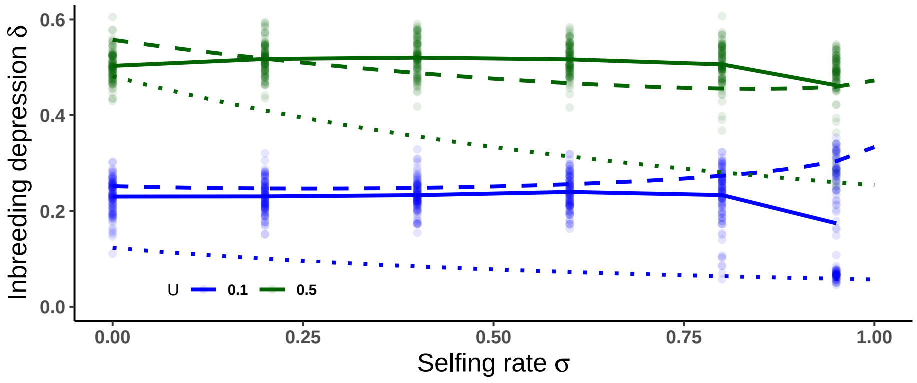


Figure S10: POD zones increase inbreeding depression. Inbreeding depression, $\delta$, as a function of the selfing rate for populations with higher ($U = 0.5$) and lower ($U = 0.1$) haploid genomic mutation rates. Solid lines represent means of the simulations run with a POD zone while dotted lines show the $\delta$ expected in the absense of overdominance (Eq. (7)). Dashed lines show the $\delta$ expected with overdominant selection (Eq. (6)). Other parameter values are $n_L = 100$ and $\ell = 10^{-6}$ Morgans. Mutations are randomly placed within the POD zone. Selection coefficients within and outside the POD zone, $s$ and $s_d$ respectively are set to 0.01, and dominances $h$ and $h_d = 0.2$. The genome size setting the recombination rate is $R = 1$ Morgans.
