## Supplementary File 1 for "Conditions for maintaining and eroding pseudo-overdominance and its contribution to inbreeding depression"

### Overdominant selection and inbreeding

#### A1 Genetic frequencies at equilibrium

Here we give the full terms used to obtain the equations in the main text. From Appendix A4 from Kimura and Ohta (1971), we have a population with a single bi-allelic overdominant locus, carrying alleles $A$ and $B$. The equilibrium frequencies of the three genotype ($P_{11}$, $P_{12}$ and $P_{22}$ for $AA$, $AB$ and $BB$) are:

$$\begin{aligned} P_{11} &= \frac{xi(1-\hat{F})}{s_1} \\ P_{12} &= 1 - \frac{(1+\hat{F})i(s_1(1-x)+s_2x)}{s_1s_2} \\ P_{22} &= \frac{(1-xi)(1-\hat{F})}{s_2} \end{aligned} \tag{A1}$$

with $s_1$ and $s_2$ the coefficients of selection associated with $P_{11}$ and $P_{22}$ respectively. As mentioned in the main text $x$, the frequency of allele $A$, and $i$ are:

$$x = \frac{s_2 - s_1\hat{F}}{(s_1+s_2)(1-\hat{F})}$$

and (A2)

$$i = \frac{s_1s_2}{s_1+s_2}$$

#### A2 Coefficient of inbreeding $\hat{F}$

The coefficient of inbreeding for overdominant selection $\hat{F}$ is expressed in terms of the coefficients of selection and the rate of self-fertilisation $\sigma$:

$$\hat{F} = \frac{2(1-i)-(1-2i)\sigma-\sqrt{D}}{4i}$$

with (A3)

$$D = 4(1-i)^2 + (1-2i)^2\sigma^2 - 4(1+i)(1-2i)\sigma$$

As shown in Figure A1, the smaller the selection against homozygotes, the more $\hat{F}$ becomes equivalent to the classically used definition of the coefficient of inbreeding, which only depends on the rate of self-fertilization $\sigma$, $F = \frac{\sigma}{2-\sigma}$ (see Glémin, 2021).

However, very strong selection against homozygotes completely cancels the effect of self-fertilization on genetic frequencies, with $\hat{F}$ tending to zero for all values of $\sigma$.

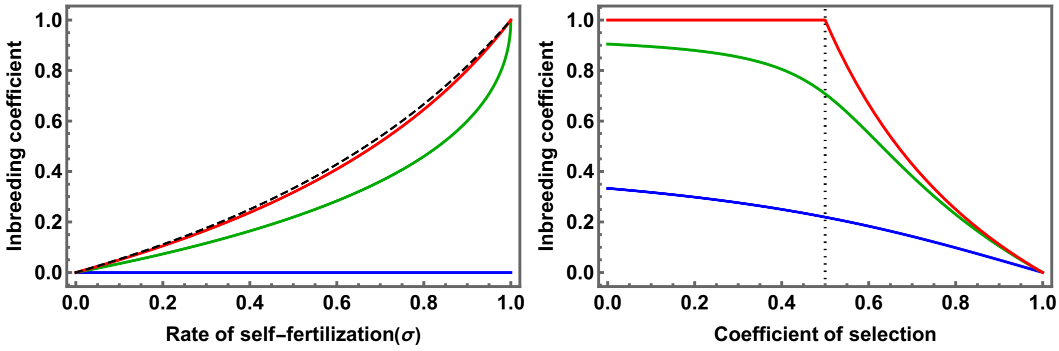


Figure A1: The effect of self-fertilisation and selection on the value of $\hat{F}$ (equation A3). In the left panel, we compare the coefficient of inbreeding $F$ (black, dashed line) to $\hat{F}$ for $s_1 = s_2 = 0.1$ (red line), 0.5 (green line) and 1 (lethal homozygotes, blue line). In the right panel we show the value of $\hat{F}$ as a function of the coefficient of selection $s_H$ (i.e considering $s_1 = s_2$ out of convenience) for three values of $\sigma$ (0.5, 0.95 and 1, in blue, green and red, respectively). The black dotted line is set at $s_H = 0.5$ to highlight the threshold above which overdominance is stable for any value of $\sigma$.

#### A3 Stability of polymorphism with overdominance

From Equation 1 in the main text, we can plot the conditions necessary, with regards to the values of $s_1$, $s_2$ and $\sigma$ for which there is a stable polymorphic equilibrium (*i.e.* $0 > x < 1$, see also Figure A.1 in Kimura and Ohta (1971), Appendix 4). From this figure, it is clear that the maintenance of a stable equilibrium is more constrained the higher the rate of self-fertilisation.

#### A4 Inbreeding depression $\delta_{od}$ (overdominant selection)

The expected level of inbreeding depression ($\delta$) depends on the fitnesses of selfed and outcrossed offspring, respectively $W_s$ and $W_o$. These variables can be expressed in

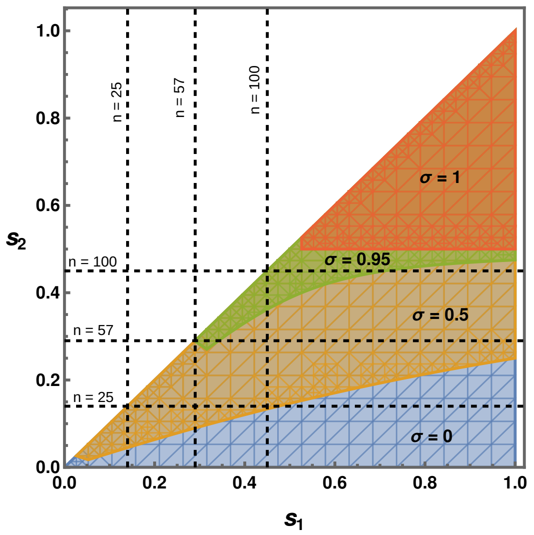


Figure A2: Representing the combinations of $s_1$ and $s_2$ (assuming $s_1 > s_2$) for which the inequality presented in Equation 1 from the main text is true (the regions above and to the right of the lines) for different values of $\sigma$ (0, 0.5, 0.95 and 1, in blue, orange, green and red, respectively). The dotted lines represent the coefficients of selection based on the number of loci carrying mutations with a coefficient of selection $s = 0.01$ and dominance $h = 0.2$ (see Eq. 3 in the main text).

terms of the coefficients of selection associated with each homozygote and the expected genotypic frequencies (see Charlesworth and Charlesworth 1990):

$$\begin{aligned} W_s &= (1 - s_1)(P_{11} + \frac{P_{12}}{4}) + (1 - s_2)(P_{22} + \frac{P_{12}}{4}) + \frac{P_{12}}{2} \\ W_o &= 1 - s_1 x(P_{11} + \frac{P_{12}}{2}) - s_2(1 - x)(P_{22} + \frac{P_{12}}{2}) \end{aligned} \tag{A4}$$

Using the expressions for the genotypic frequencies given in Eq. A1and the definitions in Eq. A2, and setting $s_1 = s_2$, we obtain the simplified expression for $\delta$, given in Eq. 9 of the main text.
