## Supplementary File 2 for "Conditions for maintaining and eroding pseudo-overdominance and its contribution to inbreeding depression"

### Selection for recombinant haplotypes

#### Coefficients of selection associated with recombinants

Any recombinant haplotype $H_c$ will have a heterozygote advantage that depends on the haplotype with which it is inherited ($H_1$ or $H_2$). If we assume that no mutation has been cleaved during recombination, and that the fitness of reference is that of the initial heterozygote $H_1H_2$ (as in Eq. 3 in the main text) then the coefficient of selection against the homozygote $H_cH_c$ is simply $s_c = s_H$. The coefficients of selection against heterozygotes $H_cH_1$ and $H_cH_2$ are given by:

$$\begin{aligned} s_{c,1} &= 1 - \frac{(1-hs)^{(2-b)n}(1-s)^{nb/2}}{(1-hs)^{2n}} \\ s_{c,2} &= 1 - \frac{(1-hs)^{bn}(1-s)^{n(2-b)/2}}{(1-hs)^{2n}} \end{aligned} \qquad \text{(B1)}$$

Here $n$ is the number of loci carrying deleterious mutations. The parameter $b$ ($0 \leq b \leq 2$) reflects the similarity between the recombinant haplotype $H_c$ and (arbitrarily) haplotype $H_1$. When $b \approx 0$ (respectively 2), this implies that the recombinant is made up mostly of the haplotype $H_2$ (respectively $H_1$), giving $s_{c,1} = 0$ (respectively $s_{c,1} = s_H$). If $b = 1$ then $H_c$ is made up of equal parts of $H_1$ and $H_2$ giving $s_{c,1} = s_{c,2}$.

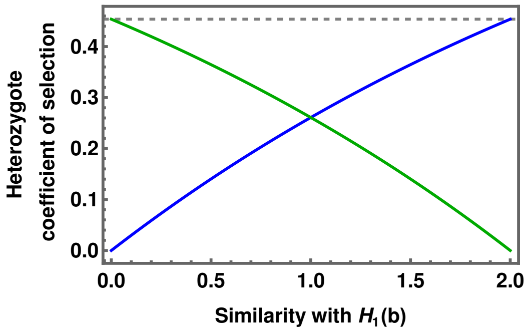


Figure B1: The relative values of the coefficients of selection against heterozygotes $H_cH_1$ and $H_cH_2$, respectively $s_{c,1}$ (blue line) and $s_{c,2}$ (green line) for different values of parameter $b$ (See Eq. B1. The dotted line represents $s_H$, the coefficient of selection against homozygotes.

In the case of a loss of a single mutation during recombination, arbitrarily initially present on $H_1$, then the coefficients of selection involving $H_c$ become:

$$\begin{aligned} s_c &= 1 - \frac{(1-s)^{n-1}}{(1-hs)^{2n}} \\ s_{c,1} &= 1 - \frac{(1-hs)(1-s)^{n-1}}{(1-hs)^{2n}} \\ s_{c,2} &= 1 - \frac{(1-hs)^{n-1}}{(1-hs)^{2n}}. \end{aligned} \tag{B2}$$

#### Changes in frequencies and equilibrium expectations

Previous works have shown that several overdominant haplotypes can co-exist when heterozygotes all have the same relative fitness of 1 (see for example Glémin, 2021). From the expressions for the coefficients of selection in Eqs. B1 and B2it is clear that this assumption cannot be made in the context of POD. To evaluate whether more than two haplotypes can exist, we derive expressions for the changes in frequencies $P_1$, $P_2$ and $P_c$ of haplotypes $H_1$, $H_2$ and $H_c$ respectively:

$$\begin{aligned} \Delta_{P_1} &= \frac{P_1((1-s_1)((1-F_i)P_1+F_i)+(1-F_i)P_2+(1-F_i)P_c(1-s_{c,1}))}{\overline{W}} - P_1 \\ \Delta_{P_2} &= \frac{P_2((1-F_i)P_1+(1-s_2)((1-F_i)P_2+F_i)+(1-F_i)P_c(1-s_{c,2}))}{\overline{W}} - P_2 \\ \Delta_{P_c} &= \frac{P_c((1-F_i)P_1(1-s_{c,1})+(1-F_i)P_2(1-s_{c,2})+(1-s_c)((1-F_i)P_c+F_i))}{\overline{W}} - P_c. \end{aligned} \tag{B3}$$

where $\overline{W}$ is the mean expected fitness, given by:

$$\begin{aligned} \overline{W} &= (1-s_1)\left((1-F_i)P_1^2+F_iP_1\right)+2(1-F_i)P_1P_2+2(1-F_i)P_1P_c(1-s_{c,1}) \\ &+(1-s_2)\left((1-F_i)P_2^2+F_iP_2\right)+2(1-F_i)P_2P_c(1-s_{c,2})+(1-s_c)\left((1-F_i)P_c^2+F_iP_c\right) \end{aligned} \tag{B4}$$

and $F_i$ is a generic coefficient of inbreeding, which may not necessarily be equal to $\hat{F}$ (see Eq. A3 in Supp. File 1) in a context of overdominance with three alleles or haplotypes. Setting $s_1 = s_2$ and using Wolfram Mathematica (REF) to resolve the system of Ordinary Differential Equations above, we find that the only possible equilibria would be for either the fixation of $H_1$, $H_2$ or $H_c$, or the loss one of the haplotypes.

As shown in the main text (Eq. 12), by assuming $P_c$ is of order $\epsilon$ ($\epsilon$ being very close to zero), we can determine whether a rare recombinant $H_c$ can increase in frequency in a population at equilibrium for the two initial haplotypes. We derive the expression for $\Delta_{P_c}$ to the first order of $\epsilon$ with $s_H = s_1, s_2$ and $P_1 = P_2 = (1-\epsilon)/2$:

$$\bar{\Delta}_{P_c} = \frac{2((1+F)s_H - s_{c,1} - s_{c,2} - F(2s_c - s_{c,1} - s_{c,2}))}{2 - s_H - Fs_H}. \tag{B5}$$

Deriving the above expression while considering $F = \hat{F}$ gives an expression for $\Delta_{P_c}$ that depends on the initial coefficient of selection of the recombinant haplotype $s_c$ :

$$\frac{2s_H^2(1+\sigma) - 2(2s_c - s_{c,1} - s_{c,2})(2 - X - \sigma) + 2s_H(2 - 3s_{c,1} - 3s_{c,2} - X + 2s_c(1-\sigma) - \sigma + (s_{c,1} + s_{c,2})\sigma)}{s_H(2 + X + \sigma - s_H(1+\sigma))} \tag{B6}$$

with $X = \sqrt{(2-s_H)^2 - 2\left(2 - s_H - s_H^2\right)\sigma + (1-s_H)^2\sigma^2}$

If $s_c = s_H$, the same conditions as presented in the two-haplotype case (see Supp. File 1) need to be filled for $H_c$ to replace either of the initial haplotypes (*i.e.* $H_c$ must be similar enough to either $H_1$ or $H_2$ to replace the genotype in question). This is the case when $b \approx 0$ or 2 (see Equation B1 and Fig. B2). As $\Delta_{p_c} \approx 0$, $H_c$ is neither selected for or against, and drift should play a major role in determining the frequency $P_c$. For $0 < b < 2$ the recombinant will always be selected against ($\Delta_{P_c} < 0$).

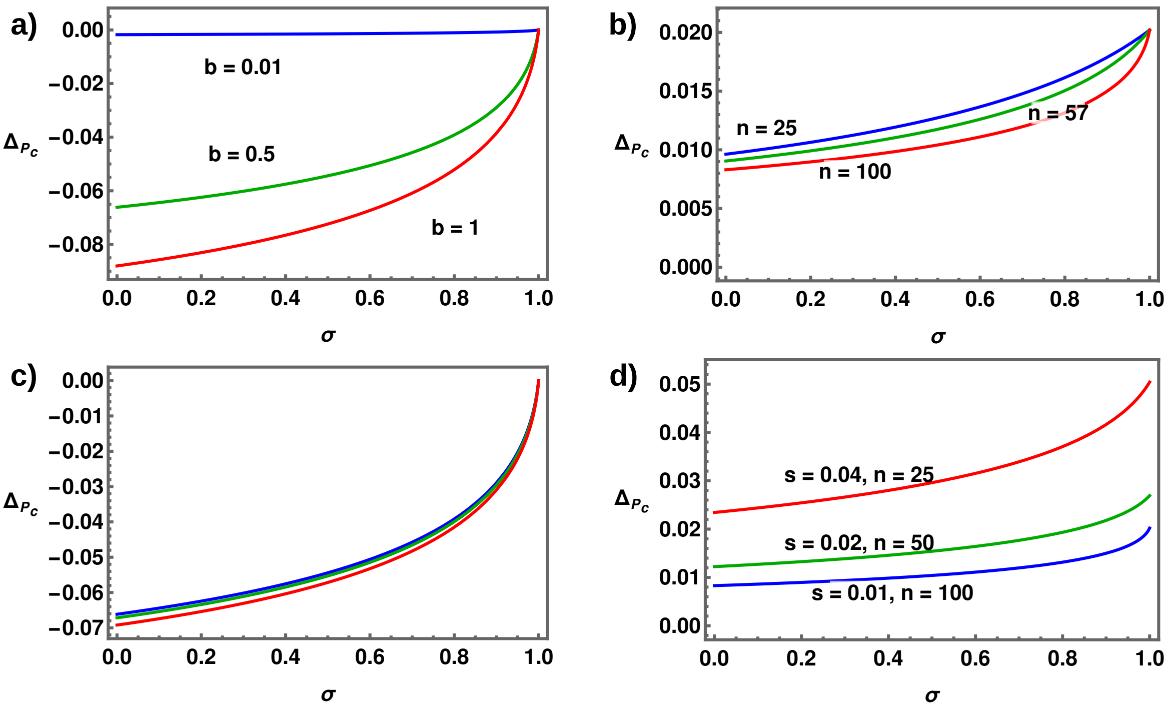


Figure B2: The change in frequency of a haplotype $H_c$ ($\bar{\Delta}_{P-c}$, see Eq. B5) as a function of the selfing rate $\sigma$. a) and c) No mutations are lost and $n = 100$, b) and d) a single mutation is cleaved. a) Each coloured line represents a different value of the parameter $b$ (see Eq. B1 and accompanying text) for $n = 100$. b) Each coloured line represents a different value of $n$. For both a) and b) $s = 0.01$ and $h = 0.2$. c) and d) show values of $\bar{\Delta}_{P_c}$ for the same $s_H$, but for different $s$ and $n$ (but fixed $h = 0.2$).
