## Supplementary material for "Conditions for maintaining and eroding pseudo-overdominance and its contribution to inbreeding depression": R script to generate figures: POD_inbreeding_script_Figures.R

```
library(ggplot2)
library(magrittr)
library(reshape2)
library(gridExtra)
require(lattice)
library(ggpubr)
library(dplyr)
library(plotly)
library(cowplot)
library(latex2exp)
library(scales)


## Fig 2 - Ideal scenario
# Theoretical expectations DeltaPc
# Different values of s
datah <- data.frame(deltap = c(0.00830169, 0.00845452, 0.00861656, 0.00878881, 0.00897245, 0.00916887,  0.00937973, 0.00960702, 0.00985319, 0.0101212, 0.0104149, 0.0107392, 0.0111003, 0.0115071, 0.0119716, 0.0125118, 0.0131557, 0.0139517, 0.0149959, 0.0165375, 0.020202), a = seq(0,1,0.05), s = 0.01)
sim1 <- data.frame(deltap = c(0.0167039, 0.0170117, 0.017338, 0.0176849, 0.0180547, 0.0184502, 0.0188749, 0.0193326, 0.0198283, 0.020368, 0.0209595, 0.0216124, 0.0223398, 0.023159, 0.0240945, 0.0251826, 0.0264799, 0.0280843, 0.0301901, 0.0333036, 0.0408163), a = seq(0,1,0.05), s = 0.02)
datah = rbind(datah, sim1)
sim1 <- data.frame(deltap = c(0.0338183, 0.0344425, 0.0351043, 0.0358077, 0.0365576, 0.0373596, 0.0382206, 0.0391487, 0.0401538, 0.0412482, 0.0424475, 0.0437714, 0.0452463, 0.0469076, 0.0488052, 0.0510126, 0.0536454, 0.0569032, 0.0611841, 0.0675313, 0.0833333), a = seq(0,1,0.05), s = 0.04)
datah = rbind(datah, sim1)


deltap_s <- ggplot(data = datah, aes(x = a, y = deltap, colour = as.factor(s)))+geom_line()+
  theme(text = element_text(size = 12, face = "bold"),legend.position=c(0.1,0.7),
        panel.background=element_blank(),axis.line = element_line(colour = "black"),axis.title.y = element_text(angle=0), axis.title = element_text(size = 14))+ theme(strip.background = element_blank(),strip.text.x = element_blank(), axis.title.y = element_text(vjust=0.5), plot.title = element_text(hjust=0.5, size = 12, vjust = -5),legend.title = element_text(size = 9), legend.text = element_text(size = 8))+ 
  ylab(TeX("$\\Delta_{pc}$"))+
  xlab(TeX("Selfing rate $\\sigma$"))+
  labs(title = TeX("Changing coefficient of selection"), color = TeX("h = 0.2\n s"))+
  scale_color_manual(breaks = c("0.01", "0.02", "0.04"), values=c("black", "blue", "darkgreen"))

deltap_s


# Different values of h
datah <- data.frame(deltap = c(0.00830169, 0.00845452, 0.00861656, 0.00878881, 0.00897245, 0.00916887,  0.00937973, 0.00960702, 0.00985319, 0.0101212, 0.0104149, 0.0107392, 0.0111003, 0.0115071, 0.0119716, 0.0125118, 0.0131557, 0.0139517, 0.0149959, 0.0165375, 0.020202), a = seq(0,1,0.05), h = 0.2)
sim1 <- data.frame(deltap = c(0.00714451, 0.00731255, 0.00749071, 0.00768011, 0.00788203, 0.008098, 0.00832986, 0.00857978, 0.00885045, 0.00914518, 0.00946813, 0.00982465, 0.0102218, 0.010669, 0.0111797, 0.0117736, 0.0124815, 0.0133565, 0.0145041, 0.0161974, 0.020202), a = seq(0,1,0.05), h = 0)
datah = rbind(datah, sim1)
sim1 <- data.frame(deltap = c(0.00772153, 0.007882, 0.00805214, 0.00823301, 0.00842584,0.00863209, 0.0088535, 0.00909216, 0.00935064, 0.00963209, 0.0099405, 0.010281, 0.0106602, 0.0110873, 0.011575, 0.0121422, 0.0128182, 0.013654, 0.0147501, 0.0163678, 0.020202), a = seq(0,1,0.05), h = 0.1)
datah = rbind(datah, sim1)
sim1 <- data.frame(deltap = c(0.00888501, 0.00903005, 0.00918383, 0.0093473, 0.00952159,0.009708, 0.00990811, 0.0101238, 0.0103574, 0.0106118, 0.0108906, 0.0111983, 0.011541, 0.0119271,0.0123679, 0.0128806, 0.0134918, 0.0142475, 0.015239, 0.0167035, 0.020202), a = seq(0,1,0.05), h = 0.3)
datah = rbind(datah, sim1)


deltap_h <- ggplot(data = datah, aes(x = a, y = deltap, colour = as.factor(h)))+geom_line()+
  theme(text = element_text(size = 12, face = "bold"),legend.position=c(0.1,0.7),
        panel.background=element_blank(),axis.line = element_line(colour = "black"),axis.title.y = element_text(angle=0), axis.title = element_text(size = 14))+ theme(strip.background = element_blank(),strip.text.x = element_blank(), axis.title.y = element_text(vjust=0.5), plot.title = element_text(hjust=0.5, size = 12, vjust = -5),legend.title = element_text(size = 9), legend.text = element_text(size = 8))+ 
  ylab(TeX(""))+
  xlab(TeX("Selfing rate $\\sigma$"))+
  labs(title = TeX("Changing coefficient of dominance"), color = TeX("s = 0.01\n h"))+
  scale_color_manual(breaks = c("0", "0.1", "0.2", "0.3"), values=c("blue", "darkgreen","black", "darkgoldenrod1"))

deltap_h

#Change in He selfing = 0.95
# diff values of s
stable = c(0.01, 0.02, 0.04)
ntable = c(100, 50, 25)
linktable = c(1e-05, "0.0001")
atable = c(0)
dHtot = data.frame();
for (m in 1:1){
  for (l in 1:2){
    for (k in 1:3){
      test<- read.table(paste("N1000_a",atable[m],"_r0_nL",ntable[k],"_nEq1_link",linktable[l],"_sP",stable[k],"_hP0.2_U0_L1.txt",sep =""), sep = " ", fill = TRUE, header = FALSE )
      
      dcal <- ifelse(test[4:403,13] == 0, 0, test[4:403,12]/test[4:403,13])
      
      dH <- data.frame( s = stable[k], gen = test[4:403,2], He = dcal, link = linktable[l])
      for(i in 1:99){
        j = i*403 + 3;
        dcal <- ifelse(test[(j+1):(j+400),13] == 0, 0, test[(j+1):(j+400),12]/test[(j+1):(j+400),13])
        
        sim1 <-data.frame(s = stable[k], gen = test[(j+1):(j+400),2], He = dcal, link = linktable[l])
        dH = rbind(dH, sim1);
      }
      dHtot = rbind(dH, dHtot)
    }
  }
}


dHe_s<-ggplot(data = dHtot)+
  geom_smooth(mapping =aes(x = gen, y = He, colour=as.factor(s), linetype = as.factor(link)))+
  coord_cartesian(ylim = c(0, 0.5), xlim = c(0,4000)) +
  theme(text = element_text(size = 12, face = "bold"), legend.position="none",
        panel.background=element_blank(),axis.line = element_line(colour = "black"))+ theme(strip.background = element_blank(),strip.text.x = element_blank(),axis.title.y = element_text(vjust=0.5, angle = 0))+ 
  ylab(TeX("$H_e$"))+
  xlab(TeX("Time (generations)"))+
  scale_color_manual(breaks = c("0.01", "0.02", "0.04"), values=c("black","blue", "darkgreen"))

dHe_s

# diff values of h
# Diff h
htable = c(0, 0.1, 0.2, 0.3)
ntable = c(60, 75, 100, 150)
linktable = c(1e-05, "0.0001")
atable = c(0)
dHtot = data.frame();
for (m in 1:1){
  for (l in 1:2){
    for (k in 1:4){
      test<- read.table(paste("N1000_a",atable[m],"_r0_nL",ntable[k],"_nEq1_link",linktable[l],"_sP0.01_hP",htable[k],"_U0_L1.txt",sep =""), sep = " ", fill = TRUE, header = FALSE )
      dcal <- ifelse(test[4:403,13] == 0, 0, test[4:403,12]/test[4:403,13])
      dH <- data.frame(h = htable[k], gen = test[4:403,2], He = dcal, link = linktable[l], a = atable[m])
      for(i in 1:99){
        j = i*403 + 3;
        dcal <- ifelse(test[(j+1):(j+400),13] == 0, 0, test[(j+1):(j+400),12]/test[(j+1):(j+400),13])
        sim1 <-data.frame(h = htable[k], gen = test[(j+1):(j+400),2], He = dcal, link = linktable[l], a = atable[m])
        dH = rbind(dH, sim1);
      }
      
      dHtot = rbind(dH, dHtot)
    }
  }
}


dHe_h<-ggplot(data = dHtot)+
  geom_smooth(mapping =aes(x = gen, y = He, colour=as.factor(h), linetype = as.factor(link), fill = as.factor(a) ))+
  coord_cartesian(ylim = c(0, 0.5), xlim = c(0,4000)) +
  theme(text = element_text(size = 12, face = "bold"),legend.position="none",
        panel.background=element_blank(),axis.line = element_line(colour = "black"))+ theme(strip.background = element_blank(),strip.text.x = element_blank(), axis.title.y = element_text(vjust=0.5, angle = 0))+ 
  ylab(TeX(""))+
  xlab(TeX("Time (generations)"))+
  scale_color_manual(breaks = c("0.2", "0", "0.1", "0.3"), values=c("black","blue", "darkgreen", "darkgoldenrod1"))
dHe_h

## Fixation patterns

# diff values of s

stable = c(0.01, 0.02, 0.04)
ntable = c(100, 50, 25)
linktable = c("0.0001")
atable = c(0)
dHtot = data.frame();
for (m in 1:1){
  for (l in 1:1){
    for (k in 1:3){
      test<- read.table(paste("N1000_a",atable[m],"_r0_nL",ntable[k],"_nEq1_link",linktable[l],"_sP",stable[k],"_hP0.2_U0_L1_Haplo.txt",sep =""), sep = " ", fill = TRUE, header = FALSE )
      
      sim1 <-data.frame(pos = melt(as.list(test[1,2:(2*ntable[k]+1)]))[1], state= melt(as.list(test[3,2:(2*ntable[k]+1)]))[1] )
      colnames(sim1) = c("pos", "state")
      sim1 <- sim1 %>% mutate(freq = if_else(state == 0, 1, if_else(state == 2,1, 0)))
      sim1 <- sim1 %>% mutate(s = stable[k])
      datah = sim1;
      
      
      for(i in 1:99){
        sim1 <-data.frame(pos = melt(as.list(test[3*i+1,2:(2*ntable[k]+1)]))[1], state= melt(as.list(test[3*i+3,2:(2*ntable[k]+1)]))[1] ) 
        colnames(sim1) = c("pos", "state")
        sim1 <- sim1 %>% mutate(s = stable[k])
        sim1 <- sim1 %>% mutate(freq = if_else(state == 0, 1, if_else(state == 2,1, 0)))
        sim1 =sim1[order(sim1$pos),];
        datah = rbind(datah, sim1);
      }
      dHtot = rbind(datah, dHtot)
    }
  }
}

dfix_s<- ggplot(dHtot) +
  geom_smooth(mapping = aes(x=pos, y=freq,colour=as.factor(s)))+
  scale_y_continuous(limit=c(0,NA),oob=squish)+
  theme(text = element_text(size = 12, face = "bold"),legend.position="none",
        panel.background=element_blank(),axis.line = element_line(colour = "black"))+ theme(strip.background = element_blank())+ 
  ylab(TeX("Proportion of loss/fixation"))+
  xlab(TeX("Position along the genome"))+
  scale_color_manual(breaks = c("0.01", "0.02", "0.04"), values=c("black","blue", "darkgreen"))

dfix_s

#ggplot(data = dHtot, aes(x = pos, y = freq, colour=as.factor(s))) + stat_summary(geom = "line", fun = "mean")


# diff values of h
htable = c(0, 0.1, 0.2, 0.3)
ntable = c(60, 75, 100, 150)
linktable = c("0.0001")
atable = c(0)
dHtot = data.frame();
for (m in 1:1){
  for (l in 1:1){
    for (k in 1:4){
      test<- read.table(paste("N1000_a",atable[m],"_r0_nL",ntable[k],"_nEq1_link",linktable[l],"_sP0.01_hP",htable[k],"_U0_L1_Haplo.txt",sep =""), sep = " ", fill = TRUE, header = FALSE )
      
      sim1 <-data.frame(pos = melt(as.list(test[1,2:(2*ntable[k]+1)]))[1], state= melt(as.list(test[3,2:(2*ntable[k]+1)]))[1] )
      colnames(sim1) = c("pos", "state")
      sim1 <- sim1 %>% mutate(freq = if_else(state == 0, 1, if_else(state == 2,1, 0)))
      sim1 <- sim1 %>% mutate(h = htable[k])
      datah = sim1;
      
      
      for(i in 1:99){
        sim1 <-data.frame(pos = melt(as.list(test[3*i+1,2:(2*ntable[k]+1)]))[1], state= melt(as.list(test[3*i+3,2:(2*ntable[k]+1)]))[1] ) 
        colnames(sim1) = c("pos", "state")
        sim1 <- sim1 %>% mutate(h = htable[k])
        sim1 <- sim1 %>% mutate(freq = if_else(state == 0, 1, if_else(state == 2,1, 0)))
        sim1 =sim1[order(sim1$pos),];
        datah = rbind(datah, sim1);
      }
      dHtot = rbind(datah, dHtot)
    }
  }
}

dfix_h<- ggplot(dHtot) +
  geom_smooth(mapping = aes(x=pos, y=freq,colour=as.factor(h)))+
  scale_y_continuous(limit=c(0,NA),oob=squish)+
  theme(text = element_text(size = 12, face = "bold"),legend.position="none",
        panel.background=element_blank(),axis.line = element_line(colour = "black"))+ theme(strip.background = element_blank(),strip.text.x = element_blank())+ 
  ylab(TeX(""))+
  xlab(TeX("Position along the genome"))+ 
  scale_color_manual(breaks = c("0.2", "0", "0.1", "0.3"), values=c("black","blue", "darkgreen", "darkgoldenrod1"))

dfix_h
grid.arrange(dfix_s, dfix_h, dHe_s, dHe_h, ncol = 2)

plot_grid(deltap_s, deltap_h, dfix_s, dfix_h, dHe_s, dHe_h, align = "v", nrow = 3, rel_heights = c(4/10, 3/10, 3/10))

grid.arrange(deltap_s, deltap_h, dfix_s, dfix_h, dHe_s, dHe_h , ncol = 2)

pdf("new_fig2.pdf",width=9,height=8)

print(
  grid.arrange(deltap_s, deltap_h, dfix_s, dfix_h, dHe_s, dHe_h, ncol = 2)
)

dev.off()

####################################################################################################################################
## figure 3 - unequal haplotypes and random positioning

# Fixation patterns
neqtable = c(0.8,1,1.2)
ntable = c(100)
linktable = c(1e-05)
atable = c(0)
dHtot = data.frame();
for (m in 1:1){
  for (l in 1:1){
    for (k in 1:3){
      test<- read.table(paste("N1000_a",atable[m],"_r0_nL",ntable[1],"_nEq",neqtable[k],"_link",linktable[l],"_sP0.01_hP0.2_U0_L1_Haplo.txt",sep =""), sep = " ", fill = TRUE, header = FALSE )
      
      nbcols = round(neqtable[k]*ntable[1]) +  ntable[1];
      sim1 <-data.frame(pos = melt(as.list(test[1,2:(nbcols+1)]))[1], state= melt(as.list(test[3,2:(nbcols+1)]))[1] )
      colnames(sim1) = c("pos", "state")
      sim1 <- sim1 %>% mutate(freq = if_else(state == 0, 1, if_else(state == 2,1, 0)))
      sim1 <- sim1 %>% mutate(neq = neqtable[k])
      sim1 <- sim1 %>% mutate(a = atable[m])
      datah = sim1;
      
      for(i in 1:99){
        sim1 <-data.frame(pos = melt(as.list(test[3*i+1,2:(nbcols+1)]))[1], state= melt(as.list(test[3*i+3,2:(nbcols+1)]))[1] ) 
        colnames(sim1) = c("pos", "state")
        sim1 <- sim1 %>% mutate(neq = neqtable[k])
        sim1 <- sim1 %>% mutate(a = atable[m])
        sim1 <- sim1 %>% mutate(freq = if_else(state == 0, 1, if_else(state == 2,1, 0)))
        sim1 =sim1[order(sim1$pos),];
        datah = rbind(datah, sim1);
      }
      
      dHtot = rbind(datah, dHtot)
    }
  }
}

neqtable = c(1,1.2)
atable = c(0.95)
for (m in 1:1){
  for (l in 1:1){
    for (k in 1:2){
      test<- read.table(paste("N1000_a",atable[m],"_r0_nL",ntable[1],"_nEq",neqtable[k],"_link",linktable[l],"_sP0.01_hP0.2_U0_L1_Haplo.txt",sep =""), sep = " ", fill = TRUE, header = FALSE )
      
      nbcols = round(neqtable[k]*ntable[1]) +  ntable[1];
      sim1 <-data.frame(pos = melt(as.list(test[1,2:(nbcols+1)]))[1], state= melt(as.list(test[3,2:(nbcols+1)]))[1] )
      colnames(sim1) = c("pos", "state")
      sim1 <- sim1 %>% mutate(freq = if_else(state == 0, 1, if_else(state == 2,1, 0)))
      sim1 <- sim1 %>% mutate(neq = neqtable[k])
      sim1 <- sim1 %>% mutate(a = atable[m])
      datah = sim1;
      
      for(i in 1:99){
        sim1 <-data.frame(pos = melt(as.list(test[3*i+1,2:(nbcols+1)]))[1], state= melt(as.list(test[3*i+3,2:(nbcols+1)]))[1] ) 
        colnames(sim1) = c("pos", "state")
        sim1 <- sim1 %>% mutate(neq = neqtable[k])
        sim1 <- sim1 %>% mutate(a = atable[m])
        sim1 <- sim1 %>% mutate(freq = if_else(state == 0, 1, if_else(state == 2,1, 0)))
        sim1 =sim1[order(sim1$pos),];
        datah = rbind(datah, sim1);
      }
      dHtot = rbind(datah, dHtot)
    }
  }
}

neqtable = c(0.8)
for (m in 1:1){
  for (l in 1:1){
    for (k in 1:1){
      test<- read.table(paste("N1000_a",atable[m],"_r0_nL",ntable[1],"_nEq",neqtable[k],"_link",linktable[l],"_sP0.01_hP0.2_U0_L1_Haplo.txt",sep =""), sep = " ", fill = TRUE, header = FALSE )
      
      nbcols = round(neqtable[k]*ntable[1]) +  ntable[1];
      sim1 <-data.frame(pos = melt(as.list(test[1,2:(nbcols+1)]))[1], state= melt(as.list(test[3,2:(nbcols+1)]))[1] )
      colnames(sim1) = c("pos", "state")
      sim1 <- sim1 %>% mutate(freq = if_else(state == 0, 1, if_else(state == 2,1, 0)))
      sim1 <- sim1 %>% mutate(neq = neqtable[k])
      sim1 <- sim1 %>% mutate(a = atable[m])
      datah = sim1;
      
      for(i in 1:99){
        sim1 <-data.frame(pos = melt(as.list(test[3*i+1,2:(nbcols+1)]))[1], state= melt(as.list(test[3*i+3,2:(nbcols+1)]))[1] ) 
        colnames(sim1) = c("pos", "state")
        sim1 <- sim1 %>% mutate(neq = neqtable[k])
        sim1 <- sim1 %>% mutate(a = atable[m])
        sim1 <- sim1 %>% mutate(freq = if_else(state == 0, 1, if_else(state == 2,1, 0)))
        sim1 =sim1[order(sim1$pos),];
        datah = rbind(datah, sim1);
      }
      
    }
  }
}

dfix_neq<- ggplot()+
  geom_smooth(data = dHtot,mapping =aes(x = pos, y = freq, colour=as.factor(neq), linetype = as.factor(a)))+
  scale_y_continuous(limit=c(0,NA),oob=squish)+
  stat_summary(data = datah,mapping =aes(x = pos, y = freq, colour=as.factor(neq), linetype = as.factor(a)), geom = "line", fun = "mean", size = 1)) + theme(text = element_text(size = 12, face = "bold"),legend.position="none",panel.background=element_blank(),axis.line = element_line(colour = "black"), axis.title = element_text(size = 14))+ theme(strip.background = element_blank(), axis.title.y = element_text(vjust=0.5), plot.title = element_text(hjust=0.5, size = 12, vjust = 0),legend.title = element_text(size = 9), legend.text = element_text(size = 8))+ 
  ylab(TeX("Proportion of loss/fixation"))+
  xlab(TeX("Position along the genome"))+
  labs(title = TeX("Equally spaced mutations"))+
  scale_color_manual(breaks = c("0.8" , "1","1.2"), values=c("blue","black", "darkgreen"))

dfix_neq

# Change in HE
neqtable = c(0.8,1,1.2)
ntable = c(100)
linktable = c(1e-05)
atable = c(0, 0.95)
dHtot = data.frame();
for (m in 1:2){
  for (l in 1:1){
    for (k in 1:3){
      test<- read.table(paste("N1000_a",atable[m],"_r0_nL",ntable[1],"_nEq",neqtable[k],"_link",linktable[l],"_sP0.01_hP0.2_U0_L1.txt",sep =""), sep = " ", fill = TRUE, header = FALSE )
      dcal <- ifelse(test[4:403,13] == 0, 0, test[4:403,12]/test[4:403,13])
      dH <- data.frame(gen = test[4:403,2], He = dcal, link = linktable[l], a = atable[m], neq = neqtable[k])
      for(i in 1:99){
        j = i*403 + 3;
        dcal <- ifelse(test[(j+1):(j+400),13] == 0, 0, test[(j+1):(j+400),12]/test[(j+1):(j+400),13])
        sim1 <-data.frame(gen = test[(j+1):(j+400),2], He = dcal, link = linktable[l], a = atable[m], neq = neqtable[k])
        dH = rbind(dH, sim1);
      }
      
      dHtot = rbind(dH, dHtot)
      
    }
  }
}

dHe_neq<- ggplot(data = dHtot)+
  geom_smooth(mapping =aes(x = gen, y = He, colour=as.factor(neq), linetype = as.factor(a)))+
  coord_cartesian(ylim = c(0, 0.52), xlim = c(0,4000)) +
  theme(text = element_text(size = 12, face = "bold"), legend.position=c(0.5,0.6), panel.background=element_blank(),axis.line = element_line(colour = "black"))+ theme(strip.background = element_blank(),strip.text.x = element_blank(),axis.title.y = element_text(vjust=0.5, angle = 0))+
  labs(color = TeX("Relative size of $2^{nd}$ haplotype"), lty = TeX("Selfing rate"))+ 
  ylab(TeX("$H_e$"))+
  xlab(TeX("Time (generations)"))+ 
  scale_color_manual(breaks = c( "0.8","1", "1.2"), values=c("blue", "black","darkgreen"))
dHe_neq


# Random positioning
# Fixation patterns

neqtable = c(0.8, 1, 1.2)
ntable = c(100)
linktable = c(1e-05)
atable = c(0)
dHtot = data.frame();
for (m in 1:1){
  for (l in 1:1){
    for (k in 1:3){
      test<- read.table(paste("N1000_a",atable[m],"_r1_nL",ntable[1],"_nEq",neqtable[k],"_link",linktable[l],"_sP0.01_hP0.2_U0_L1_Haplo.txt",sep =""), sep = " ", fill = TRUE, header = FALSE )
      
      nbcols = round(neqtable[k]*ntable[1]) +  ntable[1];
      sim1 <-data.frame(pos = melt(as.list(test[1,2:(nbcols+1)]))[1], state= melt(as.list(test[3,2:(nbcols+1)]))[1] )
      colnames(sim1) = c("pos", "state")
      sim1 <- sim1 %>% mutate(freq = if_else(state == 0, 1, if_else(state == 2,1, 0)))
      sim1 <- sim1 %>% mutate(neq = neqtable[k])
      sim1 <- sim1 %>% mutate(a = atable[m])
      datah = sim1;
      
      for(i in 1:99){
        sim1 <-data.frame(pos = melt(as.list(test[3*i+1,2:(nbcols+1)]))[1], state= melt(as.list(test[3*i+3,2:(nbcols+1)]))[1] ) 
        colnames(sim1) = c("pos", "state")
        sim1 <- sim1 %>% mutate(neq = neqtable[k])
        sim1 <- sim1 %>% mutate(a = atable[m])
        sim1 <- sim1 %>% mutate(freq = if_else(state == 0, 1, if_else(state == 2,1, 0)))
        sim1 =sim1[order(sim1$pos),];
        datah = rbind(datah, sim1);
      }
      
      dHtot = rbind(datah, dHtot)
    }
  }
}

neqtable = c(1,1.2)
atable = c(0.95)
for (m in 1:1){
  for (l in 1:1){
    for (k in 1:2){
      test<- read.table(paste("N1000_a",atable[m],"_r1_nL",ntable[1],"_nEq",neqtable[k],"_link",linktable[l],"_sP0.01_hP0.2_U0_L1_Haplo.txt",sep =""), sep = " ", fill = TRUE, header = FALSE )
      
      nbcols = round(neqtable[k]*ntable[1]) +  ntable[1];
      sim1 <-data.frame(pos = melt(as.list(test[1,2:(nbcols+1)]))[1], state= melt(as.list(test[3,2:(nbcols+1)]))[1] )
      colnames(sim1) = c("pos", "state")
      sim1 <- sim1 %>% mutate(freq = if_else(state == 0, 1, if_else(state == 2,1, 0)))
      sim1 <- sim1 %>% mutate(neq = neqtable[k])
      sim1 <- sim1 %>% mutate(a = atable[m])
      datah = sim1;
      
      for(i in 1:99){
        sim1 <-data.frame(pos = melt(as.list(test[3*i+1,2:(nbcols+1)]))[1], state= melt(as.list(test[3*i+3,2:(nbcols+1)]))[1] ) 
        colnames(sim1) = c("pos", "state")
        sim1 <- sim1 %>% mutate(neq = neqtable[k])
        sim1 <- sim1 %>% mutate(a = atable[m])
        sim1 <- sim1 %>% mutate(freq = if_else(state == 0, 1, if_else(state == 2,1, 0)))
        sim1 =sim1[order(sim1$pos),];
        datah = rbind(datah, sim1);
      }
      
      dHtot = rbind(datah, dHtot)
    }
  }
}

neqtable = c(0.8)
for (m in 1:1){
  for (l in 1:1){
    for (k in 1:1){
      test<- read.table(paste("N1000_a",atable[m],"_r1_nL",ntable[1],"_nEq",neqtable[k],"_link",linktable[l],"_sP0.01_hP0.2_U0_L1_Haplo.txt",sep =""), sep = " ", fill = TRUE, header = FALSE )
      
      nbcols = round(neqtable[k]*ntable[1]) +  ntable[1];
      sim1 <-data.frame(pos = melt(as.list(test[1,2:(nbcols+1)]))[1], state= melt(as.list(test[3,2:(nbcols+1)]))[1] )
      colnames(sim1) = c("pos", "state")
      sim1 <- sim1 %>% mutate(freq = if_else(state == 0, 1, if_else(state == 2,1, 0)))
      sim1 <- sim1 %>% mutate(neq = neqtable[k])
      sim1 <- sim1 %>% mutate(a = atable[m])
      datah = sim1;
      
      for(i in 1:99){
        sim1 <-data.frame(pos = melt(as.list(test[3*i+1,2:(nbcols+1)]))[1], state= melt(as.list(test[3*i+3,2:(nbcols+1)]))[1] ) 
        colnames(sim1) = c("pos", "state")
        sim1 <- sim1 %>% mutate(neq = neqtable[k])
        sim1 <- sim1 %>% mutate(a = atable[m])
        sim1 <- sim1 %>% mutate(freq = if_else(state == 0, 1, if_else(state == 2,1, 0)))
        sim1 =sim1[order(sim1$pos),];
        datah = rbind(datah, sim1);
      }
      
    }
  }
}


dfix_rand_neq<- ggplot()+
  geom_smooth(data = dHtot,mapping =aes(x = pos, y = freq, colour=as.factor(neq), linetype = as.factor(a)))+
  scale_y_continuous(limit=c(0,NA),oob=squish)+
  stat_summary(data = datah,mapping =aes(x = pos, y = freq, colour=as.factor(neq), linetype = as.factor(a)), geom = "line", fun = "mean", size = 1)) + theme(text = element_text(size = 12, face = "bold"),legend.position="none",panel.background=element_blank(),axis.line = element_line(colour = "black"), axis.title = element_text(size = 14))+ theme(strip.background = element_blank(), axis.title.y = element_text(vjust=0.5), plot.title = element_text(hjust=0.5, size = 12, vjust = 0),legend.title = element_text(size = 9), legend.text = element_text(size = 8))+ 
  ylab(TeX(""))+
  xlab(TeX("Position along the genome"))+
  labs(title = TeX("Randomly spaced mutations"))+
  scale_color_manual(breaks = c("0.8" , "1","1.2"), values=c("blue","black", "darkgreen"))

dfix_rand_neq

# Change in HE
neqtable = c(0.8,1,1.2)
ntable = c(100)
linktable = c(1e-05)
atable = c(0, 0.95)
dHtot = data.frame();
for (m in 1:2){
  for (l in 1:1){
    for (k in 1:3){
      test<- read.table(paste("N1000_a",atable[m],"_r1_nL",ntable[1],"_nEq",neqtable[k],"_link",linktable[l],"_sP0.01_hP0.2_U0_L1.txt",sep =""), sep = " ", fill = TRUE, header = FALSE )
      dcal <- ifelse(test[4:403,13] == 0, 0, test[4:403,12]/test[4:403,13])
      dH <- data.frame(gen = test[4:403,2], He = dcal, link = linktable[l], a = atable[m], neq = neqtable[k])
      for(i in 1:99){
        j = i*403 + 3;
        dcal <- ifelse(test[(j+1):(j+400),13] == 0, 0, test[(j+1):(j+400),12]/test[(j+1):(j+400),13])
        sim1 <-data.frame(gen = test[(j+1):(j+400),2], He = dcal, link = linktable[l], a = atable[m], neq = neqtable[k])
        dH = rbind(dH, sim1);
      }
      
      dHtot = rbind(dH, dHtot)
      
    }
  }
}


dHe_rand_neq<- ggplot(data = dHtot)+
  geom_smooth(mapping =aes(x = gen, y = He, colour=as.factor(neq), linetype = as.factor(a)))+
  coord_cartesian(ylim = c(0, 0.52), xlim = c(0,4000)) +
  theme(text = element_text(size = 12, face = "bold"), legend.position="non3", panel.background=element_blank(),axis.line = element_line(colour = "black"))+ theme(strip.background = element_blank(),strip.text.x = element_blank(),axis.title.y = element_text(vjust=0.5, angle = 0))+
  ylab(TeX(""))+
  xlab(TeX("Time (generations)"))+ 
  scale_color_manual(breaks = c( "0.8","1", "1.2"), values=c("blue", "black","darkgreen"))

dHe_rand_neq

pdf("new_fig3.pdf",width=9,height=7)
print(grid.arrange(dfix_neq, dfix_rand_neq, dHe_neq, dHe_rand_neq, ncol = 2))
dev.off()


###############################################################################################################################################

scaleFUN <- function(x) sprintf("%.2f", x)

# Figure 4 - Effect of Background mutations on POD Het

# relative He in POD with and without selection elsewhere in the genome  - He_U/He_U0
setwd("~/Desktop/pod_files/prog_final/h_0.2")
Utable = c(0.1,0.5)
Ltable = c(1, 10)
atable = c(0, 0.2, 0.4, 0.6, 0.8, 0.95)
stable = c(0.01,0)
htable = c(0.2,0)
dHtot = data.frame();
for (m in 1:6){
  for (l in 1:2){
    for (k in 1:2){
      for (o in 1:1){
        test<- read.table(paste("N1000_a",atable[m],"_r1_nL100_nEq1_link1e-06_sP",stable[1],"_hP",htable[1],"_U",Utable[k],"_L",Ltable[l],".txt",sep =""), sep = " ", fill = TRUE, header = FALSE )
        test2<- read.table(paste("N1000_a",atable[m],"_r1_nL100_nEq1_link1e-06_sP",stable[1],"_hP",htable[1],"_U0_L1.txt",sep =""), sep = " ", fill = TRUE, header = FALSE )
        dcal <- ifelse((test[403,13]) == 0, 0, (test[403,12])/(test[403,13]))
        dcal2 <- ifelse((test2[403,13]) == 0, 0, (test2[403,12])/(test2[403,13]))
        dH <- data.frame(He = mean(dcal), rHe = mean(dcal2), a = atable[m], U = Utable[k], L = Ltable[l])
        for(i in 1:99){
          j = i*403 + 3;
          dcal <- ifelse((test[(j+400),13]) == 0, 0, (test[(j+400),12])/(test[(j+400),13]))
          dcal2 <- ifelse((test2[(j+400),13]) == 0, 0, (test2[(j+400),12])/(test2[(j+400),13]))
          sim1 <-data.frame(He = mean(dcal), rHe = mean(dcal2), a = atable[m], U = Utable[k] , L = Ltable[l])
          dH = rbind(dH, sim1);
        }
        dnew <- data.frame(He = mean(dH$He), rHe = mean(dH$rHe), a = atable[m], U = Utable[k] , L = Ltable[l])
        dHtot = rbind(dnew, dHtot)
      }
    }
  }
}


dHepod_bgs_womut<- ggplot(data = dHtot)+
  geom_line(mapping =aes(x = a, y = He/rHe, colour=as.factor(U), linetype = as.factor(L)), size = 1.2)+ 
  geom_hline(yintercept=1, size=0.6, linetype="dashed", col = "black")+
  coord_cartesian(ylim = c(0.5, 4), xlim = c(0,1)) +
  theme(text = element_text(size = 12, face = "bold"),legend.position=c(0.1,0.7),panel.background=element_blank(),axis.line = element_line(colour = "black"), axis.title = element_text(size = 14))+ theme(strip.background = element_blank(), axis.title.y = element_text(vjust=0.5), legend.title = element_text(size = 9), legend.text = element_text(size = 8))+ scale_y_continuous(labels=scaleFUN)+
  ylab(TeX("Relative $H_e$ (POD zone)"))+
  xlab(TeX(""))+
  labs(color = TeX("$U"), lty = TeX("$L$"))+
  scale_color_manual(breaks = c("0","0.1", "0.5"), values=c("black","blue", "darkgreen"))

dHepod_bgs_womut


# relative He in rest of genome (w/o including pod region)  - He_pod/He_control
# different vlaues of hd (dominance of delterious mutaitons outside the POD zone) are saved in different folders
setwd("h_0.2")
Utable = c(0.1,0.5)
Ltable = c(1, 10)
atable = c(0, 0.2, 0.4, 0.6, 0.8, 0.95)
stable = c(0.01,0)
htable = c(0.2,0)
dHtot = data.frame();
for (m in 1:6){
  for (l in 1:2){
    for (k in 1:2){
      for (o in 1:1){
        test<- read.table(paste("N1000_a",atable[m],"_r1_nL100_nEq1_link1e-06_sP",stable[1],"_hP",htable[1],"_U",Utable[k],"_L",Ltable[l],".txt",sep =""), sep = " ", fill = TRUE, header = FALSE )
        test2<- read.table(paste("N1000_a",atable[m],"_r1_nL100_nEq1_link1e-06_sP",stable[2],"_hP",htable[2],"_U",Utable[k],"_L",Ltable[l],".txt",sep =""), sep = " ", fill = TRUE, header = FALSE )
        dcal <- ifelse((test[403,8]) == 0, 0, (test[403,7])/(test[403,8]))
        dcal2 <- ifelse((test2[403,8]) == 0, 0, (test2[403,7])/(test2[403,8]))
        dH <- data.frame( He = dcal, rHe = dcal2, a = atable[m], U = Utable[k], L = Ltable[l])
        for(i in 1:99){
          j = i*403 + 3;
          dcal <- ifelse((test[(j+400),8]) == 0, 0, (test[(j+400),7])/(test[(j+400),8]))
          dcal2 <- ifelse((test2[(j+400),8]) == 0, 0, (test2[(j+400),7])/(test2[(j+400),8]))
          sim1 <-data.frame(He = dcal, rHe = dcal2, a = atable[m], U = Utable[k] , L = Ltable[l])
          dH = rbind(dH, sim1);
        }
        dnew <- data.frame(He = mean(dH$He), rHe = mean(dH$rHe), a = atable[m], U = Utable[k] , L = Ltable[l])
        dHtot = rbind(dnew, dHtot)
        
      }
    }
  }
}


dHetot_bgs_wopod<- ggplot(data = dHtot)+
  geom_line(mapping =aes(x = a, y = He/rHe, colour=as.factor(U), linetype = as.factor(L)), size = 1.2)+ 
  geom_hline(yintercept=1, size=0.6, linetype="dashed", col = "black")+
  coord_cartesian(ylim = c(0.5, 2), xlim = c(0,1)) +
  theme(text = element_text(size = 12, face = "bold"),legend.position="none",panel.background=element_blank(),axis.line = element_line(colour = "black"), axis.title = element_text(size = 14))+ theme(strip.background = element_blank(), axis.title.y = element_text(vjust=0.5), legend.title = element_text(size = 9), legend.text = element_text(size = 8))+ 
  ylab(TeX("Relative $H_e$ \n (excluding POD zone)"))+
  xlab(TeX(""))+
  labs(color = TeX("$U"), lty = TeX("$L$"))+
  scale_y_continuous(labels=scaleFUN)+
  scale_color_manual(breaks = c("0","0.1", "0.5"), values=c("black","blue", "darkgreen"))

dHetot_bgs_wopod


# Effect of POD on Ne 

setwd("h_0.2")
Utable = c(0.1,0.5)
Ltable = c(1, 10)
atable = c(0, 0.2, 0.4, 0.6, 0.8, 0.95)
stable = c(0.01,0)
htable = c(0.2,0)
dHtot = data.frame();
for (m in 1:6){
  for (l in 1:2){
    for (k in 1:2){
      for (o in 1:1){
        test<- read.table(paste("N1000_a",atable[m],"_r1_nL100_nEq1_link1e-06_sP",stable[1],"_hP",htable[1],"_U",Utable[k],"_L",Ltable[l],".txt",sep =""), sep = " ", fill = TRUE, header = FALSE )
        test2<- read.table(paste("N1000_a",atable[m],"_r1_nL100_nEq1_link1e-06_sP",stable[2],"_hP",htable[2],"_U",Utable[k],"_L",Ltable[l],".txt",sep =""), sep = " ", fill = TRUE, header = FALSE )
        test$ne<- (1-test[,14])/(4*0.001*(test[,14]))
        test2$ne<- (1-test2[,14])/(4*0.001*(test2[,14]))
        dcal <- test$ne[403]
        dcal <- test2$ne[403]
        dH <- data.frame(Ne = dcal, rNe = dcal2, a = atable[m], U = Utable[k], L = Ltable[l])
        for(i in 1:99){
          j = i*403 + 3;
          dcal <- test$ne[(j+400)]
          dcal2 <- test2$ne[(j+400)]
          sim1 <-data.frame(Ne = dcal, rNe = dcal2, a = atable[m], U = Utable[k] , L = Ltable[l])
          dH = rbind(dH, sim1);
        }
        dnew <- data.frame(Ne = mean(dH$Ne), rNe = mean(dH$rNe), a = atable[m], U = Utable[k] , L = Ltable[l])
        dHtot = rbind(dnew, dHtot)
      }
    }
  }
}

Ne_bgs_wopod<- ggplot(data = dHtot)+
  geom_line(mapping =aes(x = a, y = Ne/rNe, colour=as.factor(U), linetype = as.factor(L)), size = 1.2)+ 
  geom_hline(yintercept=1, size=0.6, linetype="dashed", col = "black")+
  coord_cartesian(ylim = c(0.5, 1.5), xlim = c(0,1)) +
  theme(text = element_text(size = 12, face = "bold"),legend.position="none",panel.background=element_blank(),axis.line = element_line(colour = "black"), axis.title = element_text(size = 14))+ theme(strip.background = element_blank(), axis.title.y = element_text(vjust=0.5), legend.title = element_text(size = 9), legend.text = element_text(size = 8))+ 
  ylab(TeX("Relative $N_e$"))+
  xlab(TeX("Selfing rate $\\sigma$"))+
  labs(color = TeX("$U"), lty = TeX("$L$"))+
  scale_y_continuous(labels=scaleFUN)+
  scale_color_manual(breaks = c("0","0.1", "0.5"), values=c("black","blue", "darkgreen"))

Ne_bgs_wopod

grid.arrange(dHepod_bgs_womut,dHetot_bgs_wopod,Ne_bgs_wopod, ncol=1)

pdf("new_fig4.pdf",width=7,height=9)
print(grid.arrange(dHepod_bgs_womut,dHetot_bgs_wopod,Ne_bgs_wopod, ncol=1))
dev.off()


## Sup 4 BGS vs AOD

# relative He in POD with BGS and AOD  - He_U/He_U0

setwd("/h_0.2")
Utable = c(0.5)
Ltable = c(10,1)
atable = c(0, 0.95)
stable = c(0.01,0)
htable = c(0.2,0)
dHtot = data.frame();
for (m in 1:2){
  for (l in 1:2){
    for (k in 1:1){
      for (o in 1:1){
        test<- read.table(paste("N1000_a",atable[m],"_r1_nL100_nEq1_link1e-06_sP",stable[1],"_hP",htable[1],"_U",Utable[k],"_L",Ltable[l],".txt",sep =""), sep = " ", fill = TRUE, header = FALSE )
        test2<- read.table(paste("N1000_a",atable[m],"_r1_nL100_nEq1_link1e-06_sP",stable[1],"_hP",htable[1],"_U0_L1.txt",sep =""), sep = " ", fill = TRUE, header = FALSE )
        dcal <- ifelse((test[403,13]) == 0, 0, (test[403,12])/(test[403,13]))
        dcal2 <- ifelse((test2[403,13]) == 0, 0, (test2[403,12])/(test2[403,13]))
        dH <- data.frame(He = dcal, rHe = dcal2, a = atable[m], U = Utable[k], L = Ltable[l])
        for(i in 1:99){
          j = i*403 + 3;
          dcal <- ifelse((test[(j+400),13]) == 0, 0, (test[(j+400),12])/(test[(j+400),13]))
          dcal2 <- ifelse((test2[(j+400),13]) == 0, 0, (test2[(j+400),12])/(test2[(j+400),13]))
          sim1 <-data.frame(He = dcal, rHe = dcal2, a = atable[m], U = Utable[k] , L = Ltable[l])
          dH = rbind(dH, sim1);
        }
        dnew <- data.frame(He = mean(dH$He), rHe = mean(dH$rHe), a = atable[m], U = Utable[k] , L = Ltable[l], hval = 0.2)
        dHtot = rbind(dnew, dHtot)
      }
    }
  }
}


setwd("/h_0.5")
Utable = c(0.5)
Ltable = c(1, 10)
atable = c(0, 0.95)
stable = c(0.01,0)
htable = c(0.2,0)
for (m in 1:2){
  for (l in 1:2){
    for (k in 1:1){
      for (o in 1:1){
        test<- read.table(paste("N1000_a",atable[m],"_r1_nL100_nEq1_link1e-06_sP",stable[1],"_hP",htable[1],"_U",Utable[k],"_L",Ltable[l],".txt",sep =""), sep = " ", fill = TRUE, header = FALSE )
        test2<- read.table(paste("~/Desktop/pod_files/prog_final/h_0.2/N1000_a",atable[m],"_r1_nL100_nEq1_link1e-06_sP",stable[1],"_hP",htable[1],"_U0_L1.txt",sep =""), sep = " ", fill = TRUE, header = FALSE )
        dcal <- ifelse((test[403,13]) == 0, 0, (test[403,12])/(test[403,13]))
        dcal2 <- ifelse((test2[403,13]) == 0, 0, (test2[403,12])/(test2[403,13]))
        dH <- data.frame(He = dcal, rHe = dcal2, a = atable[m], U = Utable[k], L = Ltable[l])
        for(i in 1:99){
          j = i*403 + 3;
          dcal <- ifelse((test[(j+400),13]) == 0, 0, (test[(j+400),12])/(test[(j+400),13]))
          dcal2 <- ifelse((test2[(j+400),13]) == 0, 0, (test2[(j+400),12])/(test2[(j+400),13]))
          sim1 <-data.frame(He = dcal, rHe = dcal2, a = atable[m], U = Utable[k] , L = Ltable[l])
          dH = rbind(dH, sim1);
        }
        dnew <- data.frame(He = mean(dH$He), rHe = mean(dH$rHe), a = atable[m], U = Utable[k] , L = Ltable[l], hval = 0.5)
        dHtot = rbind(dnew, dHtot)
      }
    }
  }
}

dHe_bgs_aod<- ggplot(data = dHtot)+
  geom_point(mapping =aes(x = factor(a), y = He/rHe, colour=as.factor(hval), shape = as.factor(L)), size = 3)+
  coord_cartesian(ylim = c(0, 4)) +
  theme(text = element_text(size = 12, face = "bold"),legend.position=c(0.2,0.8), legend.direction = "horizontal", panel.background=element_blank(),axis.line = element_line(colour = "black"))+ theme(strip.background = element_blank(),strip.text.x = element_blank())+ 
  ylab(TeX("Relative $H_e$ (POD zone)"))+
  xlab(TeX(""))+
  labs( shape = TeX("$L$"),color = TeX("$h_d$"))+
  scale_color_manual(breaks = c("0","0.2", "0.5"), values=c("black","blue", "darkgreen"))+
  geom_hline(yintercept=1, size=0.6, linetype="dashed", col = "black")
dHe_bgs_aod

# relative He in rest of genome (w/o including pod region)  - He_pod/He_control
setwd("~/Desktop/pod_files/prog_final/h_0.2")
Utable = c(0.5)
Ltable = c(1, 10)
atable = c(0, 0.95)
stable = c(0.01,0)
htable = c(0.2,0)
dHtot = data.frame();
for (m in 1:2){
  for (l in 1:2){
    for (k in 1:1){
      for (o in 1:1){
        test<- read.table(paste("N1000_a",atable[m],"_r1_nL100_nEq1_link1e-06_sP",stable[1],"_hP",htable[1],"_U",Utable[k],"_L",Ltable[l],".txt",sep =""), sep = " ", fill = TRUE, header = FALSE )
        test2<- read.table(paste("N1000_a",atable[m],"_r1_nL100_nEq1_link1e-06_sP",stable[2],"_hP",htable[2],"_U",Utable[k],"_L",Ltable[l],".txt",sep =""), sep = " ", fill = TRUE, header = FALSE )
        dcal <- ifelse((test[403,8]) == 0, 0, (test[403,7])/(test[403,8]))
        dcal2 <- ifelse((test2[403,8]) == 0, 0, (test2[403,7])/(test2[403,8]))
        dH <- data.frame( He = dcal, rHe = dcal2, a = atable[m], U = Utable[k], L = Ltable[l])
        for(i in 1:99){
          j = i*403 + 3;
          dcal <- ifelse((test[(j+400),8]) == 0, 0, (test[(j+400),7])/(test[(j+400),8]))
          dcal2 <- ifelse((test2[(j+400),8]) == 0, 0, (test2[(j+400),7])/(test2[(j+400),8]))
          sim1 <-data.frame(He = dcal, rHe = dcal2, a = atable[m], U = Utable[k] , L = Ltable[l])
          dH = rbind(dH, sim1);
        }
        dnew <- data.frame(He = mean(dH$He), rHe = mean(dH$rHe), a = atable[m], U = Utable[k] , L = Ltable[l], hval = 0.2)
        dHtot = rbind(dnew, dHtot)
        
      }
    }
  }
}

setwd("~/Desktop/pod_files/prog_final/h_0.5")
Utable = c(0.5)
Ltable = c(1, 10)
atable = c(0, 0.95)
stable = c(0.01,0)
htable = c(0.2,0)
for (m in 1:2){
  for (l in 1:2){
    for (k in 1:1){
      for (o in 1:1){
        test<- read.table(paste("N1000_a",atable[m],"_r1_nL100_nEq1_link1e-06_sP",stable[1],"_hP",htable[1],"_U",Utable[k],"_L",Ltable[l],".txt",sep =""), sep = " ", fill = TRUE, header = FALSE )
        test2<- read.table(paste("N1000_a",atable[m],"_r1_nL100_nEq1_link1e-06_sP",stable[2],"_hP",htable[2],"_U",Utable[k],"_L",Ltable[l],".txt",sep =""), sep = " ", fill = TRUE, header = FALSE )
        dcal <- ifelse((test[403,8]) == 0, 0, (test[403,7])/(test[403,8]))
        dcal2 <- ifelse((test2[403,8]) == 0, 0, (test2[403,7])/(test2[403,8]))
        dH <- data.frame( He = dcal, rHe = dcal2, a = atable[m], U = Utable[k], L = Ltable[l])
        for(i in 1:99){
          j = i*403 + 3;
          dcal <- ifelse((test[(j+400),8]) == 0, 0, (test[(j+400),7])/(test[(j+400),8]))
          dcal2 <- ifelse((test2[(j+400),8]) == 0, 0, (test2[(j+400),7])/(test2[(j+400),8]))
          sim1 <-data.frame(He = dcal, rHe = dcal2, a = atable[m], U = Utable[k] , L = Ltable[l])
          dH = rbind(dH, sim1);
        }
        dnew <- data.frame(He = mean(dH$He), rHe = mean(dH$rHe), a = atable[m], U = Utable[k] , L = Ltable[l], hval = 0.5)
        dHtot = rbind(dnew, dHtot)
        
      }
    }
  }
}

dHe_bgs_aod_wopod <- ggplot(data = dHtot)+
  geom_point(mapping =aes(x = factor(a), y = He/rHe, colour=as.factor(hval), shape = as.factor(L)), size = 3)+
  coord_cartesian(ylim = c(0, 2)) +
  theme(text = element_text(size = 12, face = "bold"),legend.position="none", legend.direction = "horizontal", panel.background=element_blank(),axis.line = element_line(colour = "black"))+ theme(strip.background = element_blank(),strip.text.x = element_blank())+ 
  ylab(TeX("Relative $H_e$ (excluding POD zone)"))+
  xlab(TeX(""))+
  labs( shape = TeX("$L$"),color = TeX("$h_d$"))+
  scale_color_manual(breaks = c("0","0.2", "0.5"), values=c("black","blue", "darkgreen"))+
  geom_hline(yintercept=1, size=0.6, linetype="dashed", col = "black")

dHe_bgs_aod_wopod

# Effect of POD on Ne 

setwd("/h_0.2")
Utable = c(0.5)
Ltable = c(1, 10)
atable = c(0, 0.95)
stable = c(0.01,0)
htable = c(0.2,0)
dHtot = data.frame();
for (m in 1:2){
  for (l in 1:2){
    for (k in 1:1){
      for (o in 1:1){
        test<- read.table(paste("N1000_a",atable[m],"_r1_nL100_nEq1_link1e-06_sP",stable[1],"_hP",htable[1],"_U",Utable[k],"_L",Ltable[l],".txt",sep =""), sep = " ", fill = TRUE, header = FALSE )
        test2<- read.table(paste("N1000_a",atable[m],"_r1_nL100_nEq1_link1e-06_sP",stable[2],"_hP",htable[2],"_U",Utable[k],"_L",Ltable[l],".txt",sep =""), sep = " ", fill = TRUE, header = FALSE )
        dcal <- (1-test[403,14])/(4*0.001*test[403,14])
        dcal2 <- (1-test2[403,14])/(4*0.001*test2[403,14])
        dH <- data.frame( Ne = dcal, rNe = dcal2, a = atable[m], U = Utable[k], L = Ltable[l])
        for(i in 1:99){
          j = i*403 + 3;
          dcal <- (1-test[(j+400),14])/(4*0.001*test[(j+400),14])
          dcal2 <- (1-test2[(j+400),14])/(4*0.001*test2[(j+400),14])
          sim1 <-data.frame(Ne = dcal, rNe = dcal2, a = atable[m], U = Utable[k] , L = Ltable[l])
          dH = rbind(dH, sim1);
        }
        dnew <- data.frame(Ne = mean(dH$Ne), rNe = mean(dH$rNe), a = atable[m], U = Utable[k] , L = Ltable[l], hval = 0.2)
        dHtot = rbind(dnew, dHtot)
      }
    }
  }
}

setwd("/h_0.5")
Utable = c(0.5)
Ltable = c(1, 10)
atable = c(0, 0.95)
stable = c(0.01,0)
htable = c(0.2,0)
for (m in 1:2){
  for (l in 1:2){
    for (k in 1:1){
      for (o in 1:1){
        test<- read.table(paste("N1000_a",atable[m],"_r1_nL100_nEq1_link1e-06_sP",stable[1],"_hP",htable[1],"_U",Utable[k],"_L",Ltable[l],".txt",sep =""), sep = " ", fill = TRUE, header = FALSE )
        test2<- read.table(paste("~/Desktop/pod_files/prog_final/h_0.2/N1000_a",atable[m],"_r1_nL100_nEq1_link1e-06_sP",stable[2],"_hP",htable[2],"_U",Utable[k],"_L",Ltable[l],".txt",sep =""), sep = " ", fill = TRUE, header = FALSE )
        dcal <- (1-test[403,14])/(4*0.001*test[403,14])
        dcal2 <- (1-test2[403,14])/(4*0.001*test2[403,14])
        dH <- data.frame( Ne = dcal, rNe = dcal2, a = atable[m], U = Utable[k], L = Ltable[l])
        for(i in 1:99){
          j = i*403 + 3;
          dcal <- (1-test[(j+400),14])/(4*0.001*test[(j+400),14])
          dcal2 <- (1-test2[(j+400),14])/(4*0.001*test2[(j+400),14])
          sim1 <-data.frame(Ne = dcal, rNe = dcal2, a = atable[m], U = Utable[k] , L = Ltable[l])
          dH = rbind(dH, sim1);
        }
        dnew <- data.frame(Ne = mean(dH$Ne), rNe = mean(dH$rNe), a = atable[m], U = Utable[k] , L = Ltable[l], hval = 0.5)
        dHtot = rbind(dnew, dHtot)
      }
    }
  }
}

Ne_bgs_aod_wopod<- ggplot(data = dHtot)+
  geom_point(mapping =aes(x = factor(a), y = Ne/rNe, colour=as.factor(hval), shape = as.factor(L)), size = 3)+ 
  geom_hline(yintercept=1, size=0.6, linetype="dashed", col = "black")+
  coord_cartesian(ylim = c(0, 1.5)) +
  theme(text = element_text(size = 12, face = "bold"),legend.position="none",panel.background=element_blank(),axis.line = element_line(colour = "black"), axis.title = element_text(size = 14))+ theme(strip.background = element_blank(), axis.title.y = element_text(vjust=0.5), legend.title = element_text(size = 9), legend.text = element_text(size = 8))+ 
  ylab(TeX("Relative $N_e$"))+
  xlab(TeX("Selfing rate $\\sigma$"))+
  scale_color_manual(breaks = c("0","0.2", "0.5"), values=c("black","blue", "darkgreen"))

Ne_bgs_aod_wopod

grid.arrange(dHe_bgs_aod,dHe_bgs_aod_wopod,Ne_bgs_aod_wopod, ncol=1)

pdf("new_fig4_supp.pdf",width=7,height=9)
grid.arrange(dHe_bgs_aod,dHe_bgs_aod_wopod,Ne_bgs_aod_wopod, ncol=1)

dev.off()

########################################################################################################################################

# relative ID (w/o including pod region)  - He_pod/He_control
setwd("/h_0.2")
Utable = c(0.1,0.5)
Ltable = c(1, 10)
atable = c(0, 0.2, 0.4, 0.6, 0.8, 0.95)
stable = c(0.01,0)
htable = c(0.2,0)
dHtot = data.frame();
for (m in 1:6){
  for (l in 1:2){
    for (k in 1:2){
      for (o in 1:1){
        test<- read.table(paste("N1000_a",atable[m],"_r1_nL100_nEq1_link1e-06_sP",stable[1],"_hP",htable[1],"_U",Utable[k],"_L",Ltable[l],".txt",sep =""), sep = " ", fill = TRUE, header = FALSE )
        test2<- read.table(paste("N1000_a",atable[m],"_r1_nL100_nEq1_link1e-06_sP",stable[2],"_hP",htable[2],"_U",Utable[k],"_L",Ltable[l],".txt",sep =""), sep = " ", fill = TRUE, header = FALSE )
        dcal <- test[403,4]
        dcal2 <- test2[403,4]
        dH <- data.frame( He = dcal, rHe = dcal2, a = atable[m], U = Utable[k], L = Ltable[l])
        for(i in 1:99){
          j = i*403 + 3;
          dcal <- test[(j+400),4]
          dcal2 <- test2[(j+400),4]
          sim1 <-data.frame(He = dcal, rHe = dcal2, a = atable[m], U = Utable[k] , L = Ltable[l])
          dH = rbind(dH, sim1);
        }
        dnew <- data.frame(He = mean(dH$He), rHe = mean(dH$rHe), a = atable[m], U = Utable[k] , L = Ltable[l])
        dHtot = rbind(dnew, dHtot)
        
      }
    }
  }
}


ID<- ggplot(data = dHtot)+
  geom_line(mapping =aes(x = a, y = He/rHe, colour=as.factor(U), linetype = as.factor(L)), size = 1.2)+
  coord_cartesian(ylim = c(0, 4), xlim = c(0,1)) +
  theme(text = element_text(size = 12, face = "bold"),legend.position=c(0.1,0.8), legend.direction ="horizontal",panel.background=element_blank(),axis.line = element_line(colour = "black"), axis.title = element_text(size = 14))+ theme(strip.background = element_blank(), axis.title.y = element_text(vjust=0.5), legend.title = element_text(size = 9), legend.text = element_text(size = 8))+
  ylab(TeX("Relative $\\delta$"))+
  xlab(TeX("Selfing rate $\\sigma$"))+
  labs(color = TeX("U"), lty = TeX("$L$"))+
  scale_color_manual(breaks = c("0","0.1", "0.5"), values=c("black","blue", "darkgreen"))+
  geom_hline(yintercept=1, size=0.6, linetype="dashed", col = "black")

ID

pdf("new_fig5_supp_rel.pdf",width=7,height=3)
ID
dev.off()


df_OD <- data.frame("a" = seq(0,1,0.05), "id"=c(0.146716, 0.149368, 0.15215, 0.155073, 0.158152, 0.161402, 0.164844, 0.1685, 0.172397, 0.176568, 0.181056, 0.185912, 0.191203, 0.197019, 0.203481, 0.210765, 0.219143, 0.229064, 0.241401, 0.258339, 0.293431),"s" = as.factor(3), U =0);
df_U01 <- data.frame("a" = seq(0,1,0.05), "id"=c(0.251727, 0.249979, 0.24862, 0.247636, 0.247021, 0.246768, 0.246878, 0.247356, 0.248211, 0.249462, 0.251134, 0.253262, 0.255897, 0.259111, 0.263004, 0.267725, 0.27351, 0.280752, 0.290219, 0.30384, 0.333584),"s" = as.factor(3), U = 0.1);
df_U05 <- data.frame("a" = seq(0,1,0.05), "id"=c(0.557488, 0.546685, 0.536529, 0.526999, 0.518075, 0.509737, 0.501969, 0.494758, 0.488094, 0.481972, 0.476393, 0.471364, 0.466902, 0.463036, 0.459816, 0.457318, 0.45567, 0.455095, 0.456031, 0.459589, 0.472636),"s" = as.factor(3), U = 0.5);
# 
df_U01s <- data.frame("a" = seq(0,1,0.05), "id"=c(0.123067, 0.116425, 0.110432, 0.105005, 0.100071, 0.0955678, 0.0914436, 0.0876539, 0.0841604, 0.0809306, 0.0779362, 0.0751527, 0.0725591, 0.0701367, 0.0678693, 0.0657428, 0.0637444, 0.061863, 0.0600888, 0.0584129, 0.0568274),"s" = as.factor(4), U = 0.1);
df_U05s <- data.frame("a" = seq(0,1,0.05), "id"=c(0.481401, 0.46146, 0.442949, 0.425748, 0.409743, 0.394826, 0.380902,0.367882, 0.355687, 0.344245, 0.333493, 0.323372, 0.313831, 0.304823,0.296306, 0.288242, 0.280597, 0.27334, 0.266443, 0.259879, 0.253627),"s" = as.factor(4), U = 0.5);

setwd("/h_0.2")
Utable = c(0.1,0.5)
Ltable = c(1)
atable = c(0, 0.2, 0.4, 0.6, 0.8, 0.95)
stable = c(0.01,0)
htable = c(0.2,0)
dHtot = data.frame();
dnew <- c()
for (m in 1:6){
  for (l in 1:1){
    for (k in 1:2){
      for (o in 1:1){
        test<- read.table(paste("N1000_a",atable[m],"_r1_nL100_nEq1_link1e-06_sP",stable[1],"_hP",htable[1],"_U",Utable[k],"_L",Ltable[l],".txt",sep =""), sep = " ", fill = TRUE, header = FALSE )
        dcal <- test[403,4]
        dH <- data.frame( He = dcal, a = atable[m], U = Utable[k], L = Ltable[l])
        for(i in 1:99){
          j = i*403 + 3;
          dcal <- test[(j+400),4]
          sim1 <-data.frame(He = dcal, a = atable[m], U = Utable[k] , L = Ltable[l])
          dH = rbind(dH, sim1);
        }
        dHtot = rbind(dH, dHtot)
        d <- data.frame(He = mean(dH$He),  a = atable[m], U = Utable[k] , L = Ltable[l], s = stable[o])
        dnew = rbind(dnew, d)
        
      }
    }
  }
}


IDnorm1<- ggplot(data = dHtot)+
  geom_line(data = dnew, mapping =aes(x = a, y = He, colour=as.factor(U)), size = 1)+
  geom_point(mapping =aes(x = a, y = He, colour=as.factor(U)), alpha = 0.1)+
  coord_cartesian(ylim = c(0, 0.6), xlim = c(0,1))  +
  theme(text = element_text(size = 12, face = "bold"),legend.position=c(0.2,0.1), legend.direction ="horizontal",panel.background=element_blank(),axis.line = element_line(colour = "black"), axis.title = element_text(size = 14))+ theme(strip.background = element_blank(), axis.title.y = element_text(vjust=0.5), legend.title = element_text(size = 9), legend.text = element_text(size = 8))+
  ylab(TeX("Inbreeding depression $\\delta$"))+
  xlab(TeX("Selfing rate $\\sigma$"))+
  labs(color = TeX("U"))+
  scale_color_manual(breaks = c("0","0.1", "0.5"), values=c("black","blue", "darkgreen"))+ #geom_line(data = df_OD, mapping =aes(x = a, y = id),color = "black",linetype=2, size = 1 )+ 
  geom_line(data = df_U01, mapping =aes(x = a, y = id),color = "blue",linetype=2, size = 1 )+ 
  geom_line(data = df_U05, mapping =aes(x = a, y = id),color = "darkgreen",linetype=2, size = 1 )+
  geom_line(data = df_U01s, mapping =aes(x = a, y = id),color = "blue",linetype=3, size = 1 )+ 
  geom_line(data = df_U05s, mapping =aes(x = a, y = id),color = "darkgreen",linetype=3, size = 1 )

IDnorm1

pdf("new_fig5_supp_L1.pdf",width=7,height=3)
IDnorm1
dev.off()


p1<-ggplot(data = dHtot)+
  geom_violin(mapping = aes(factor(a), He, colour=as.factor(U)))+
  coord_cartesian(ylim = c(0, 0.6))  +
  theme(text = element_text(size = 12, face = "bold"),legend.position=c(0.2,0.1), legend.direction ="horizontal",panel.background=element_blank(),axis.line = element_line(colour = "black"), axis.title = element_text(size = 14))+ theme(strip.background = element_blank(), axis.title.y = element_text(vjust=0.5), legend.title = element_text(size = 9), legend.text = element_text(size = 8))+
  ylab(TeX("Inbreeding depression $\\delta$"))+
  xlab(TeX("Selfing rate $\\sigma$"))+
  labs(color = TeX("U"))+
  scale_color_manual(breaks = c("0","0.1", "0.5"), values=c("black","blue", "darkgreen"))

p1

pdf("new_fig5_supp_vio.pdf",width=7,height=3)
p1
dev.off()
```
